# Driving Forces of RNA Condensation Revealed through Coarse-Grained Modeling with Explicit Mg^2+^

**DOI:** 10.1101/2024.11.17.624048

**Authors:** Shanlong Li, Jianhan Chen

## Abstract

RNAs are major drivers of phase separation in the formation of biomolecular condensates, and can undergo protein-free phase separation in the presence of divalent ions or crowding agents. Much remains to be understood regarding how the complex interplay of base stacking, base pairing, electrostatics, ion interactions, and particularly structural propensities governs RNA phase behavior. Here we develop an **i**ntermediate resolution model for **<u>con</u>**densates of **<u>RNA</u>**s (iConRNA) that can capture key local and long-range structure features of dynamic RNAs and simulate their spontaneous phase transitions with Mg^2+^. Representing each nucleotide using 6-7 beads, iConRNA accurately captures base stacking and pairing and includes explicit Mg^2+^. The model does not only reproduce major conformational properties of poly(rA) and poly(rU), but also correctly folds small structured RNAs and predicts their melting temperatures. With an effective model of explicit Mg^2+^, iConRNA successfully recapitulates experimentally observed lower critical solution temperature phase separation of poly(rA) and triplet repeats, and critically, the nontrivial dependence of phase transitions on RNA sequence, length, concentration, and Mg^2+^ level. Further mechanistic analysis reveals a key role of RNA folding in modulating phase separation as well as its temperature and ion dependence, besides other driving forces such as Mg^2+^-phosphate interactions, base stacking, and base pairing. These studies also support iConRNA as a powerful tool for direct simulation of RNA-driven phase transitions, enabling molecular studies of how RNA conformational dynamics and its response to complex condensate environment control the phase behavior and condensate material properties.

**SIGNIFICANCE STATEMENT:** Dynamic RNAs and proteins are major drivers of biomolecular phase separation that has been recently discovered to underlie numerous biological processes and be involved in many human diseases. Molecular simulation has an indispensable role to play in dissecting the driving forces and regulation of biomolecular phase separation. The current work describes a high-resolution coarse-grained RNA model that is capable of describing the structure dynamics and complex sequence, concentration, temperature and ion dependent phase transitions of flexible RNAs. The study further reveals a central role of RNA folding in coordinating Mg^2+^-phosphate interactions, base stacking, and base pairing to drive phase separation, paving the road for studies of RNA-mediated phase separation in relevant biological contexts.

## INTRODUCTION

Biomolecular condensates have attracted intensive interest in recent years and are believed to play important roles in myriad cellular functions ranging from RNA storage and processing, stress responses, metabolism, and immune response, to cellular signaling [1–5]. They have also been implicated in a variety of human diseases including cancers and neurodegenerative diseases [6–11]. Formation of condensates involves spontaneous phase separations, mediated by dynamic and multivalent biological macromolecules such as proteins and nucleic acids [12–14]. Interestingly, even though the first biomolecular condensate discovered was germline P granules containing RNA and RNA-binding proteins [1], phase separation of proteins has drawn more intensive research interests in the field. Extensive efforts from experiment, theory, and simulation have provided many insights into the molecular driving forces and sequence-specific phase behaviors as well as the thermodynamic, dynamic, and material properties of the condensates [15–25]. In contrast, less attention has been paid to RNA-mediated phase separation [26].

The importance of RNAs in biomolecular condensates such as Ribonucleoprotein (RNP) granules is well recognized [27]. On one hand, RNAs can drive the formation and stability of phase-separated condensates. RNA binding proteins (RBPs) interact with RNAs through both specific bindings, medicated by well-structured RNA binding domains such as RNA recognition motifs (RRMs) [28–30], and nonspecific interactions, through positively charged arginine-glycine-glycine (RGG) domains [31–33]. These interactions can promote the phase separation through seeding and accelerating condensation of RBPs [34–37], where RNA, especially long noncoding RNA (lncRNA), serves as a scaffold to bind multi-RBPs or directly participates in the phase separation. On the other hand, RNAs have potent effects on controlling material properties of phase-separated droplets, such as viscosity and fluidity [30,31,38,39]. For example, the addition of RNA can increase the observed fluidity of LAF-1 condensates [31]. Importantly, recent experiments have reported that RNA can undergo protein-free phase separation in the presence of multivalent cations or crowding agents [39–45]. In the presence of Mg^2+^ ions, RNA homopolymers, including poly(adenylic acid) (poly(rA)), poly(uridylic acid) (poly(rU)), and poly(cytidylic acid) (poly(rC)), display thermo-responsive phase separations with lower critical solution temperature (LCST) [44]. RNA triplet repeats, like CAG or CUG repeats, are able to lead to their coalescence into nuclear foci [40], which is involved in several neurological and neuromuscular disorders [46–49]. These RNA condensates show strong [Mg^2+^], temperature, sequence, and length dependence, giving rise to fascinating complexity of RNA phase transitions. At present, much remains to be understood regarding the molecular mechanisms and driving forces of RNA-mediated phase separation. In particular, there is a critical need to dissect how the interplay of base stacking, base pairing, electrostatics, ion interactions, and dynamic structural propensities governs the phase behaviours of RNAs.

Molecular dynamics (MD) simulations have played an important role in dissecting the molecular mechanisms and sequence determinant of protein phase separation [50,51], particularly with various coarse-grained (CG) models designed to reach the timescales and length scales required for simulation of condensates [52–56]. There have also been numerous CG models of RNAs, which have been summarized in a comprehensive recent review [57]. Notably, most existing CG RNA models have been designed for the modeling and prediction of 2D and 3D folded structures and thermodynamics, with different levels of resolutions and consideration of key interactions such as stacking, pairing, and various electrostatic interactions [58–64]. At present, only RNA models with one bead per nucleotide have been designed and applied to study phase separation, including the HPS-based model, Mpipi, and COCOMO [65–68]. These models primarily focus on supplementing existing protein models for simulating heterotypic RNA-protein phase separation. The pioneering work for simulating homotypic RNA phase separation was based on the single interaction site (SIS) model developed by the Thirumalai group [69]. SIS simulations were able to reproduce key experimental findings in the phase separation of RNA triplet repeats [40] and provided a qualitative description of the morphologies and dynamics of RNA chains in condensates. However, the single-bead models do not provide sufficient spatial resolution to describe and resolve the complexity of dynamic RNA structure and interactions, severely limiting their utility in studying RNA phase separation.

Here, we developed a new **i**ntermediate resolution model for **<u>con</u>**densates of **<u>RNA</u>**s (iConRNA) that represents each nucleotide using 6 or 7 beads and explicitly considers a range of backbone and base-mediated interactions. The model is carefully balanced to capture a range of local structures and long-range dynamics of flexible RNAs and thermodynamic properties of folded RNAs. Furthermore, iConRNA includes explicit Mg^2+^ ions and provides a protocol to calibrate the temperature and concentration dependence of Mg^2+^/phosphate interactions based on experiment, simulation, and/or theory. The final model does not only capture the salt and Mg^2+^ dependence of model flexible RNAs but also successfully folds small RNAs. We further demonstrate that iConRNA is suitable for direct simulation of RNA phase separation. It recapitulates a wide range of sequence, length, Mg^2+^ and temperature dependence of the phase separation of RNA homopolymers and triplet repeats, allowing in-depth analysis of the driving forces of RNA phase transitions. Taken together, iConRNA may provide a powerful platform for the study of homotypic RNA phase separation and potentially the heterotypic protein-RNA phase separation.

## RESULTS

### Design of iConRNA: all-Atom to CG mapping and energy function

Each ribonucleotide is represented by six or seven CG beads in iConRNA, similar to what is adopted in Martini-RNA [59], As illustrated in, the phosphate group is represented as one negatively charged bead (B1), while the ribose sugar (including the backbone methylene) is mapped to two beads (B2 and B3). For bases, we use four-bead and three-bead rings to model purines in adenine (A) and guanine (G) and pyrimidines in cytosine (C) and uracil (U), respectively. Detailed all-atom (AA) to CG mapping scheme is given in **Table S1.** Note that, when converting the atomistic model to the CG model, B2 and B2 are fixed at the positions of C4’ and C1’ of ribose, while the center of mass are used for other CG beads. Given the mapping scheme summarized in Figure 1, the total potential energy contains contributions from four major types of interactions,

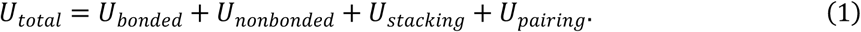

**Figure 1.**
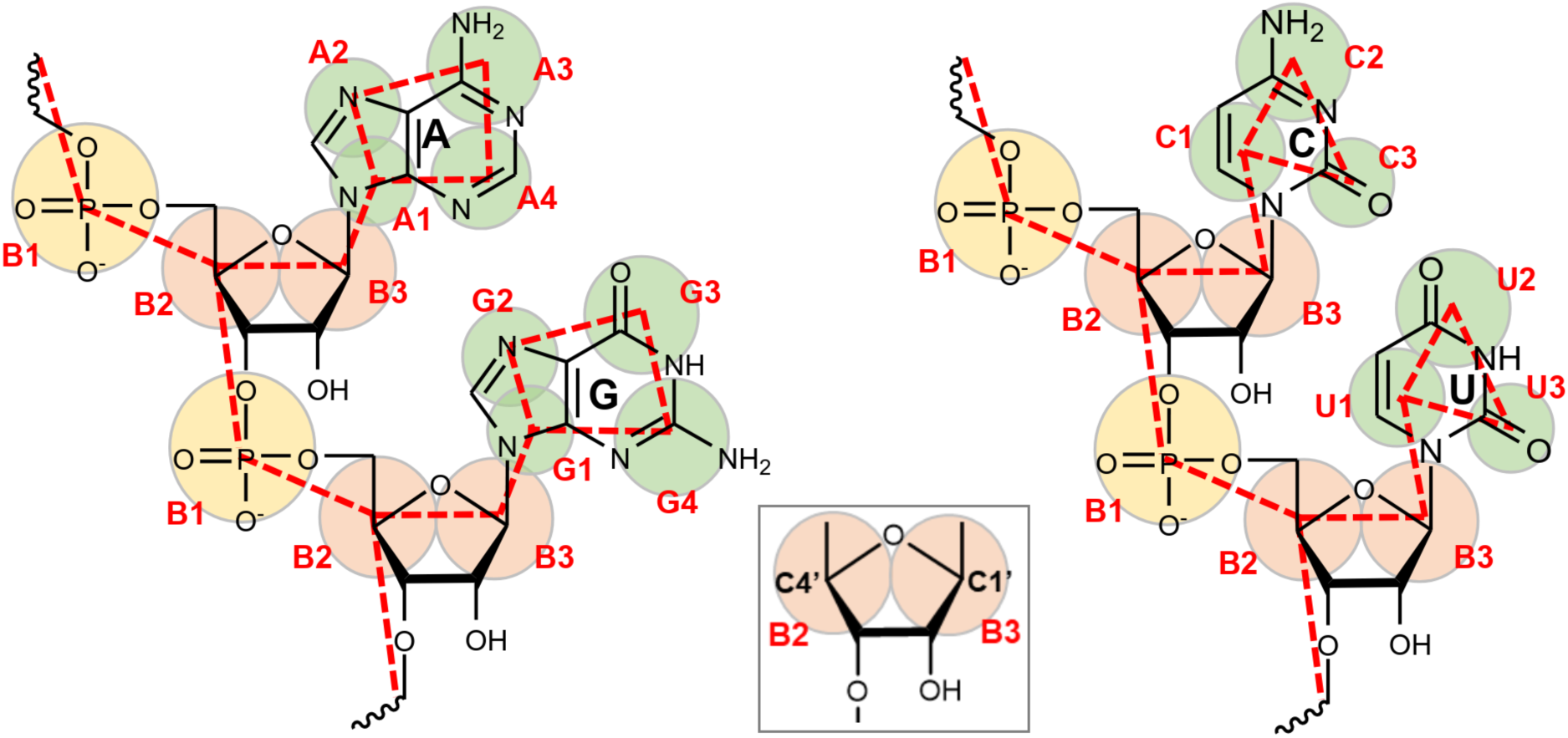
All-atom to CG mapping of RNA nucleotides in iConRNA model. The CG beads are shown by colored circles, which roughly contain their corresponding atom groups shown in the chemical structures. The bead names are given in red labels, and red dashed lines are the pseudo-bonds. B2 and B3 are located at the positions of C1’ and C4’ of ribose sugar, respectively, as shown in the inset.

### Bonded and nonbonded interactions

*U_bonded_* contains contributions from the standard bond, angle, dihedral angle, and improper dihedral angle terms, as used in our previous HyRes protein model [70,71]. The only exception is that a restricted bending (ReB) potential [72] is used for the angle term to prevent the bending angle from reaching the value of 180° and avoid numerical instability of related dihedral terms,

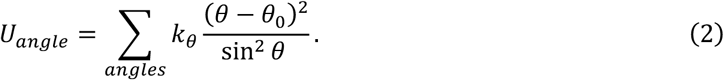

Nonbonded interactions *U_nonbonded_* includes both electrostatic and Van de Waals terms,

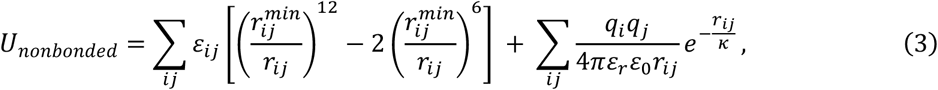

where 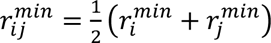, and 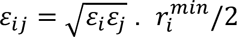 and ε_*i*_ are the vdW radius and interaction strength of the bead *i*. *q_i_* is the charge (-1 for B1 and 0 otherwise), and *r_ij_* is the distance between beads *i* and *j*.*ε* is the permittivity of vacuum. *κ* is the Debye screening length, determined as 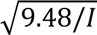 at 300 K, where *I* is the ionic strength in molar. ε_;_is the effective dielectric constant, which is set as 20 at 300 K following the HyRes protein model.

Parameters for bonded terms, including all force constants and equilibrium values, were assigned by reproducing the corresponding distributions derived from all-atom (AA) explicit solvent simulations of tetramer single-strand RNAs (ssRNAs), to provide an implicit description of the structural flexibility at the CG level. A set of ssRNA tetramers as used in the Martini-RNA model [59], including GNRA, CUUG, and UNCG tetramers (where N = any nucleotide and R = A or G), was selected to generate the atomistic reference distributions. For nonbonded parameters, the vdW radius 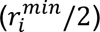 of each CG bead was assigned to reproduce the average total vdW volume of the corresponding groups of atoms derived from the tetramer simulations. The initial values of vdW interaction strengths (ε_5_) were taken from the similar atom groups in the Martini force field [59,73] and then scaled down to the same level of ε_5_ values in the HyRes protein model [70,71] for future compatibility. The overall strengths of vdW interactions were further optimized by a single scaling factor during the optimization of base stacking and pair interactions to produce the structure and chain flexibilities of RNA homopolymers, including the radius of gyration (*R*_g_) of poly(adenylic acid)_30_ (rA_30_) and poly(uridylic acid)_30_ (rU_30_), and the end-to-end distance (*R*_e_) and persistence length (*l*_p_) of (rU_40_).

### Base stacking and pairing interactions

Besides the standard nonbonded interactions, iConRNA also contains explicit base stacking interactions between neighboring bases,

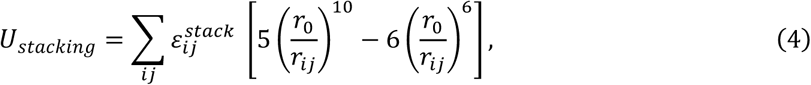

where 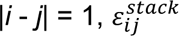 is the stacking strength between bases *i* and *j*. In this work, 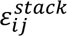 only depends on the types of stacking bases (denoted as *i*//*j*), which are taken from Li and Chen (2023) [74] and summarized in **Table S2** (relative to A//A stacking interaction). To mimic the A-form-like helical structure, this stacking interaction is imposed between two virtual sites on bases. As shown in **Figure S1**, each base *i* has two virtual sites, named S*_i1_* and S*_i2_*, located at the mass centers of the two bonded beads. For example, for base A, S*_i1_* is the mass center of beads A1 and A2 and S*_i2_* is the mass center of beads A3 and A4. In the current version of this model, only two neighboring bases along the RNA chain have stacking interactions. Thus, from the 5’ to the 3’ end, the stacking force between base *i* and *j* is assigned between *S_il_* and *S_ji_*, with a distance denoted as *r_ij_* . The equilibrium distance *r*_0_ between different types of base stacking was set as a uniform value of 0.34 nm.

For base-pairing interactions, only canonical WC A-U and G-C pairs are explicitly included in iConRNA. As shown in **Figure S2**, pairwise interactions U_&’()(!*_ are applied to pairs of hydrogen-bonding CG beads, namely, A3-U2 and A4-U3 for the A-U pair and G3-C2 and G4-C3 for the G-C pair, as

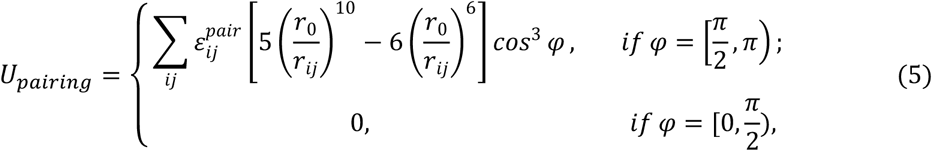

where 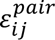 is the strength of hydrogen-bonding pairing, *r_ij_* is the distance between corresponding CG beads. φ is the angle of C2-G3-G2 for the G-C pair and U2-A3-A2 for the A-U pair, which is used to roughly mimic the planar structure of two pairing bases. *r*_0_ is the equilibrium distance for each pair of hydrogen bonding CG beads, which is determined from atomistic structures and summarized in **Table S3**.

To determine the strength of A//A stacking (and thus all stacking interactions), we examine both the radius of gyration (*R*_g_) and the structural propensity of rA_30_ in comparison to the experimental results [75]. The latter is quantified by the orientation correlation function (OCF),

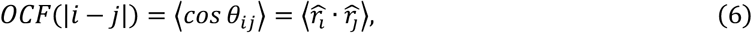

where 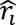 is the normalized bond vector between the *i*th and (*i* + 1)th phosphate groups along the RNA chain. After the determination of base stacking strength, the base pairing strength was optimized to reproduce the melting temperature of referenced double-stranded RNA (dsRNA), including (CAG)_20_ and human telomerase RNA (hTR) hairpin. The strengths of canonical A-U and G-C pairs are fixed at the ratio of 2:3 [74]. The melting curve was measured through the constant volume heat capacity (*C*_V_). The final parameters for various pairing and stacking interactions are given in **Tables S2** and **S3**.

### Treatment of explicit Mg^2+^ ions: concentration and temperature dependence

Screening of electrostatic interactions due to monovalent ions can be treated effectively using the Debye-Hückel-type electrostatic potential (Eqn. 3). as described above. However, such a mean-field treatment is problematic for interactions involving divalent ions such as Mg^2+^ due to ion-ion correlations and complex coordination with the phosphate backbone of RNA [76–79]. To more accurately describe specific Mg^2+^ interactions in RNA structural dynamics and phase separation, we include explicit Mg^2+^ ions in iConRNA. A complication is that Mg^2+^ can interact with the phosphate group either as a fully hydrated ion, mostly through electrostatics, or through first-shell coordination, which has significant non-electrostatic components such as polarization and charge transfer [77,80,81]. In this work, we include an effective Mg^2+^ ion, denoted as Mg, to roughly describe the essential charge screening and coordination effects. This effective Mg has a comparable volume as the fully hydrated [Mg(H_2_O)_6_]^2+^ cluster with a radius of 0.3 nm and +2 charge [81,82].

### Phosphate-Mg interaction

In the mixed salt solution, magnesium binding with RNA depends strongly on the monovalent salt concentration [83]. Previous experimental, theoretical and simulation studies have shown that the ion competition between Na^+^ and Mg^2+^ can be described by the excess Mg^2+^ ions per phosphate, *Δn*_Mg_ = Δ*N*_Mg_/*N*_P_, where Δ*N*_Mg_ is the number of excess Mg^2+^ in the RNA-containing sample compared to the bulk buffer and *N*_P_ is the number of phosphates [82–85]. Even though *Δn*_Mg_ depends on the sequence and structure of RNA as well as the concentration of competing monovalent cations [86], it has been shown previously that *Δn*_Mg_ can be fit to an effective Hill equation [75],

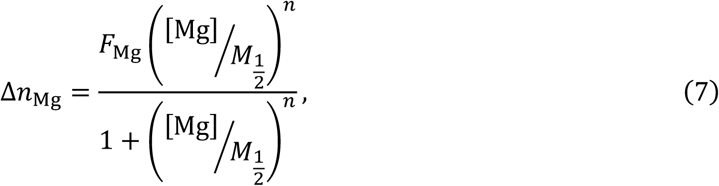

where *F*_Mg_ is the number of excess Mg per phosphate in the limit [Mg]→∞, *n* is the Hill coefficient, and *M*_1/2_ is the competition coefficient (with respect to Na^+^). *F*_Mg_ is generally ∼0.5. In iConRNA, the number of explicit Mg beads in the simulation box is fixed as the number of phosphate groups, which can be considered as an Mg buffer. Increased RNA binding at higher [Mg^2+^] can be effectively described by strengthening the electrostatic interaction between phosphate groups and Mg using a scaling factor *λ*_P-Mg_. As will be shown later, there is an effective Hill relationship between *λ*_P-Mg_ and *Δn*_Mg_ as well, which will be empirically determined and applied to simulating RNA structure and phase separation under different Mg^2+^ concentrations.

### Temperature dependence of phosphate-Mg interaction

To capture the temperature dependence of electrostatic screening, we adopt an empirical relation previously derived from experimentally measured dielectric constants [87],

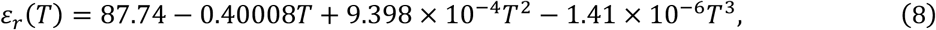

where *T* is the temperature in Celsius. Because the value of ε_;_= 20.0 is used in iConRNA at 303 K, this relation was then scaled down as,

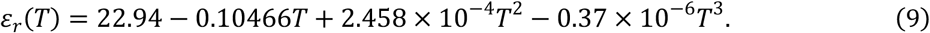

In addition, entropic contributions due to the release of counterions and coordination water molecules strengthen the interaction between reversely charged groups at higher temperatures, which drives the lower critical solution temperature (LCST)-type phase separation of polyelectrolytes (including RNAs) [44,88–93]. The increased interaction between Mg and phosphate can be effectively captured by introducing a temperature dependence in *λ*_P-Mg_, which can be derived from *Δn*_Mg_, from simulations at various temperatures (e.g., see **Figure S3**). As will be shown in Results, a simple linear relation between *λ*_P-Mg_ and temperature also apparently exists and can be empirically derived from atomistic simulations.

### Model parametrization and validation

The force field parameters for the iConRNA model were determined largely in a sequential matter, starting from bonded interactions, then nonbonded and base stacking interactions, and finally, base-pairing interactions. The paramterization was guided by theoretical analysis, atomistic simulations, and experimental measurements. All model dynamic and structured RNAs as well as simulation configurations can be found in **Table S4** and **S5** (see Methods).

### Bonded terms

Parameters involved in bonded interactions were first iteratively optimized to match the corresponding CG and AA distributions, which are summarized in **Figures S4-S7**. Details of all bonded parameters are provided in **Tables S6-S9**. Overall, the CG distributions match the reference AA distributions very well, but show fewer features in several cases, such as the B2-B3-A1/G1-A2/G2 dihedral angle distribution (**Figure S6**). At the atomistic level, the relative orientation of the base and ribose ring is highly flexible and the B2-B3-A1/G1-A2/G2 dihedral angle could sample both cis- and trans-like configurations for ssRNAs, especially at the termini. Thus a soft term is used to cover the main peak of the reference distribution (**Table S8**, *k*ψ = 1.0 kcal/mol).

### Nonbonded and base stacking interactions

These paramters were tuned by examining the salt-dependent conformational properties of homopolymer RNAs in comparison to the available experimental results [75,94]. The key parameters include the overall scaling of vdW interactions and the A//A stacking strength; the latter is used to set all stacking strengths using the relative scales in **Table S2**. The optimized vdW parameters are given in **Tables S10** and **S11**. The final iConRNA model shows a strong ability to describe the structure and dynamics of model ssRNAs and their salt dependence. **Figures 2** and **S8** summarize simulated *R*_g_ and OCF of rU_30_ and *R*_e_ and *l*_p_ of rU_40_ under different salt concentrations in comparison to available experimental data. Without any base stacking between uracils, rU_30_ is a random coil (**Figure S9A**) and its OCF profiles are featureless (**Figure 2B**), also consistent with other experiments [94–96]. Even though *R*_g_ (and *l*_p_) from iConRNA simulations is slightly over-estimated compared to the experiment, strong agreement on their salt dependence suggests that the model captures well the balance between electrostatic and vdW interactions. Curiously, it proved difficult to further reduce the overestimation of *R*_e_ of rU_40_ (**Figure 2C**). Similar *R*_e_ deviations were reported in some other CG RNA models [65,67]. Our analysis suggested that this was mainly due to the strong repulsive interactions along the negatively charged backbone, leading to a longer persistence length (**Figure 2D**). Weakening of electrostatic interactions could rescue *R*_e_ of rU_40_, but this would severely reduce the ability of the model to capture salt dependence of structural properties. With the A//A stacking strength set to 2.05 kcal/mol, iConRNA correctly predicts the helical structure of rA_30_ (**Figure S9B**), and both the OCF profiles and their salt dependences show high consistency with experimental results, as illustrated in **Figures 2F** and **S8D-F**. Nonetheless, rA_30_ appears to be slightly more dynamic in iConRNA, leading to slightly shallower features in the OCF profiles. This is likely due to the smooth potentials of the CG model and the higher freedom of the bases. Thus, the strengths of the dihedral angles B1-B2-B3-A1(G1) and B2-B3-A1(G1)-A2(G2) were increased from 8.0 and 5.0 kcal/mol to 35.0 kcal/mol to prevent the deformation.

**Figure 2.**
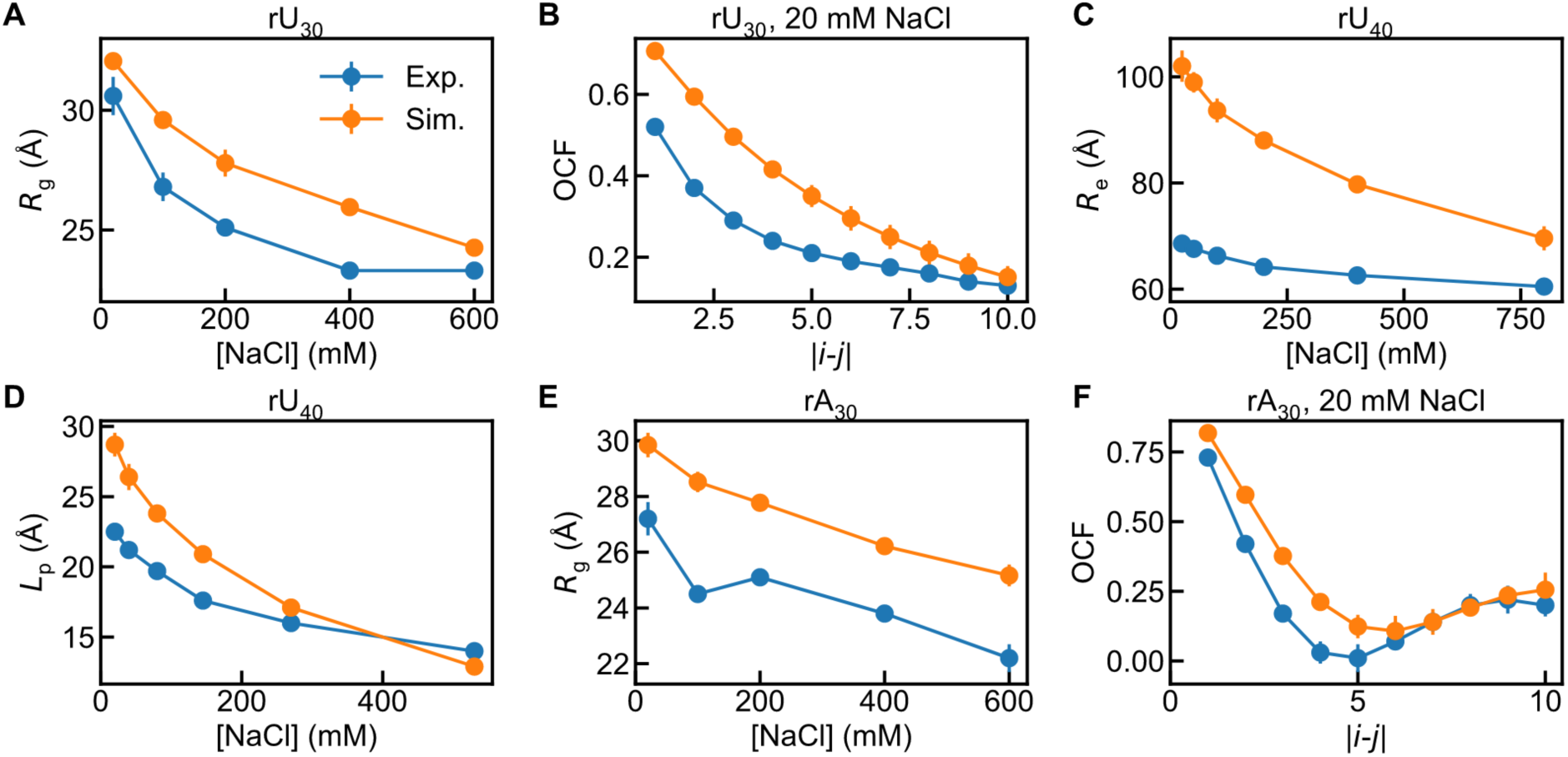
Salt-dependent chain properties of homopolymer RNAs. (**A**) The radius of gyration (*R*_g_) of rU_30_ as a function of NaCl concentration. (**B**) Orientation correlation function (OCF), as defined in Eqn. 6, of rU_30_ with 20 mM NaCl. (**C, D**) The end-to-end distance (*R*_e_) and persistence length (*L*_p_) of rU_40_ as a function of NaCl concentration. (**E**) *R*_g_ of rA_30_ as a function of NaCl concentration. (**F**) OCF of rA_30_ with 20 mM NaCl. The experimental results [65,67] are shown in blue whereas the iConRNA simulation results are in orange. The error bars of simulation results were estimated from block analysis.

### Base pairing

Their strengths were determined at the last. In iConRNA, the ratio between A-U and G-C pairing strength was fixed at 2:3 as frequently done in CG RNA models [74,97]. Both pairs form two (pseudo-)hydrogen bonds to help retain the planar geometry (**Figure S2**). For the A-U pair, hydrogen bonds A3-U2 and A4-U3 take equilibrium distances of 0.33 nm and 0.40 nm, respectively. For the G-C pair, the equilibrium distances of G3-C2 and G4-C3 are 0.33 nm and 0.38 nm. In the final model, the strength of each hydrogen bond of the A-U pair was set as 1.44 kcal/mol, while it was 2.15 kcal/mol for the G-C pair (**Table S3**). With these parameters, iConRNA reproduces the melting temperatures of both (CAG)_20_ [98] and hTR hairpin [99] (**Figure 3A**) as well as the Beet Western Yellow Virus pseudoknot (**Figure S10**). Note that experimental melting profiles frequently contain minor peaks, suggesting the existence of alternative structures and/or multiple unfolding intermediates. iConRNA cannot capture all these details, which is not surprising due to the CG nature and absence of non-canonical base pairing interactions.

**Figure 3.**
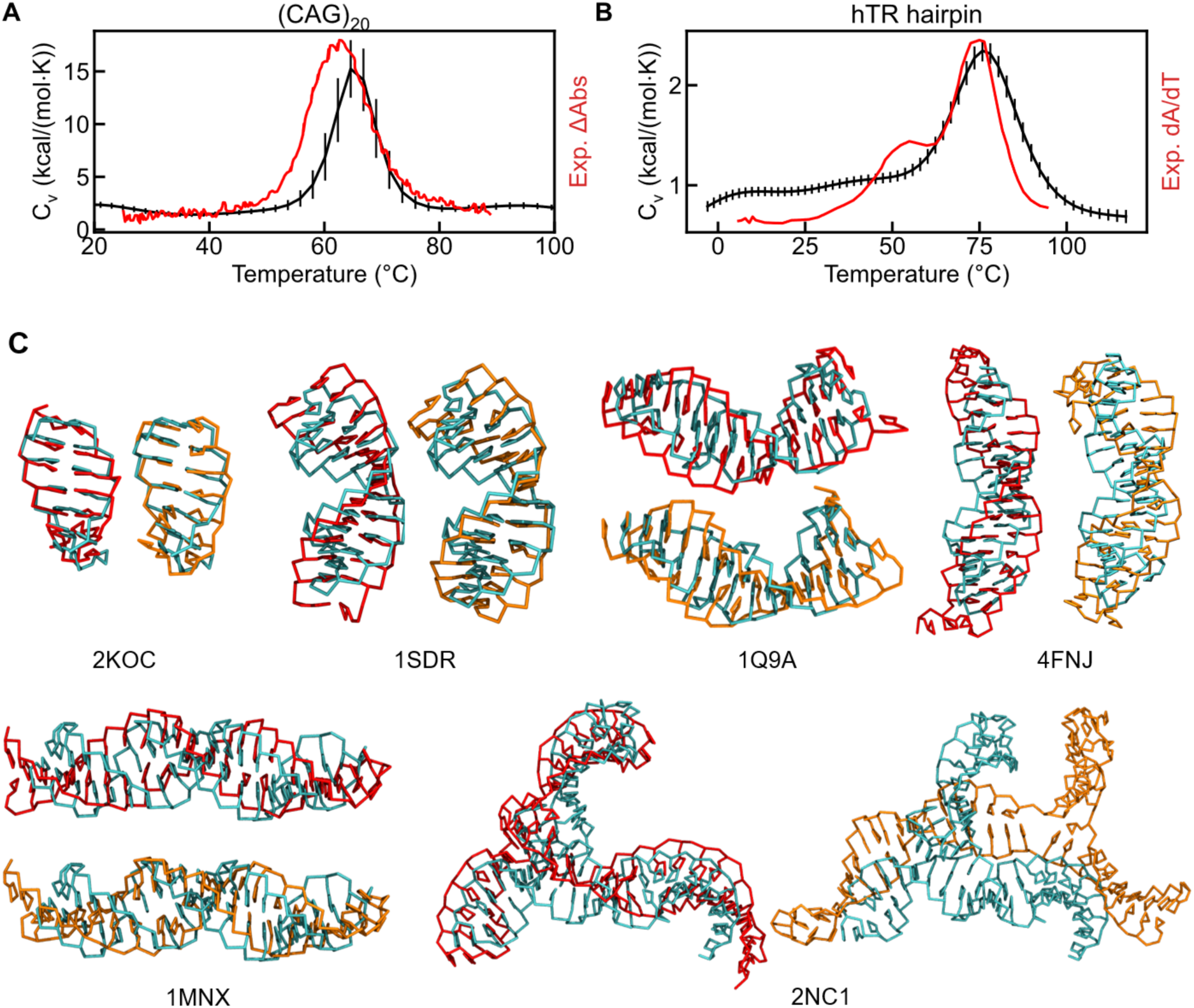
Melting temperature and folding of dsRNAs. (**A, B**) Heat capacity (C_v_) of (CAG)_20_ (**A**) and hTR hairpin (**B**). The uncertainties of simulation results were calculated using bootstrap resampling [100]. Experimental data are red lines, where the change of UV-absorbance (ΔAbs) at 260 nm and the first derivative of UV-absorbance with respect to temperature (dA/dT) at 280 nm are used as the melting profiles for (CAG)_20_ [98] and hTR hairpin [99], respectively. (**C**) Comparison between experimental crystal structures (cyan) and predicted structures of dsRNA (PDB id: 2KOC, 1SDR, 1Q9A, 4FNJ, 1MNX, 2NC1) (see **Table S5** for sequences). The predicted structures were obtained from single chain simulation starting from crystal structures (red) or linear chains (orange), respectively.

We further examine the ability of iConRNA to fold small dsRNA structures. Starting from crystal structures and coil single strands, we simulated a series of simple dsRNA, including 2KOC, 1SDR, 1Q9A, 4FNJ, 1MNX, and 2NC1 (in PDB id). As illustrated in **Figure 3C**, the model can basically predict the hairpin structures with A-form-like helices, although the detailed helical characteristics are not always consistent with those in crystal structures. For complex RNA like 2NC1, some special interactions including intra-molecular cross-base stacking (not the stacking between neighboring bases) and non-canonical base pairing are necessary to retain the structure, which is not included in iConRNA. As such, the folded structure sampled by iConRNA displays large deviations. The ability of iConRNA to correctly fold small RNAs is notable, as the model has been mainly designed to capture the structural properties of dynamic RNAs. The apparent balance between structure and dynamics provided by iConRNA is an important strength for capturing the complex interplay of various interactions and structural propensities of RNA in phase separation.

### Parameterization of explicit Mg^2+^ interactions

As described above, with a fixed number of Mg in the simulation box, the actual effect of Mg^2+^ concentration on binding to RNA backbone is effectively modeled through scaling the phosphate-Mg interaction strength with *λ*_P-Mg_, which is related to the excess number of Mg^2+^ per phosphate group (*Δn*_Mg_). We first determined the optimal *λ*_P-Mg_ by reproducing the [Mg^2+^] dependence of *R*_g_ of rA_30_ and rU_30_ (**Figures 4A** and **4B**) [75]. The final optimized *λ*_P-Mg_ at various [Mg^2+^] are summarized in **Table S12**. The OCF profiles derived from the simulations at 1 and 2 mM [Mg^2+^] are in reasonable agreement with the experimental references (**Figure S11**), but display larger discrepancies compared to Mg^2+^-free conditions (e.g. **Figure 2**). This is likely due to that the effective Mg bead in iConRNA is as big as the fully hydrated Mg^2+^, which does not account for the dehydration of chelated Mg^2+^ ions and resulting structural differences.

**Figure 4.**
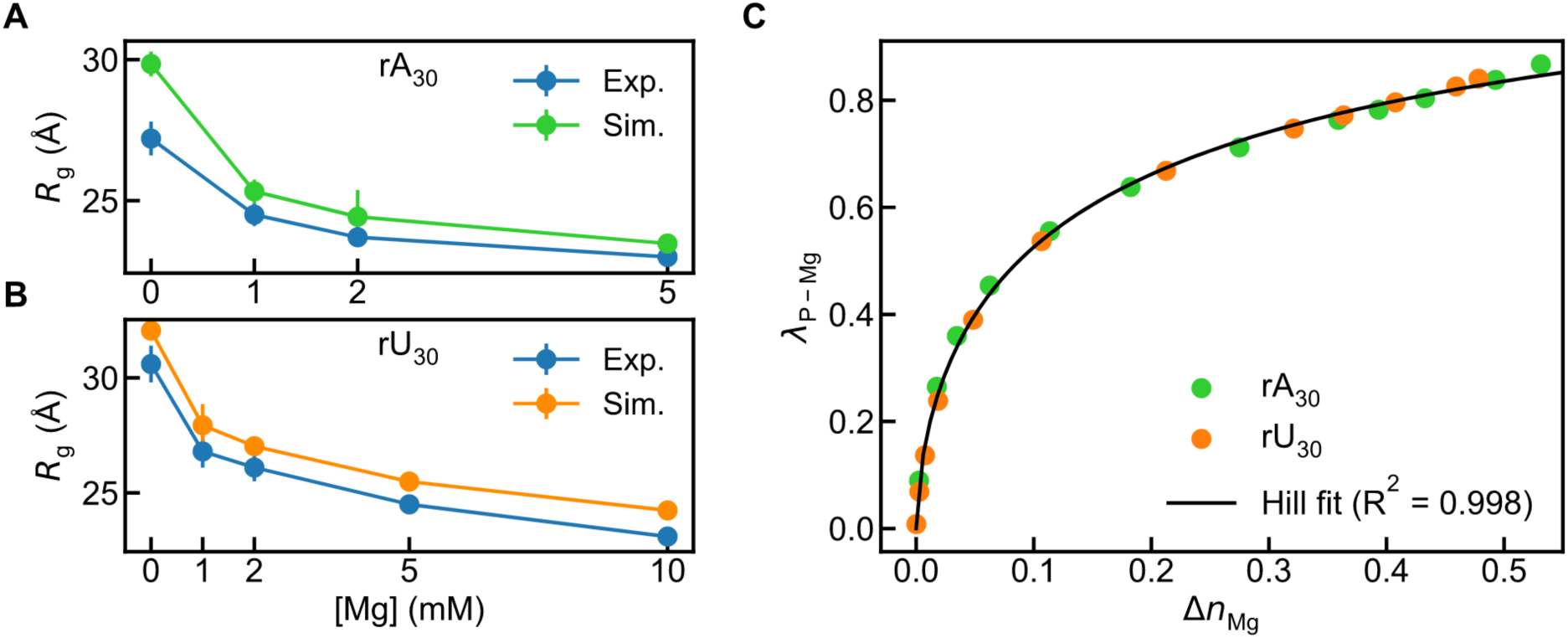
Calibration of phosphate-Mg interaction. (**A, B**) [Mg] dependence of *R*_g_ of rA_30_ (**A**) and rU_30_ (**B**). Experimental data (blue lines) are taken from Ref [75]. The error bars of simulation results are the standard deviations of block averages. (**C**) Hill fit of optimal *λ*_P-Mg_ against *Δn*_Mg_ with *R*^2^ = 0.998.

Interestingly, a Hill fit of the optimal *λ*_P-Mg_ against *Δn*_Mg_ was determined from reproducing *R*_g_ of rA_30_ and rU_30_ (**Figure 4C**),

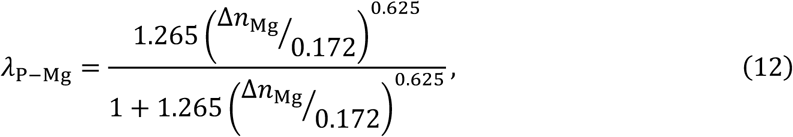

with *R*^2^ of 0.998. This suggests that, for any RNA system in the mixed Na^+^/Mg^2+^ solution, the effective phosphate-Mg interaction can be determined through *Δn*_Mg_ without the need for reparameterization based on measured or predicted *R*_g_ of the RNA of interest. Note that *Δn*_Mg_ itself can be determined through experiment or predicted using simulation or theory [75,82–85,101–104]. For rA_30_ and rU_30_, *Δn*_Mg_ and its fit to the Hill equation have been determined experimentally [75]. For RNA triplet repeats to be studied in this work, we performed additional all-atom simulations to determine *Δn*_Mg_, using (CAG)_31_ as a representative system In the simulations. The calculated *Δn*_Mg_ converges well within the 100 ns simulation timeframe (**Figure 5A**), and can be fitted well to the non-cooperative Hill-equation (Eqn. 7) (**Figure 5B**). Clearly, (CAG)_31_ has a lower level of Mg^2+^ binding compared to RNA homopolymers. The fit, given in **Table S13**, will be used for the simulation of all RNA triplet repeats in this work.

**Figure 5.**
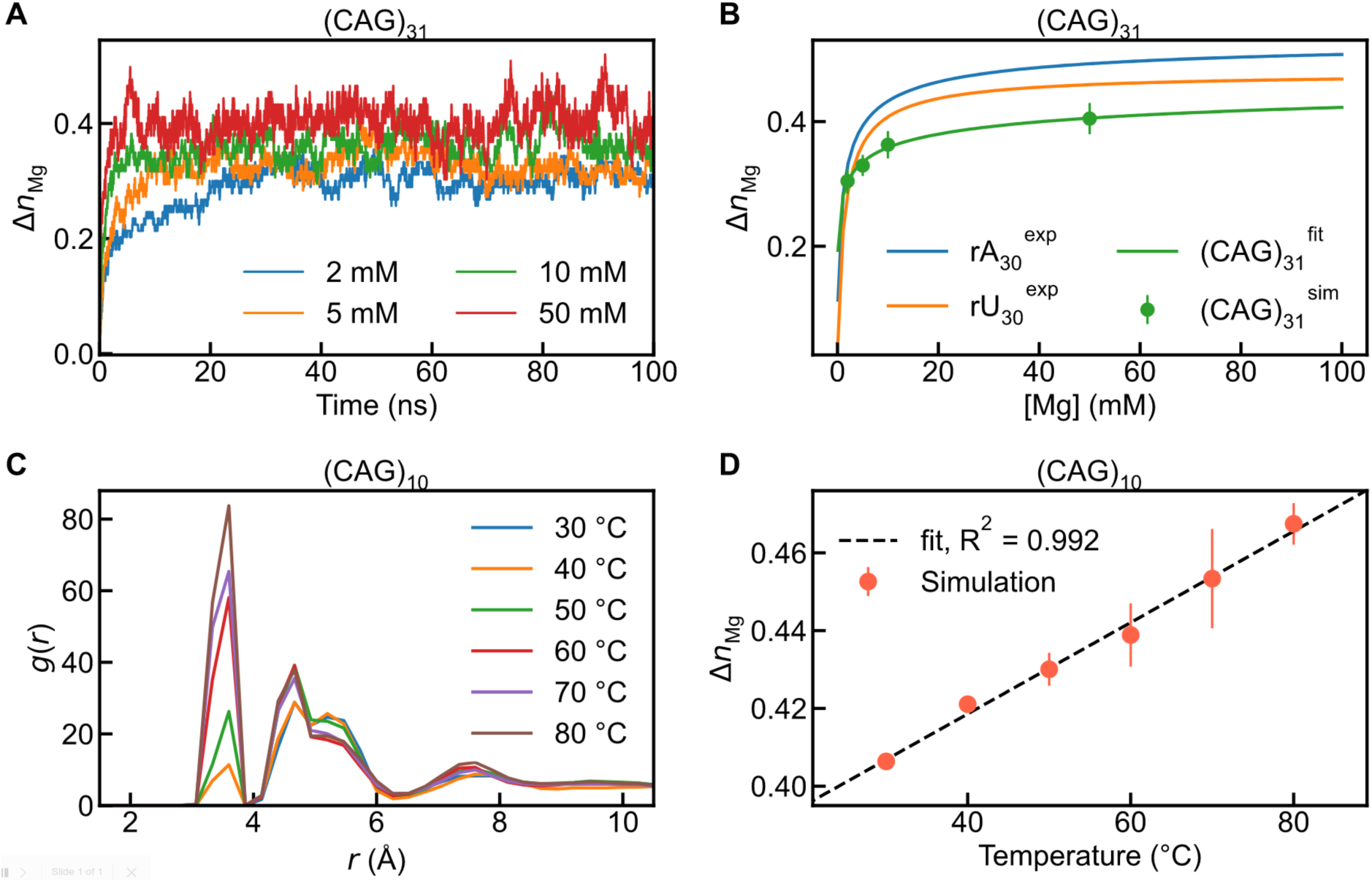
Determination of *Δn*_Mg_ and its temperature dependence for CAG repeat. (**A**) *Δn*_Mg_ of (CAG)_31_ as a function of simulation time under different Mg^2+^ concentrations. (**B**) Hill fits of *Δn*_Mg_ to [Mg] for rA_30_, rU_30_, and (CAG)_31_. Labels for legend entries: “exp” is for experimental data, “sim” is for simulation data estimated from the last 50 ns, and ‘fit’ is for the fit curve. Error bars for simulation data were evaluated as block deviation. (**C**) RDFs of Mg^2+^ ions around phosphorus atoms at different temperatures. (**D**) Linear fit of *Δn*_Mg_ to temperature (in Celsius). Simulation results are given in orange scatters with error bars estimated from multiple replicas. The black dashed line shows the linear fit with *R*^2^ = 0.992.

The temperature dependence of *Δn*_Mg_ was further investigated through all-atom simulations of short RNA triplet repeats (CAG)_10_. The distribution of Mg^2+^ around phosphate groups was quantified within the temperature range of 303 K to 353 K under 25 mM NaCl and 50 mM MgCl_2_. Similar to previous simulations of polyphosphate [44], the resulting radial distribution functions (RDFs) show that the binding of Mg^2+^ ions with phosphate groups strengthens with the increase in temperature (**Figure 5C**). The increase is particularly pronounced in the inner-sphere binding within 4 Å (the first peak), where the Mg^2+^ ion was directly coordinated with the oxygen of the phosphate group (**Figure S12**) [105,106]. Interestingly, a linear dependence exists between *Δn*_Mg_ and the temperature (**Figure 5D**),

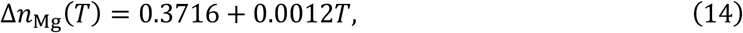

where *T* is in Celsius, or

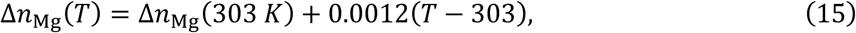

where *T* is in Kelvin, and *Δn*_Mg_ (303 K) is the excess binding Mg^2+^ per phosphate at 303 K. The linear relationship suggests that the entropic effects do dominate the temperature dependence of Mg^2+^ binding to RNA backbone phosphates as expected. As such, the above relationship should roughly hold for all RNAs, and it will be used to model the temperature dependence of Mg^2+^-phosphate interaction for all subsequent simulations in the work.

Notably, the number of Mg is fixed to be the same as the number of phosphate groups, which does not change with the box size. In principle, *λ*_P-Mg_ should be re-optimized for different box sizes to reproduce *Δn*_Mg_. Therefore, we further evaluated the effect of box size with a fixed number or concentration of Mg on RNA structure. The results show that, without adjusting *λ*_P-Mg_, only up to 4% change was observed in *R*_g_ even with box sizes varying from 15 nm to 50 nm in both setups (**Figure S13**). Therefore, the final model will not correct for the box size dependence of *λ*_P-Mg_ with a fixed number of explicit Mg beads. In summary, we developed a protocol for setting up explicit Mg ions in iConRNA simulations. The number of Mg was first set as the number of phosphate groups. Next, *Δn*_Mg_ was determined based on reported experiments, theories or atomistic simulations at the temperature of interest. If temperature dependence needs to be considered, *Δn*_Mg_ at different temperatures could be predicted by **Eqn. 14** or **15**. Finally, *Δn*_Mg_ was converted into *λ*_P-Mg_ following **Eq 12** as an input parameter of the iConRNA simulation script.

### Phase separation of poly(rA) and poly(rU)

Recent works have demonstrated that RNA homopolymers can undergo homotypic phase separation in the presence of multivalent cations, and exhibit very different phase behaviors when composed of different nucleotides [43–45,92,107]. For example, rA_30_ readily undergoes phase separation under 50 mM MgCl_2_, while poly(rU) remains soluble under the same conditions [45]. We first examined the ability of iConRNA to recapitulate the phase separation of RNA homopolymers in the presence of Mg^2+^. Starting from dispersed initial states with 240 copies in a cubic box of 200 nm in dimension (50 μM), rU_30_ remained dispersed and can only form small oligomers (up to pentamers) with temperatures ranging from 303 to 353 K even with 50 mM MgCl_2_ (and 25 mM NaCl) (**Figure S14**). In contrast, rA_30_ clearly shows LCST-type phase separation behaviors (**Figure 6A** and **6B**), as observed in experiments [44]. At lower temperatures (300 K and 313 K), only dimers and trimers were obtained. Starting from 323 K, stable droplets can be observed and the number of monomers in the dilute phase decreases rapidly with the increase in temperature (**Figure 6A**). We note that experimentally rA_30_ undergoes phase separation as low as 303 K with 50 mM MgCl_2_ [45], which indicates that iConRNA slightly underpredicts the phase separation propensity of rA_30_. A possible reason is that only the stacking between neighboring bases is included in our model, such that rA_30_ is unable to form intermolecular base stacking interactions potentially involved in the phase separation of rA_30_.

**Figure 6.**
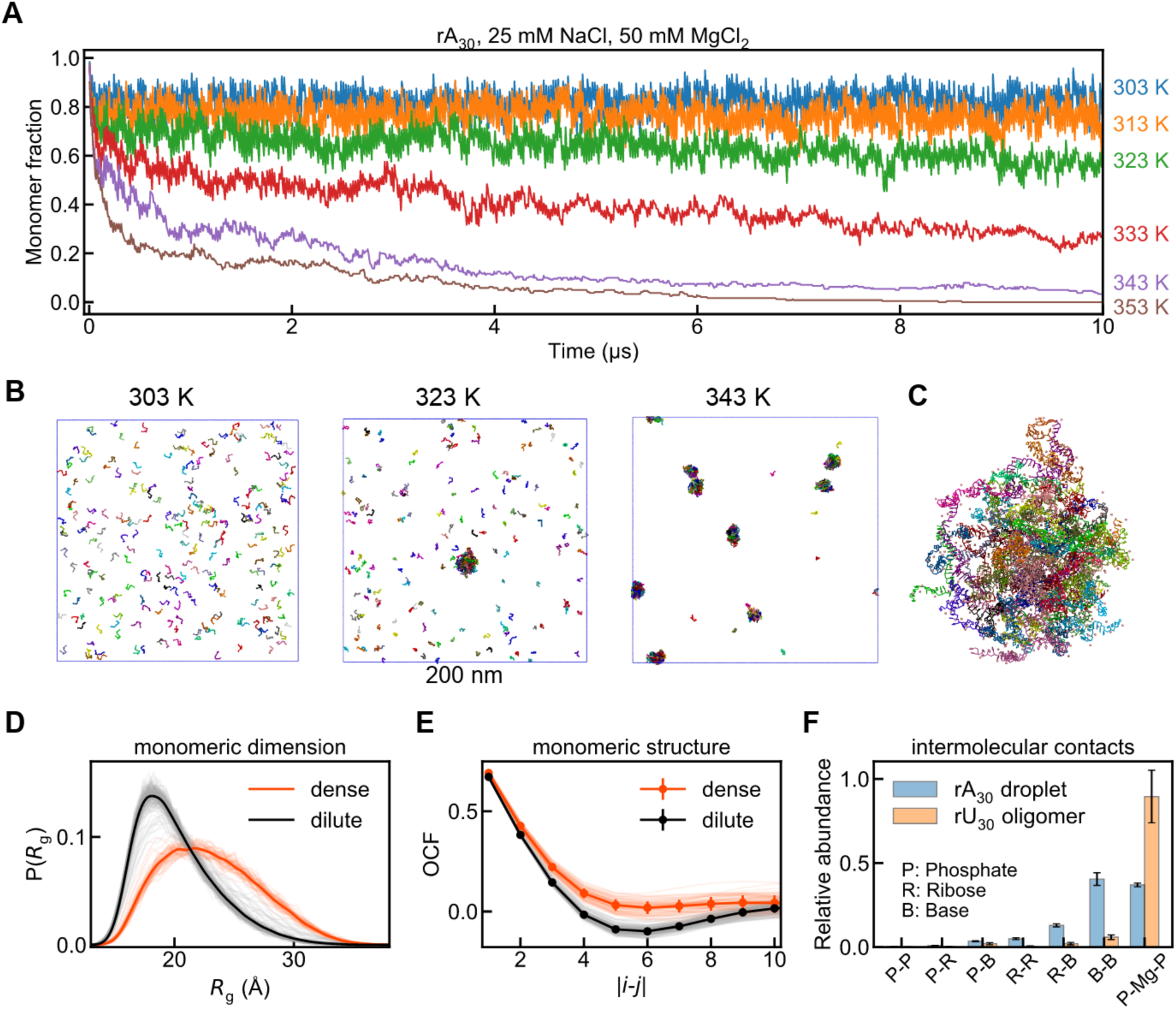
Phase separation of rA_30_. (**A**) Monomer fractions as a function of simulation time at different temperatures under 25 mM NaCl and 50 mM MgCl_2_. The total RNA concentration is 50 µM. (**B**) Final snapshots at 303, 323, and 343 K. Mg were omitted for clarity. The box size is 200 nm. (**C**) Snapshot of a representative rA_30_ droplet. Each RNA chain is colored differently, while Mg ions are shown as pink beads. (**D, E**) *R*_g_ distributions and OCFs of RNA chains in the dense (orange and red lines) and dilute phases (black and grey lines) at 323 K. Light-coloured lines plot the distributions of individual chains, and their averages in bold lines. Error bars are the standard deviation of related chain groups. (**F**) The relative abundance of various intermolecular contacts within rA_30_ droplets (blue) and rU_30_ oligomers (orange).

Within the condensate, rA_30_ forms extensive intermolecular interactions, apparently through a wide range of interaction involving backbone phosphate, side chain bases as well as Mg^2+^ ions (**Figure 6C**). We further characterized the conformations of rA_30_ inside the dense phase. The results show that RNA chains within the condensates are more extended, exhibiting a larger average value and broader distribution of *R*_g_ (**Figure 6D**), Furthermore, each individual *R*_g_ distribution (the light orange lines) is very close to the average one (bold orange line), which indicates a relatively homogeneous conformation distribution of rA_30_ inside the condensates. Additionally, we found condensates are less favorable for the single-strand helical structure of rA_30_ compared to the dilute phase. OCFs of RNA chains within the condensates exhibit flatter trends and fewer structural details (**Figure 6E**), indicating weakened intramolecular base stacking. In the crowded environment of condensates, intermolecular interactions among rA_30_ chains are highly enhanced, which results in the partial disruption of local intramolecular base stacking [108]. Intermolecular contact analysis indicates that within the condensate, rA_30_ forms extensive intermolecular interactions through phosphate-Mg electrostatic interactions and base-mediated inter-chain vdW interactions (**Figure 6F**). The latter class of interactions is the distringuishing feature between rA_30_ and rU_30_. The base-base interactions observed in the droplet resemble A-A stacking and non-canonical A-A pairing, although iConRNA doesn’t contain these terms explicitly. In addition, there is a significant level of base-sugar interactions in rA30 condensates. Clearly, due to weaker base-mediated interactions, phosphate-Mg electrostatic interactions, the major driver of oligomerization and a common feature of RNA backbone, are not sufficient to drive phase transitions of rU_30_ by themselves.

### Phase Transitions of RNA triplet repeats

Nucleotide triplet repeats have been implicated in several neurological and neuromuscular disorders including Huntington disease [46,48,49]. Recently, it was observed that CAG/CUG repeat-containing RNAs could undergo phase separation and gelation, which could be a contributing factor to neurological disease [40]. The complex temperature, sequence and Mg^2+^ dependence of the phase separation of CAG/CUG repeats and additional percolation transitions within the condensate have been further characterized in detail [44]. The formidable complexity of trinucleotide repeat phase separation provides a challenging test case for evaluating the strengths and weaknesses of the new RNA CG model and for dissecting the molecular driving forces of RNA phase separation.

We first examine the ability of iConRNA to capture the structures of these triplet repeats in the monomer state. It has been shown that CAG/CUG repeats can form stable hairpins, while the CUU repeat remains a single strand without any higher-order structure [49,109,110]. As shown in **Figure S15A**, starting from single strands, iConRNA successfully predicts the hairpin structures of (CAG)_31_ and (CUG)_31_ and the disordered single strand of (CUU)_31_. Consistent with experimental observations [110], CAG hairpins were observed to be more stable than CUG ones. The backbone RMSD is 11.8 ± 3.5 Å for CAG, but 34.1 ± 12.4 Å for CUG. Although only canonical A-U and G-C pairing interactions are included in our model, some semi-stable A-A pairs can be found in CAG hairpins due to neighboring A-U and G-C pairs and base stacking interactions, while U-U pairs are very dynamic in CUG hairpins.

We then calculated the phase diagrams of (CAG)_31_ RNA as a function of magnesium concentration (**Figure 7A**) and RNA concentration (**Figure 7B**). As detailed above, for CAG repeats, *Δn*_Mg_ and its temperature dependence were first determined through all-atom simulations. Similar to rA_30_, (CAG)_31_ undergoes LCST-type phase separation, where the transition temperature decreases with the increase of [Mg], indicating the intrinsic entropic contribution of phosphate-Mg interactions to phase separation. Notably, an excellent agreement was observed between simulations and experiments for both [Mg]-dependence and [RNA]-dependence [44]. For the [Mg]-dependence, the critical temperature is slightly overestimated at the low-Mg^2+^ region (below 25 mM) but slightly underestimated at the high-Mg^2+^ region above 50 mM. It might result from the corresponding under- or over-estimation of excess Mg^2+^ binding in the all-atom simulations. In the 1-Φ region, at the temperatures close to the phase boundary, there always exist small RNA oligomers (from dimer to pentamer, left panel in **Figure S15B**), which are dynamic and undergo fast exchange with monomers in the dilute phase.

**Figure 7.**
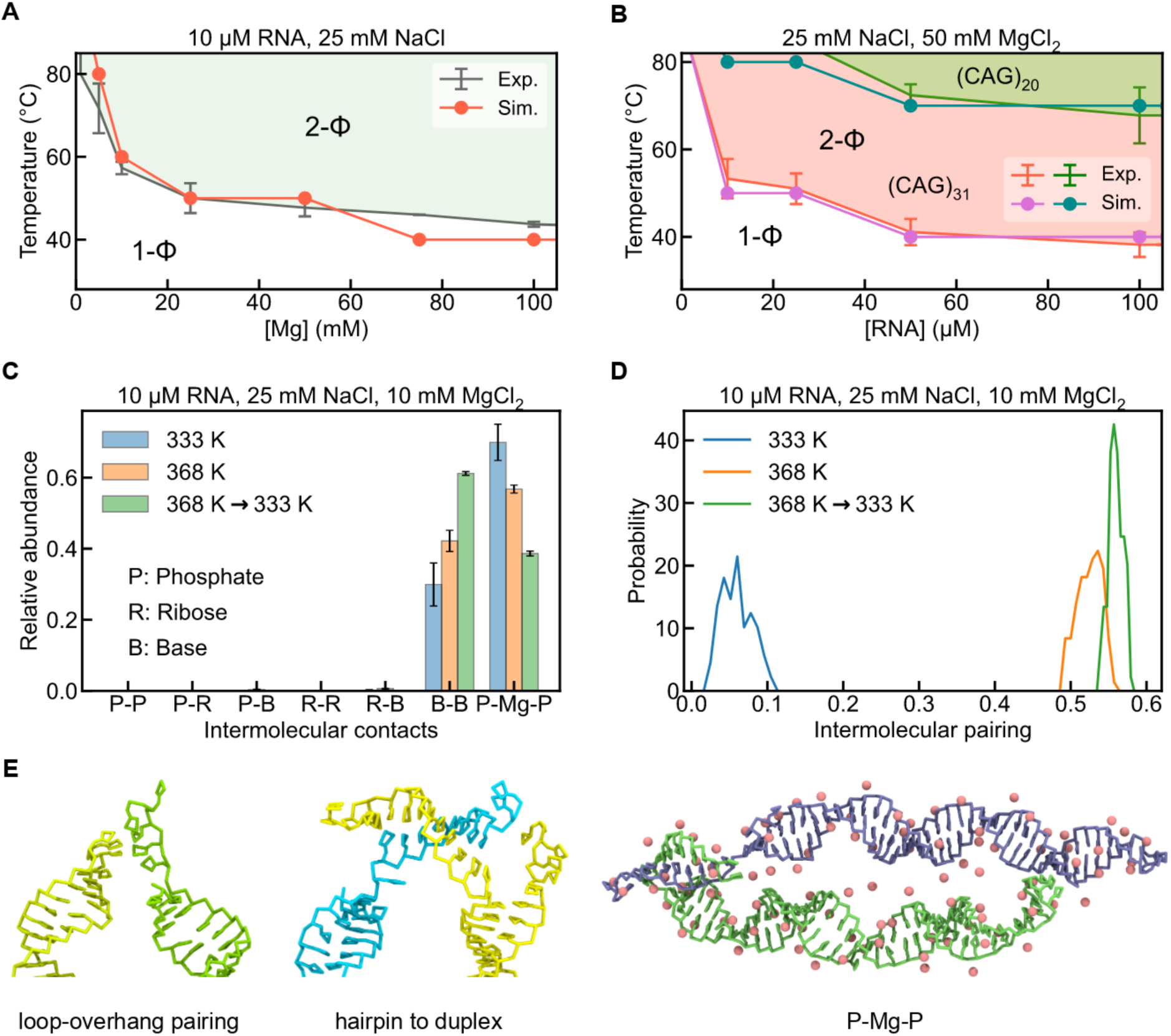
Phase separation of RNA triplet repeats. (**A**) Phase diagram of 10 μM (CAG)_31_ under 25 mM NaCl, where transition temperatures are plotted as a function of [Mg]. (**B**) Phase diagrams of (CAG)_31_ and (CAG)_20_ under 25 mM NaCl and 50 mM MgCl_2_, where transition temperatures are plotted as a function of [RNA]. Experimental 2-phase regions of (CAG)_31_ and (CAG)_20_ are shown in pink and lime areas. In B and C, experimental phase boundaries are plotted as lines with error bars, while simulation ones are lines with filled circles. (**C, D**) Relative abundance of intermolecular contacts (**C**) and the fraction of intermolecular base pairing (**D**) within CAG-repeats condensates obtained at different temperatures. 368 Kè333 K indicates condensates obtained from annealing those originally formed at 368 K to 333 K. (**E**) Representative snapshots of various types of intermolecular interactions within the condensate. Pink spheres represent the effective Mg beads.

We further evaluated the ability of iConRNA to capture the length dependence of LCST-type phase transition of CAG repeats observed experimentally [44]. As summarized in **Figure 7B**, decreasing the RNA length to 20 repeats dramatically weakens the propensity of CAG repeats phase separation. In particular, the critical temperature increases throughout the concentration range examined. For example, it increases from 40 to 70 °C at 100 μM RNAs in simulation, while in experiments, it increases from 38.0 °C for 100 μM (CAG)_31_ RNA to 59.9 °C for 155 μM (CAG)_20_. In addition, under all the conditions tested for (CAG)_20_ and (CAG)_31_ repeats in **Figure 7B**, (CAG)_10_ repeats didn’t undergo any phase separation. For example, under the most favorable conditions, 80 °C and 100 μM RNAs, only a few dimers and trimers were found throughout the simulation box (**Figure S16**).

Experimentally, it was reported the purine-to-uracil substitution could suppress LCST phase transitions [44]. Compared to (CAG)_31_, (CUG)_31_ exhibited slightly lower phase separation propensity, and (CUU)_31_ did not undergo phase separation under the same conditions. iConRNA simulations correctly predicted significant reduction in the phase separation propensity of (CUU)_31_, whch requires much higher temperature (and thus stronger Mg^2+^/phosphate interactions) to phase separate in simulation (**Figure S15C**). However, (CUG)_31_ shows a very similar phase boundary with (CAG)_31_, indicating the weakening effect of A-to-U substitution for (CAG)_31_ is not fully captured by the current model. The limited ability of iConRNA to capture the negative effect of purine-to-uracil substitution in phase separation of trinucleotide repeats suggests that there is still room for improvement in balancing various base-mediated interactions within iConRNA, particularly regarding the strength of nonspecific vdW interactions between uracils. Furtrhermore, the excess binding Mg^2+^ derived from atomisitic simulations may have been over-estimated (**Figure 5**), leading to an over-estimated *λ*_P-Mg_ used in iConRNA simulations.

### Molecular driving forces of RNA triplet repeats condensation: the role of RNA folding

The ability of iConRNA to accurately recaptitulate the length, concentration and Mg2+ level dependence of (CAG)_31_ phase diagrams (**Figure 7A-B**) allows dissection of the contributions of various molecular forces towards driving phase separation. Analysis of intermolecular contacts within the condensate (**Figure 7C-D**) show that, similar to poly(rA), two major contributions come from the Mg-mediated electrostatic interactions (P-Mg-P) and base-base interactions. The former indicates the necessity of divalent Mg^2+^ ions, which bridge RNA backbones to promote their condensation (**Figure 7E**). Besides, this bridging effect also results in the preference of parallel arrangements of multiple hairpins, further promoting the probability of intermolecular base pairing. The base-base interactions were observed to be dominanted by G-C pairing, which occurs through loop, overhang, and tail regions of hairpins **(Figure 7E**). The latter promotes the hairpin to duplex transition during phase separation and further arresting and aging to gel. As as result, the RNA become more expanded within the condensate.

The apparent competition between hairping and duplex formation suggests an important role of intramolecular folding of RNA in phase separation. This is particularly evident when examining the temperature dependence of the distributions of P-Mg-P interactions and base pairing in condensates. Below the melting temperature, the hairpin structures of CAG repeats are stable in both dilute and dense phases, where over 90% G/C residues form intramolecular pairing (**Figure 7D**). Only less than 10% of base pairing are intermolecular, mainly found between loops and overhangs and between two overhangs. Therefore, Mg^2+^-mediated electrostatic interactions are the major driver of phase separation. At 368 K, above the (CAG)_31_ meting temperature, the hairpin structures were partially denatured and more dynamic, which resulted in dramatic increases of intermolecular base pairing in the condensates, to slightly above 50% (**Figure 7D**). Interestingly, even though P-Mg interactions are also stronger at higher temperature, they account for a smaller percentage of intermolecular interactions at 368K wheras the base-base interactions become more abundant (**Figure 7C**). Interestingly, when we anneal the condensates formed at 368 K to 333 K, we observed a significant increase of additional intermolecular base pairing, while Mg^2+^-mediated backbone interactions were further reduced (**Figure 7C-D**). The implication is that the condensate would undergo gelation and the phase transition is not reversible as observed experimentally [44], even though our current simulations are not long enough to directly observe full gelation.

## DISCUSSION

We have developed a high resolution coarse-grained RNA model with explicit magnesium ions, named iConRNA, for molecular simulation of dynamic RNAs and their phase transitions. Representing each nucleotide using 6 or 7 CG beads, iConRNA is designed to explicitly describe a diverse range of interactions that RNA can make through its backbone phosphate and side chan bases, including electrostatics, vdW interactions, base stacking and base pairing. With careful pameterization using theory, atomistic simulation and experimental data, the model is able to capture the essential conformational features of flexible RNAs and at the same time fold small RNAs including tetraloop, duplex and hairpins. One of the major challenges in RNA modeling is describing the interactions with Mg^2+^ ions, which are important in RNA structure and phase separation and display complex structure, ion concentration and temperature dependence. For this, we developed an efficient framework for including explicit effective Mg^2+^ ions in iConRNA. The sequence, concentration and temperature dependence of Mg^2+^-mediated interactions are captured by parametrizing a scaling factor for effective Mg^2+^ ion/phosphate interaction strengths, based on excess Mg^2+^ binding to RNA. The later property can be measured experimentally or derived from atomistic simulations and/or theory. The resulting iConRNA with explicit Mg^2+^ successfully captures the Mg^2+^ dependence of conformational properties of flexible RNAs, and reproduces a wide range of nontrivial phase separation properties of homopolymer and trinucleotide repeat RNAs, including length, sequence, Mg^2+^, and temperature dependence.

The success of iConRNA in recapitulating the phase behaviors of homopolymer and trinucleotide repeat RNAs allows carful dissection of how the interplay of various backbone and base-mediated interactions contribute towards driving phase separation. The analysis reveals the major roles of intermolecular base pairing and Mg^2+^-bridged backbone phosphate interactions in RNA phase separation. For homopolymers, nonspecific base-base and base-ribose interactions are mainly responsible for supplementing Mg^2+^/phosphate interactions to drive phase separation, such as for rA_30_ vs rU_30_. For trinucleotide repeats, intermolecular base pairing is the major driving force besides Mg^2+^/phosphate interactions in phase separation. RNA folding has a central role in controlling the competition between intramolecular and intermolecular base pairing, which is also a key process underlying the complex temperature and Mg^2+^-dependence of the phase diagram and likely gelation and reversibility. We note that cellular RNA foci form at room temperature, which is usually below the melting temperature of (partially) structured RNAs. The dynamic and structural heterogeneity of driver RNAs, as well as how these properties respond to cellular conditions and/or binding of ions and other small and macro-molecules, can be expected to be crucial to formation, regulation and function of RNA foci. As a high-resolution CG model, iConRNA is uniquely positioned to work hand-in-hand with experimental studies to understand the molecular basis of RNA condensation and its role in various biological processes [26].

Our benchmarks also revealed areas for further improvement of iConRNA. In particular, iConRNA only considers the canonical WC base pairs and neglects the formation of non-canonical base pairs [111–113]. It’s a challenge to mimic each kind of base pairing by specific interactions. A simple and practical strategy may be to introduce a general hydrogen bond formation ability into each potential donor and acceptor that could contribute to the formation of non-canonical base pairing. However, it has proven challenging to balance the strength and propensity of different types of base pairing at the CG level. In addition, although the effective magnesium did perform well in the benchmarks summarized in this work, it’s still limited in describing chelated Mg^2+^ binding through the inner sphere. Using the same effective concentration and temperature dependence for a given RNA also neglects the difference in Mg^2+^ binding depending on the local structural environment. Despite these limitations, the successes demonstrated in this work support the great potential of using intermediate resolution CG models for simulating RNA dynamic conformation and phase separation. The current iConRNA model, albeit with limitations, already provides a powerful tool for studying RNA condensates with much greater details compared to the existing 1 or 2-bead models.

## METHODS

### Atomistic simulations

All atomistic simulations were performed using the GROMACS 2022 package [114–116] in the NPT ensemble. The initial structures of RNAs were generated using the Nucleic Acids Builder module available in the AmberTools package [117]. The AMBER99 force field [118] with TIP3P waters [119] was used in combination with the parmbsc0 [120] and the parmchiOL3 [121] corrections for RNA. The ‘microMg’ parameters from Grotz et al [122] were used for Mg^2+^ with the parameters of Cl^-^ from Mamatkulov-Schwierz [123]. Packmol [124] and CHARMM-GUI [125–127] were used to build the initial simulation boxes. Periodic boundary conditions (PBC) and the particle-mesh Ewald (PME) algorithm [128,129] were applied to calculate the long-range electrostatic interactions, while the cut-off of both the short-range electrostatic and vdW potential were set to 1.1 nm. The V-rescale thermostat [130] was used to maintain the temperature with a coupling time constant of 1.0 ps, while the Parrinello-Rahman method [131] was used to maintain the pressure at 1 bar with a coupling time constant of 5.0 ps.

To derive the reference distributions of bonded terms, 500 ns production simulations were performed for ssRNA tetramers after appropriate equilibration. For each bonded term, the distribution was obtained from any tetramer that contain this term. To calculate the concentration and temperature dependence of excess Mg^2+^, a single copy of (CAG)_31_ or (CAG)_10_ in the native hairpin conformation (see **Figure S3**) was placed in a simulation box with different concentrations of MgCl_2_ and NaCl, where MgCl_2_ was used to do neutralization. The box size is 20×20×20 nm^3^ for (CAG)_31_ or 10×10×10 nm^3^ for (CAG)_10_. For (CAG)_10_, six 100-ns simulations were performed at every 10 K increment from 303 K to 353 K with 25 mM NaCl and 50 mM MgCl_2_, to derive the temperature dependence of excess Mg^2+^. For (CAG)_31_, four 100-ns simulations were performed to predict the concentration dependence of excess Mg^2+^, which was then used to derive *F*_Mg_ and *M*_1/2_ by fitting to the Hill equation (Eqn. 7). The number of Mg^2+^ (*N*_Mg_) in excess to the bulk around the RNA was calculated as [103,132,133]

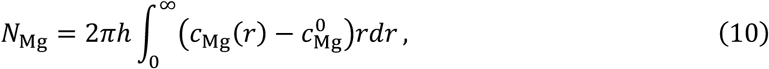

where *h* is the dimension of RNA along the axis direction, and 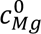 is the bulk concentration.

### CG simulation protocols

All CG simulations were performed using OpenMM 8.1.0 [134] in the NVT ensemble. Langevin dynamics was used with a friction coefficient of 0.1 ps^-1^. In-house Python packages were used to create initial CG RNA structures or convert atomistic models into iConRNA representation. For all single-chain simulations, RNA was centered in the box sized from 20 nm to 50 nm for different RNAs and then simulated for at least 2 μs. For the simulations of structured RNAs, two replica simulations were performed starting from either disordered chains or structured chains. At least 2-μs simulations were run to obtain the equilibrium structures. For phase separation simulations, multi-copies of RNAs were randomly packed into cubic boxes to build the initial solution with specific RNA concentrations. The detailed numbers of copies and box sizes are summarized in **Table S4**. To build the phase diagrams, 2 replicas of at least 2-μs simulations were performed at every 10 K increment from 303 K to 353 K for each RNA and/or Mg^2+^ concentrations.

### Heat capacity calculation

Following the framework presented by Walker et al. [100], we first performed replica-exchange MD simulations with 15 replicas, where the temperatures were distributed exponentially from 273 K to 393 K. Langevin integrator with a collision frequency of 5 ps^-1^ and time step of 8 fs was used. Replica exchange moves were attempted every 200 MD time steps and a total of 300 ns per replica was run. The constant volume heat capacity was computed from the mean square fluctuation of the enthalpy using the multistate Bennett acceptance ratio (MBAR) method as implemented in the pymbar package [135],

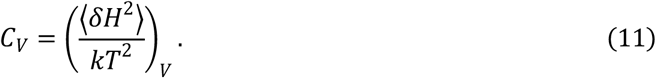

Sequences of all RNAs simulated in this work are provided in **Table S5**.

## SUPPLEMENTARY DATA

The data underlying this article including all iConRNA force field files are available freely from GitHub at: https://github.com/lslumass/iConRNA.

Supplementary Data are available online through https//xxx.

## ACKNOWLEDGEMENTS

All simulations were performed on the Pikes GPU cluster housed in the Massachusetts Green High-Performance Computing Cluster (MGHPCC).

## Supplementary Information

### Supplementary Tables

**Table S1.**
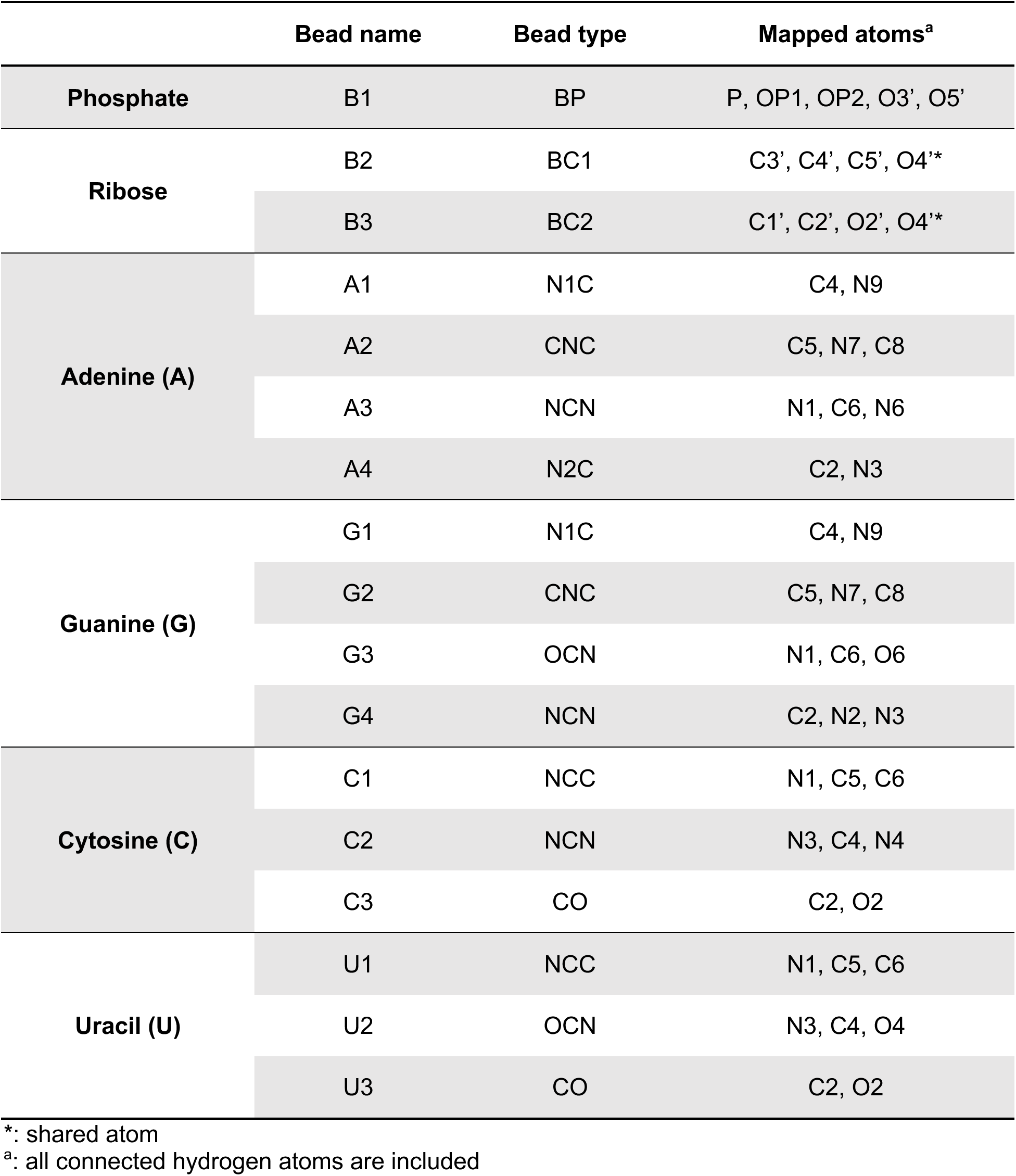
Atomistic to CG mapping scheme (also see Figure 1)

**Table S2.**
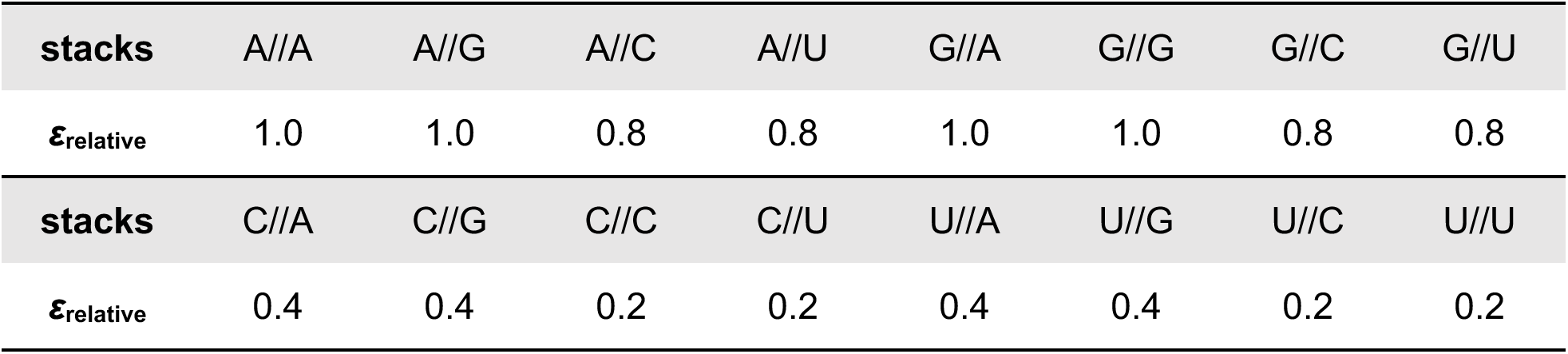
Relative stacking strength to A//A stacking. In the final model, *ε_A//A_* = 2.05 kcal/mol (see main text).

**Table S3.**
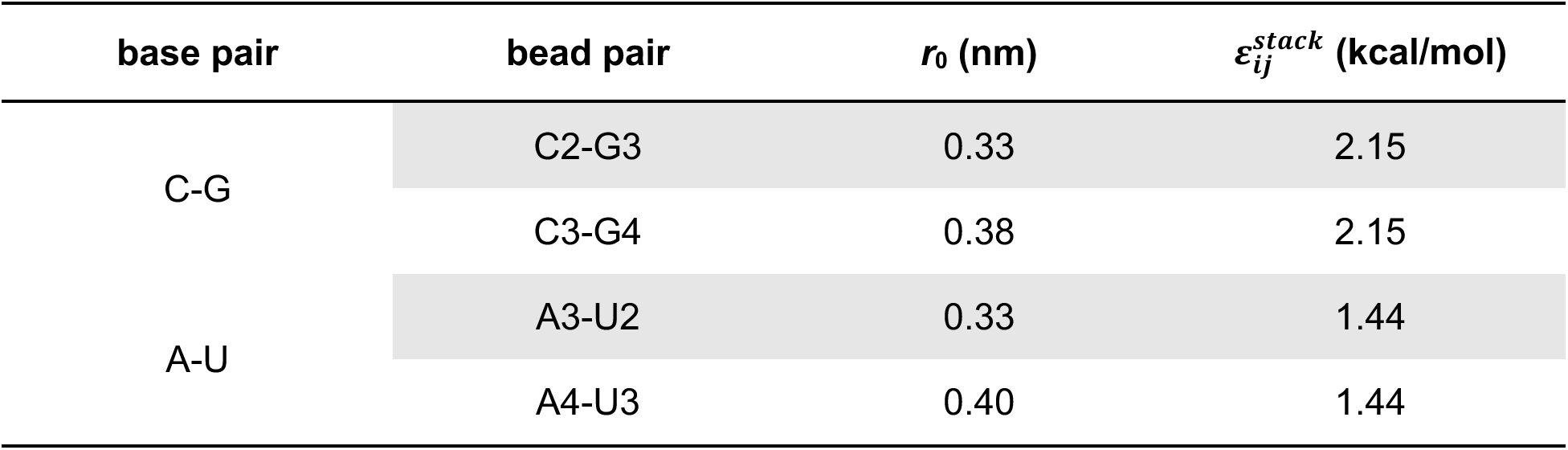
Equilibrium distance for different hydrogen bonding CG beads and the optimized hydrogen bonding pair strength (see main text).

**Table S4.**
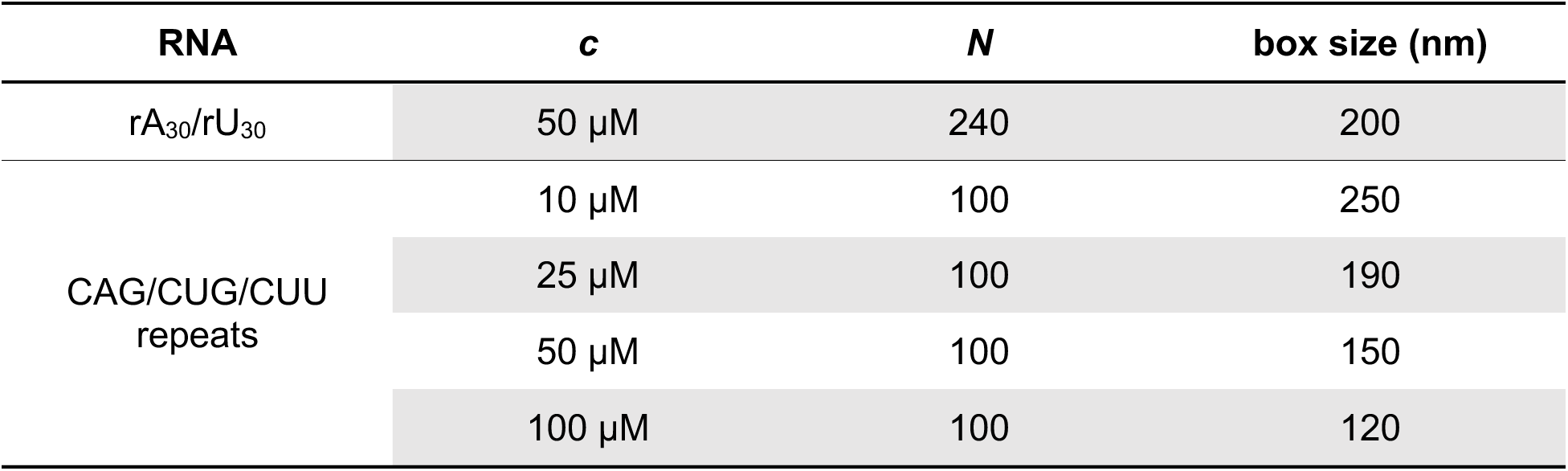
RNA concentrations (*c*), RNA chain numbers (*N*), and box sizes for the simulations of phase separation.

**Table S5.**
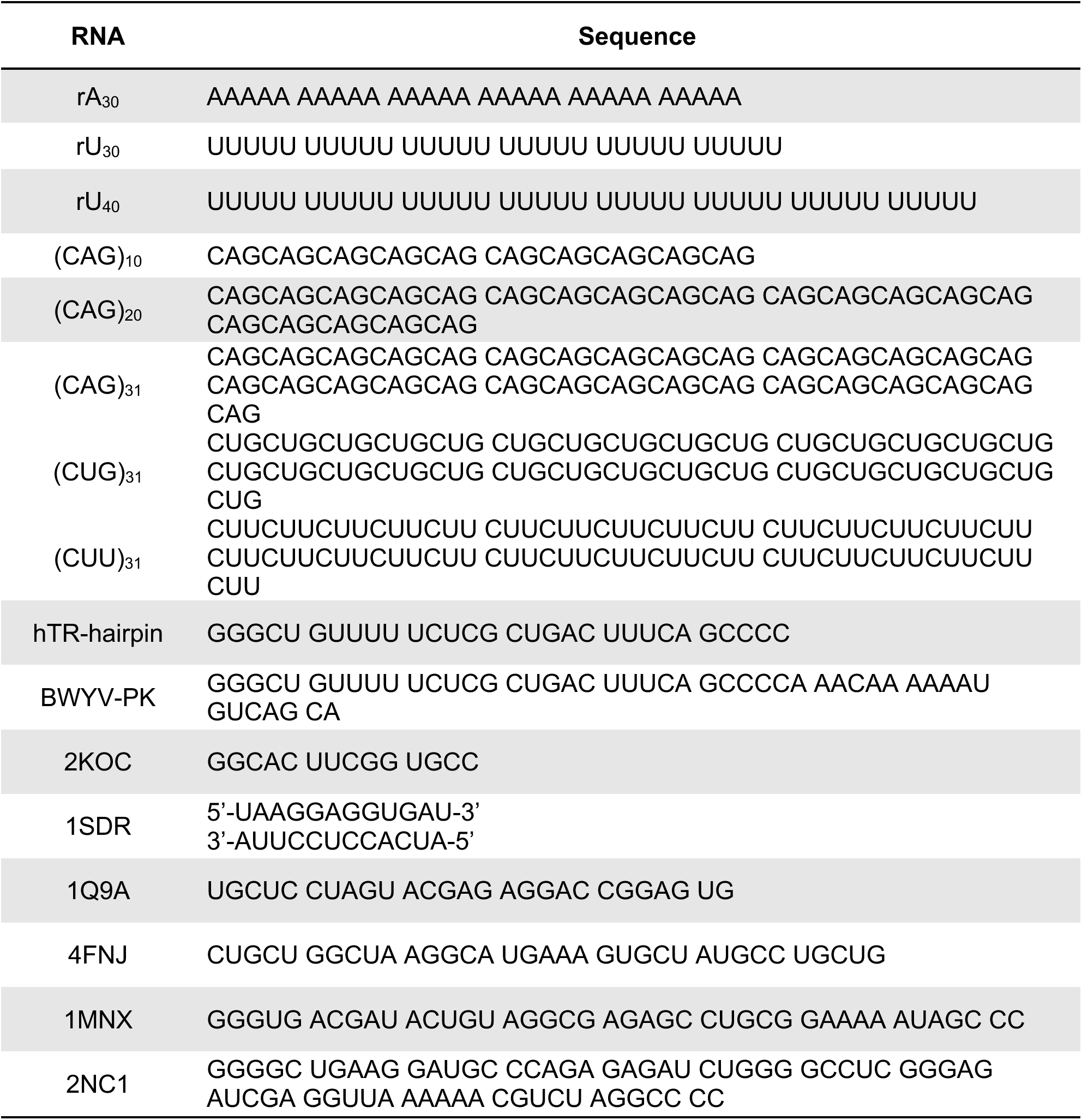
The sequences of all RNAs simulated in this work (except RNA tetramers used for deriving bonded parameters, which are provided in the main text).

**Table S6.**
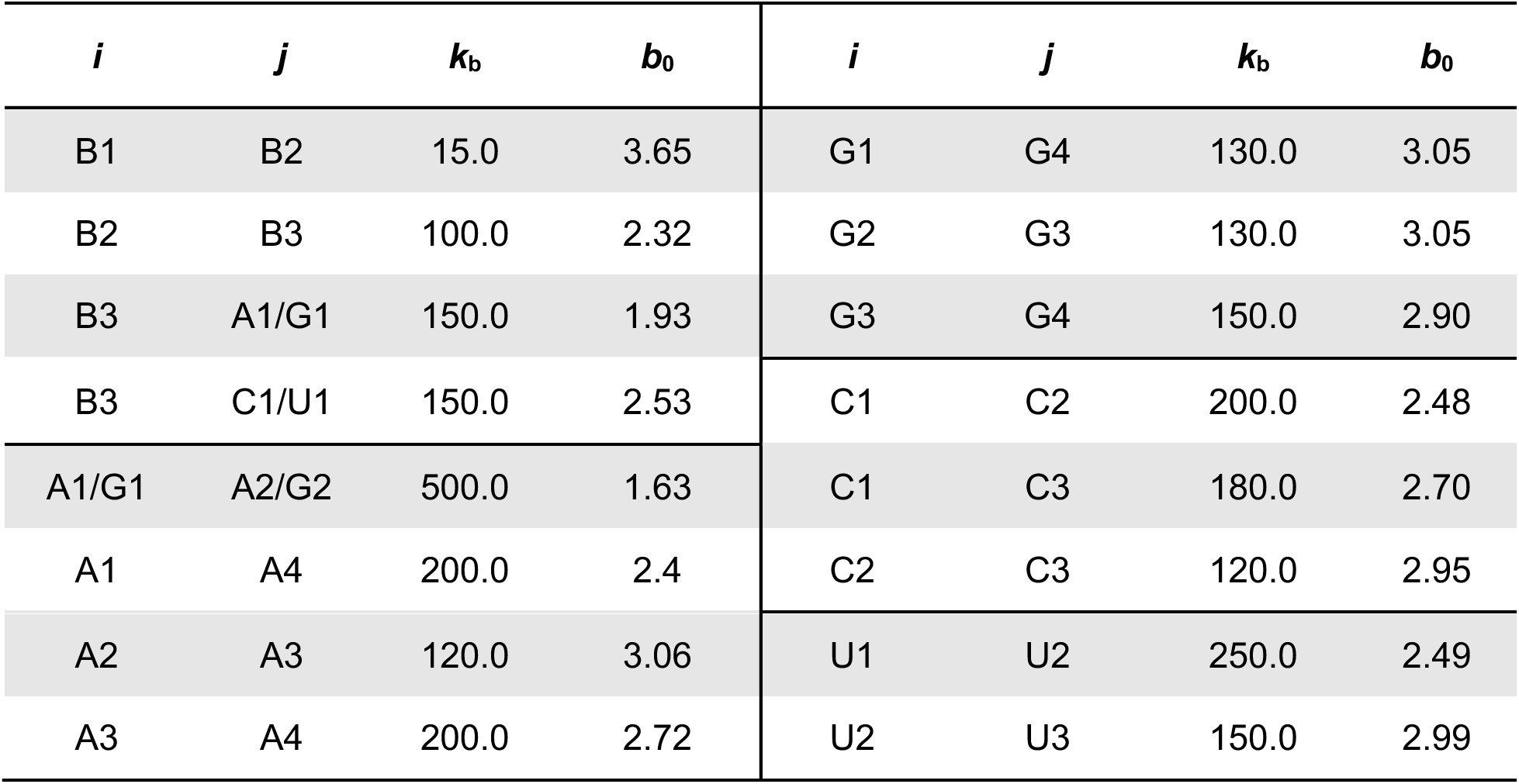
Bond parameters, *k*_b_ in kcal/(mol‧Å^2^) and *b*_0_ in Å.

**Table S7.**
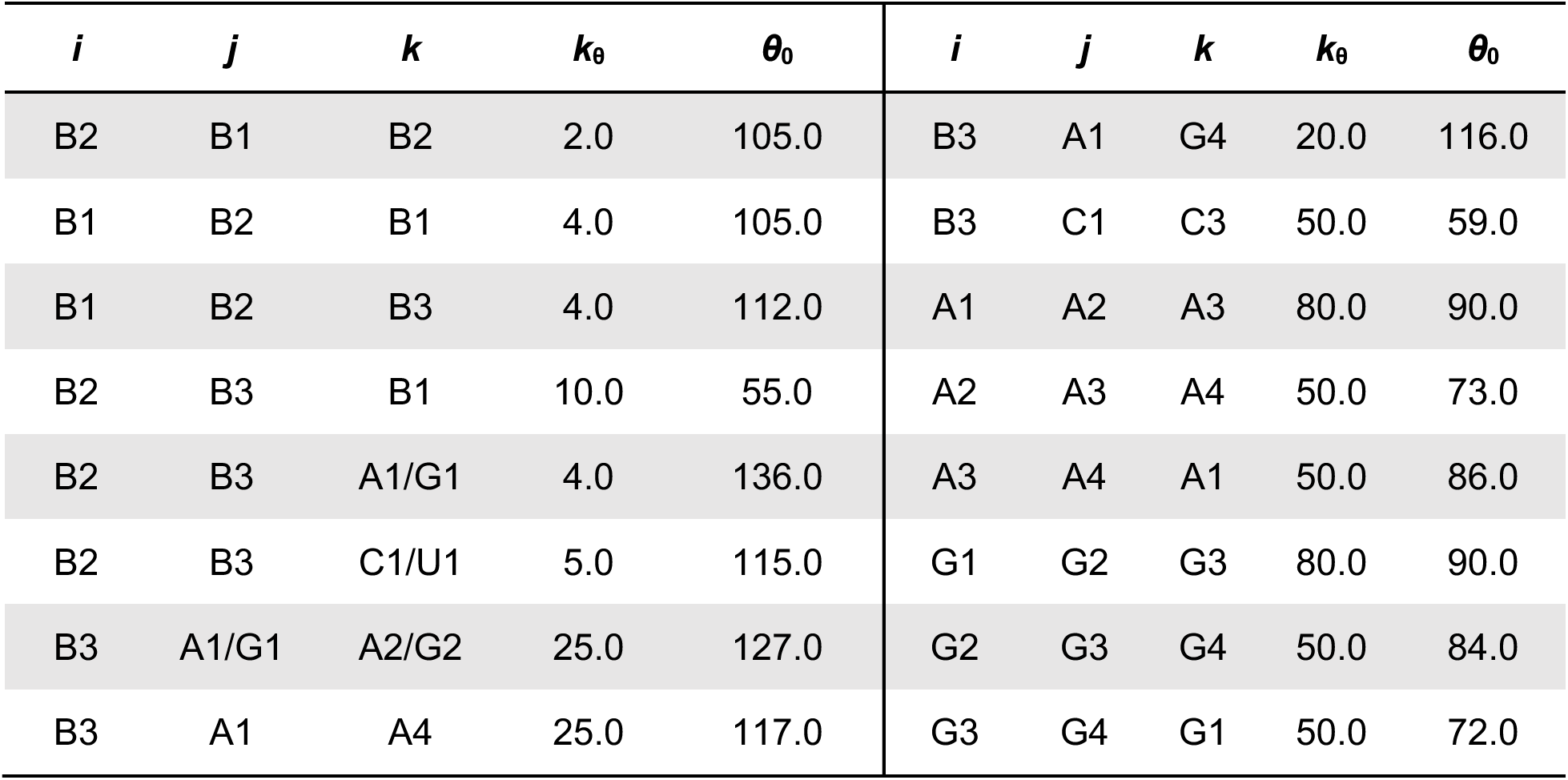
Angle parameters, *k*_θ_ in kcal/(mol‧rad^2^) and *θ*_0_ in degree.

**Table S8.**
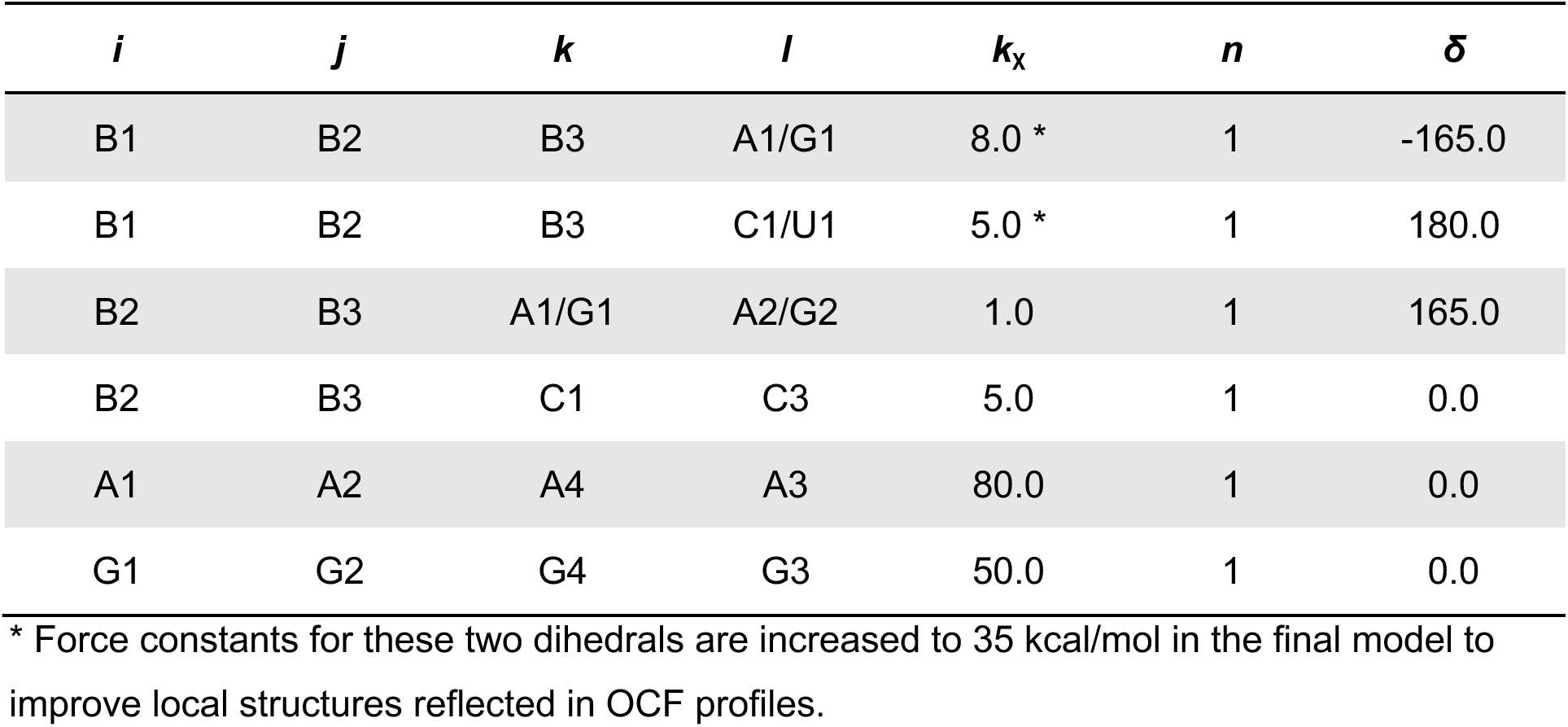
Dihedral angle parameters, *k*_χ_ in kcal/mol and *δ* in degree.

**Table S9.**
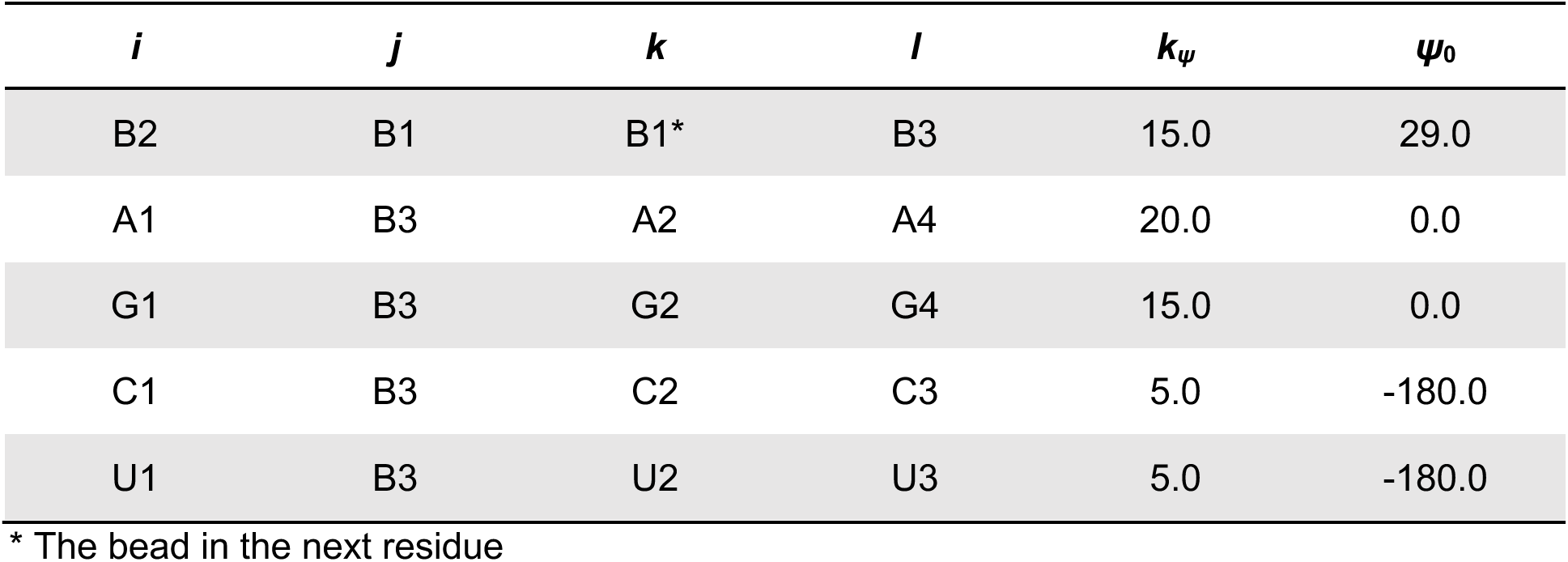
Improper dihedral parameters, *k*_ψ_ in kcal/(mol‧rad^2^) and *ψ_0_* in degree.

**Table S10.**
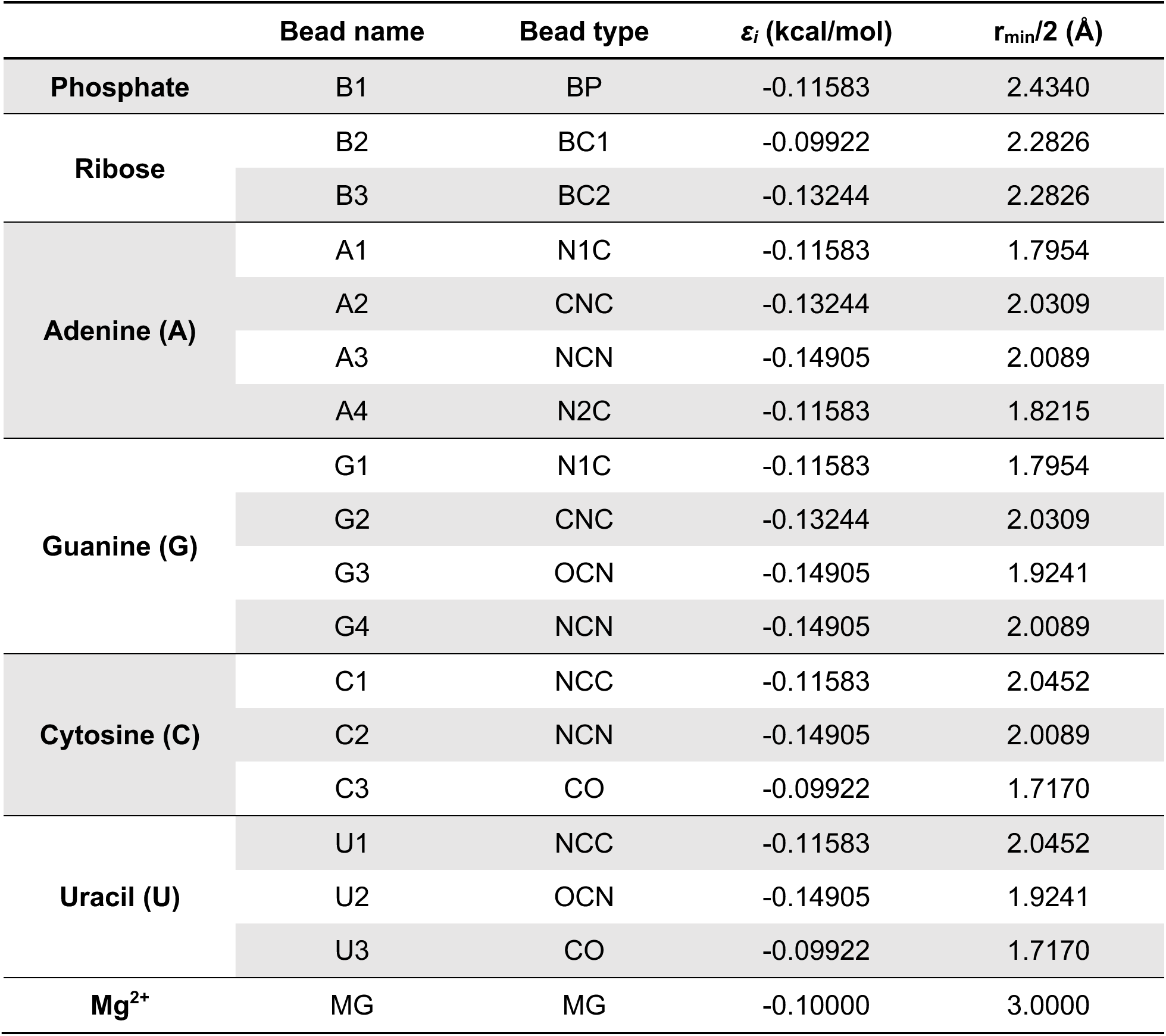
Final vdW parameters of all CG bead types.

**Table S11.**
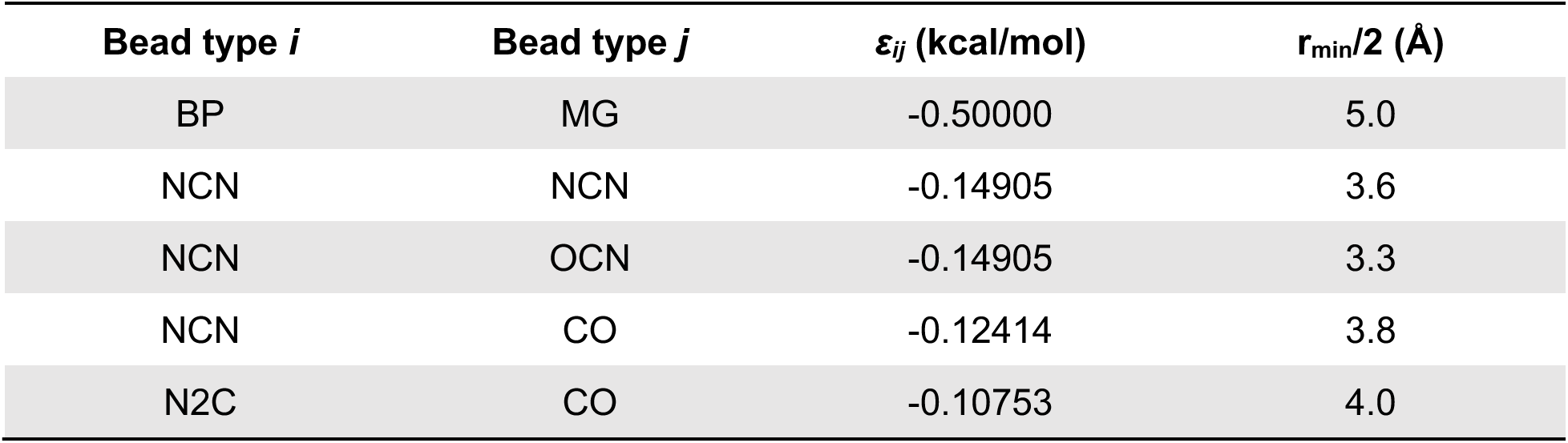
NBFIX.

**Table S12.**
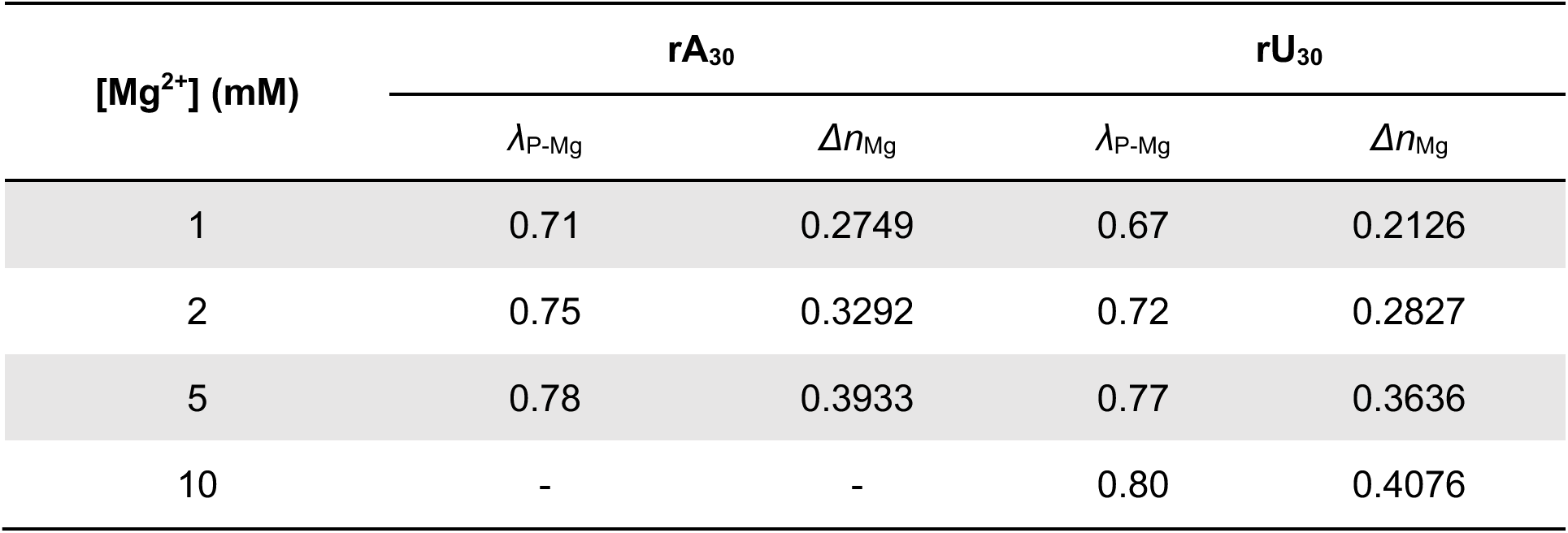
***λ*_P-Mg_ and *Δn*_Mg_ of rA_30_ and rU_30_ at different concentration of Mg^2+^**

**Table S13.**
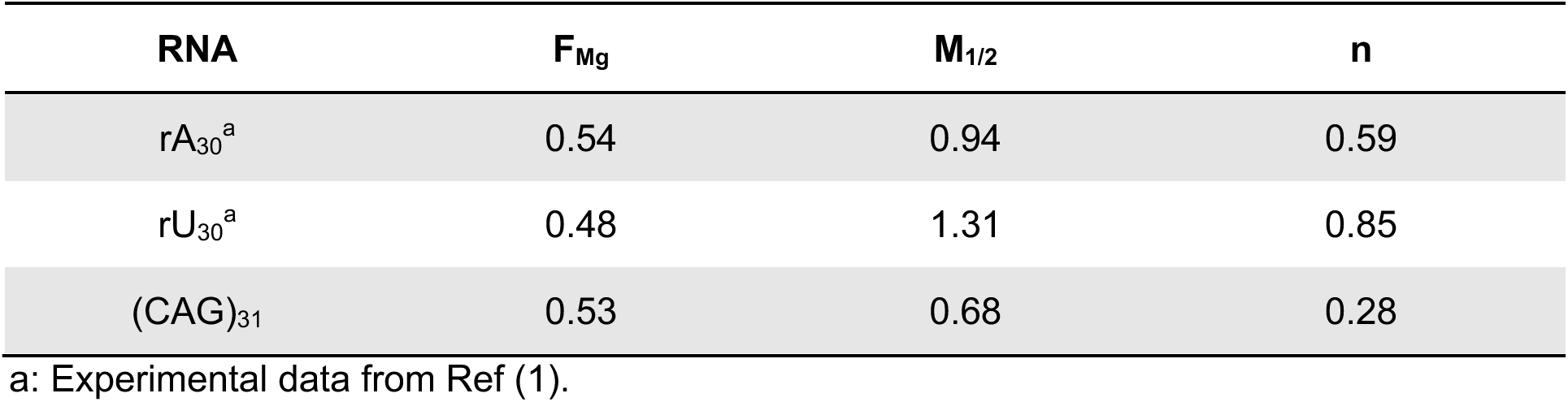
Hill fit parameters for rA_30_, rU_30_ and (CAG)_31_.

### Supplementary Figures

**Figure S1.**
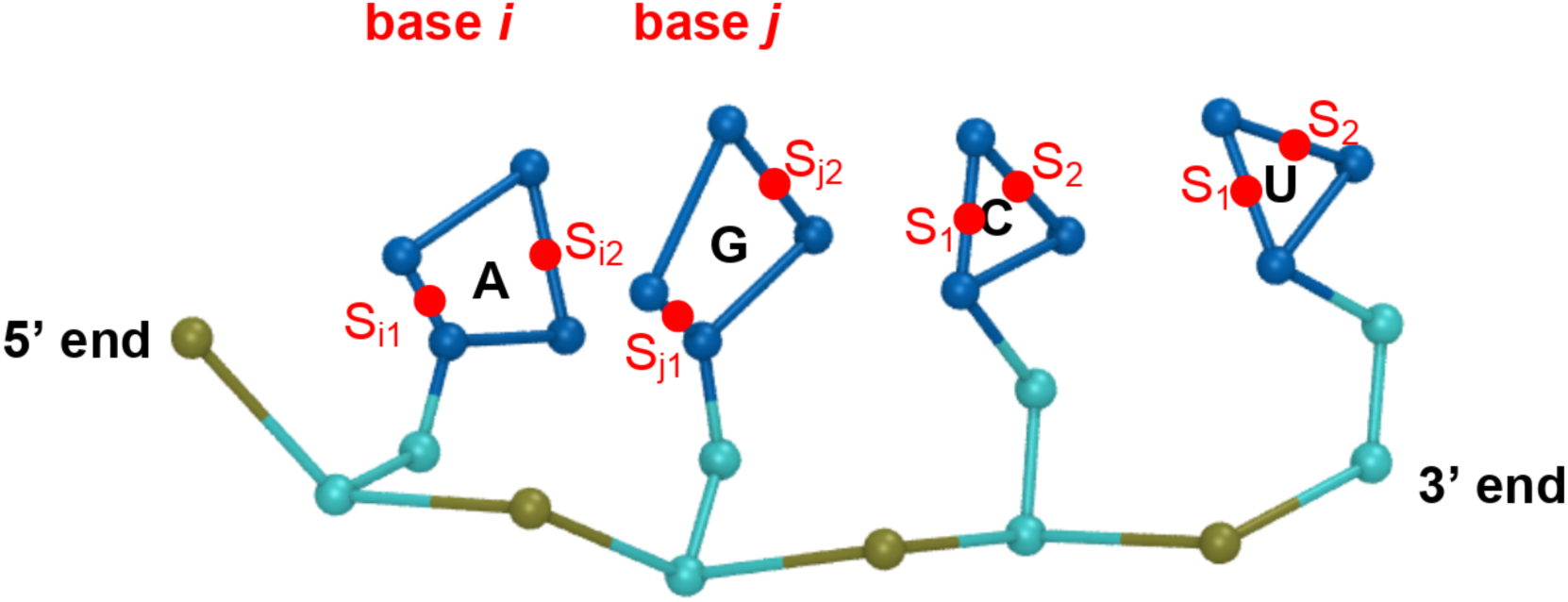
Scheme of the assignment of base stacking. Red points S_i1_ and S_i2_ are the two virtual sites of base *i*. From the 5’ end to the 3’ end, the stacking interaction is applied between S_i2_ and S_j1_.

**Figure S2.**
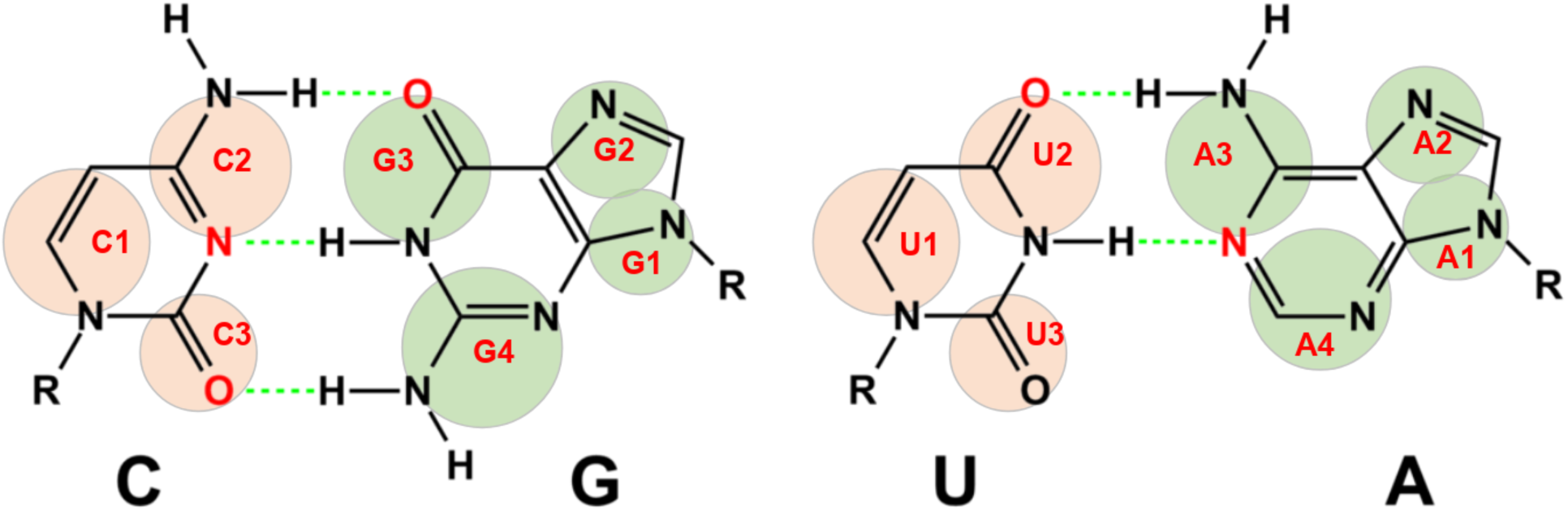
Scheme of A-U and G-C base pairing. Related bead names are in red fonts. The green dashed lines show the hydrogen bonds for each pair.

**Figure S3.**
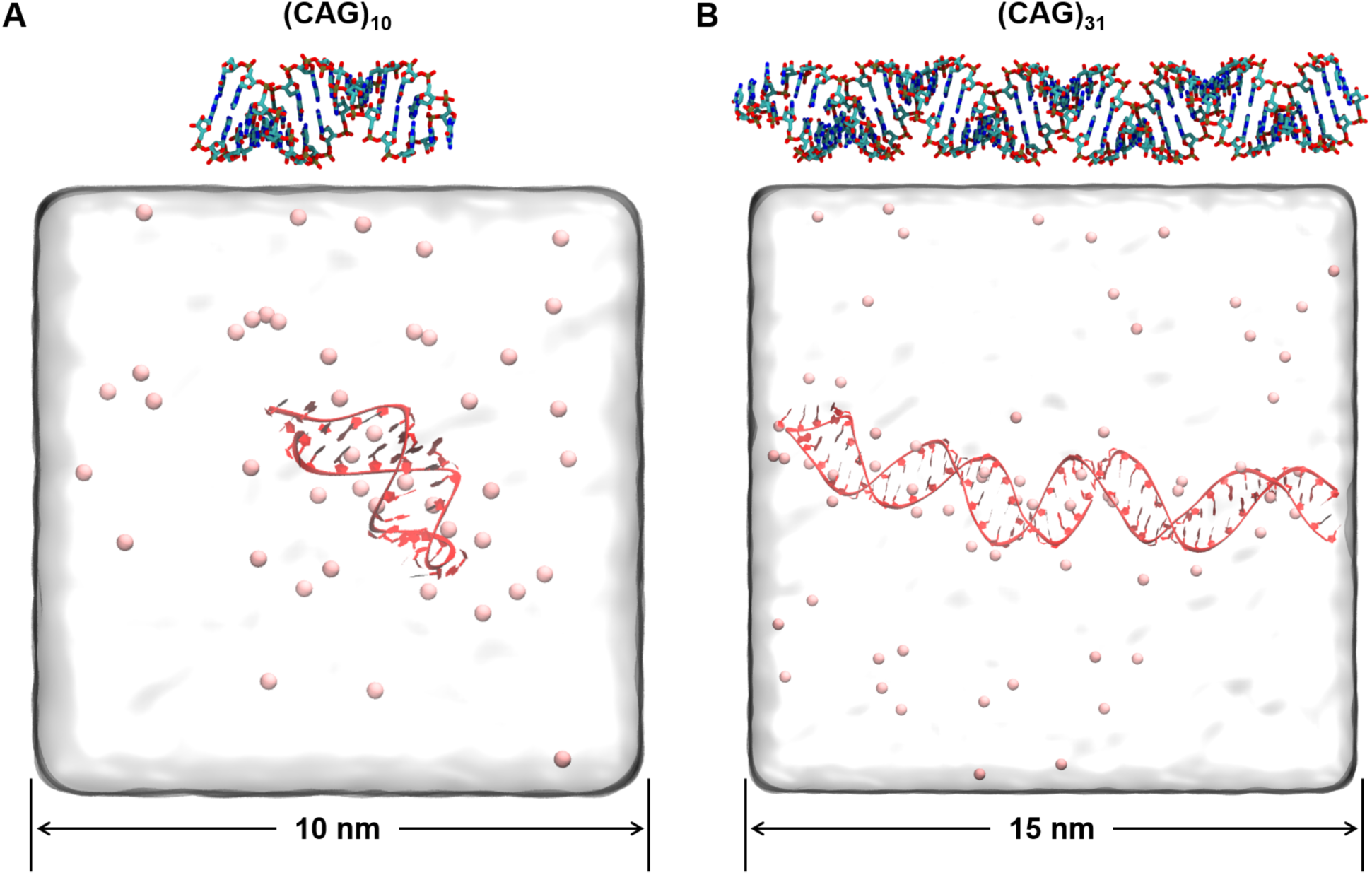
The ideal hairpin structures and equilibrated states with Mg^2+^ ions (pink beads) of (CAG)_10_ (**A**) and (CAG)_31_ (**B**).

**Figure S4.**
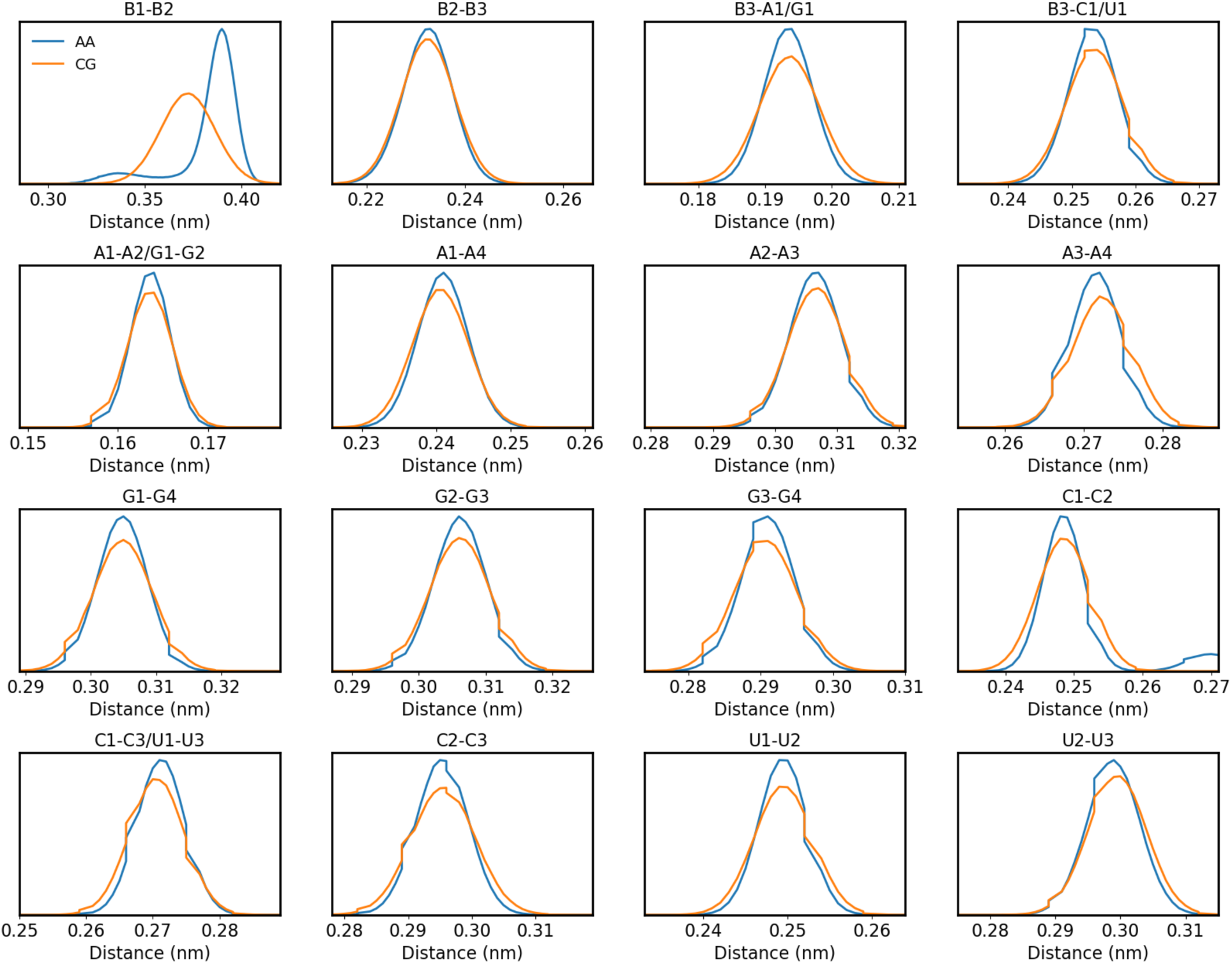
AA and CG distributions of bonds. The average distributions from the AMBER AA simulations are shown in blue, whereas the average CG distributions are shown in orange. The title of each subplot lists the bead names that participate in the bond.

**Figure S5.**
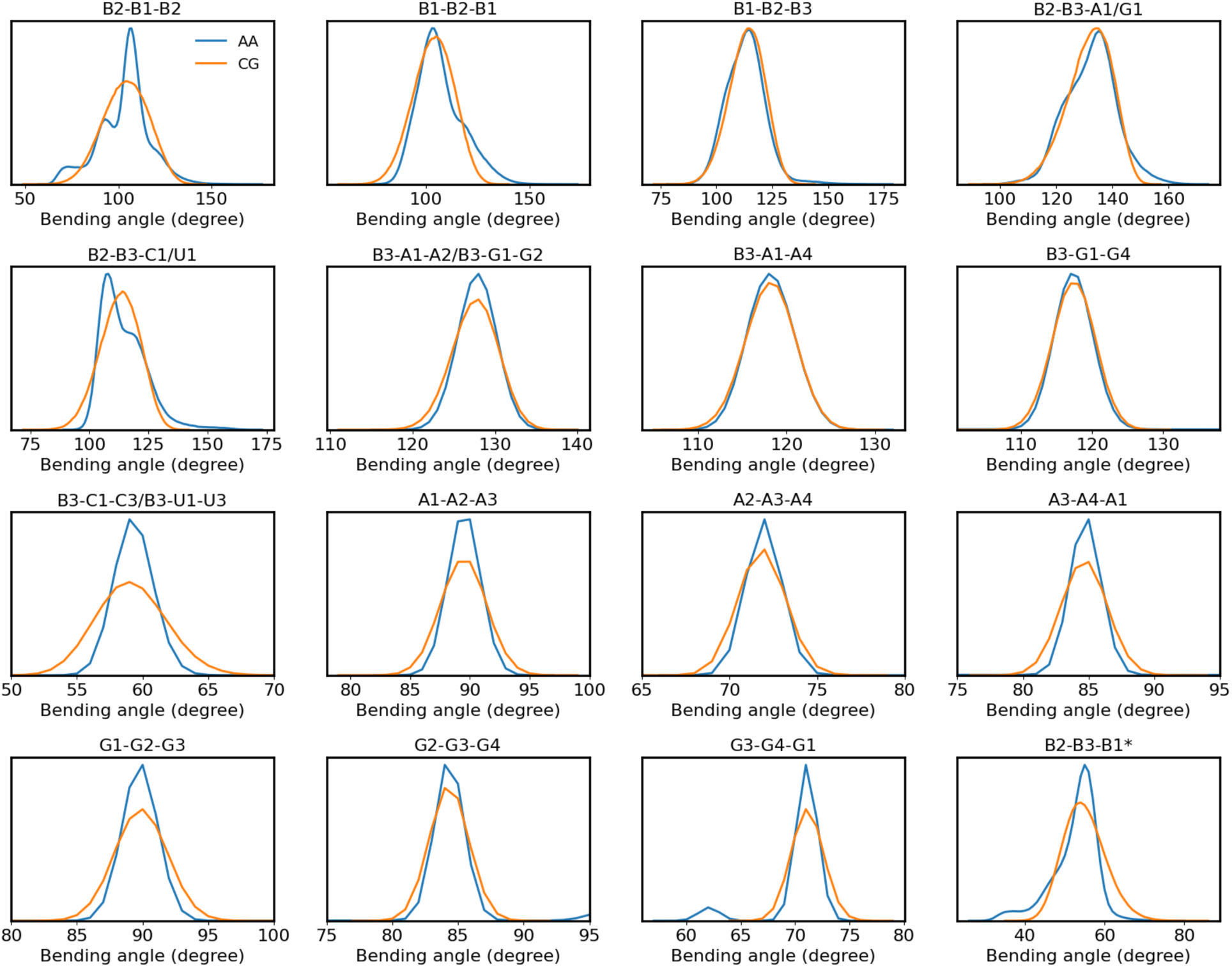
AA and CG distributions of angles. The average distribution from the AMBER AA simulations is shown in blue, whereas the average CG distributions are shown in orange. The title of each subplot lists the bead names that participate in the angle following the same order as the angle.

**Figure S6.**
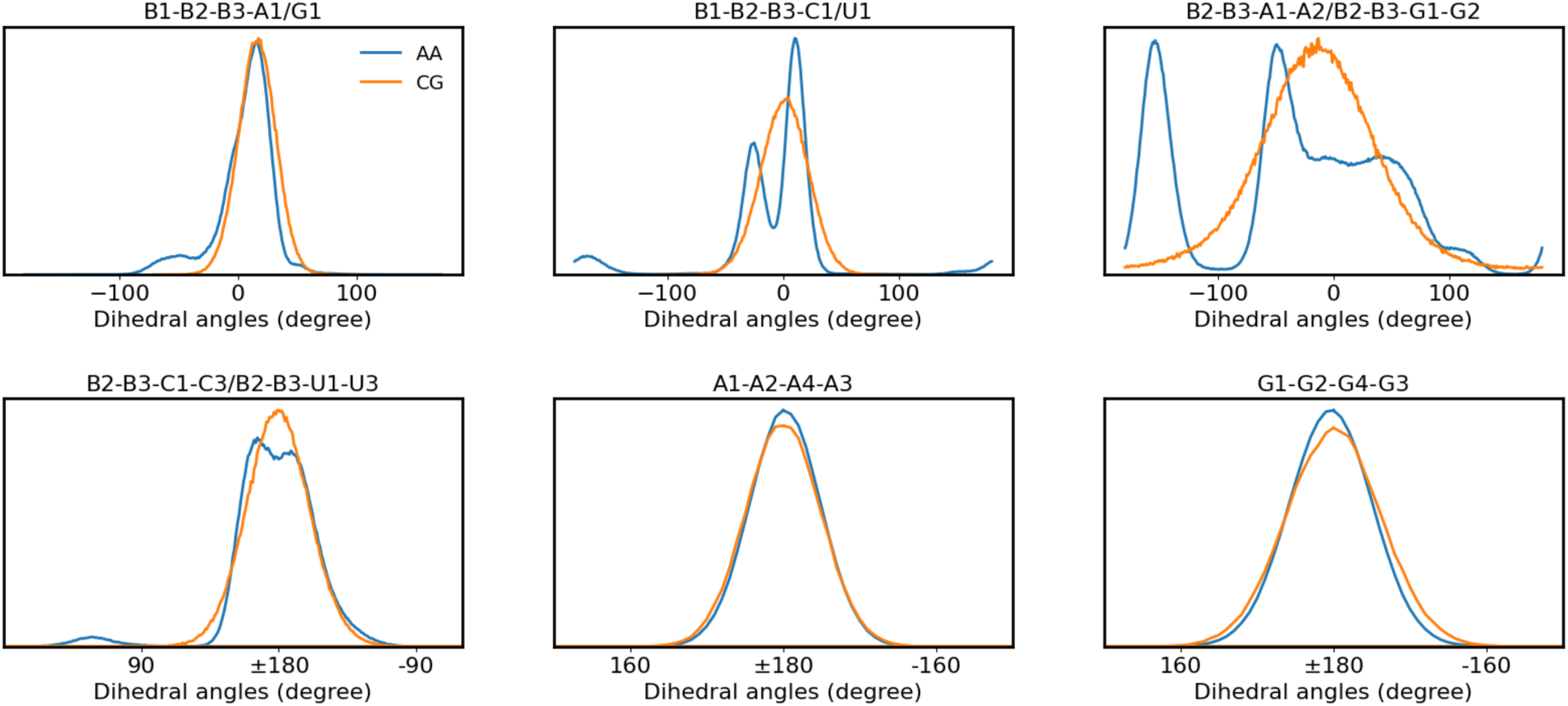
AA and CG distributions of dihedral angles. The average distributions from the Amber AA simulation are shown in blue, whereas the average CG distributions are shown in orange. The title of each subplot lists the bead names that participate in the angle following the same order as the dihedral angles.

**Figure S7.**
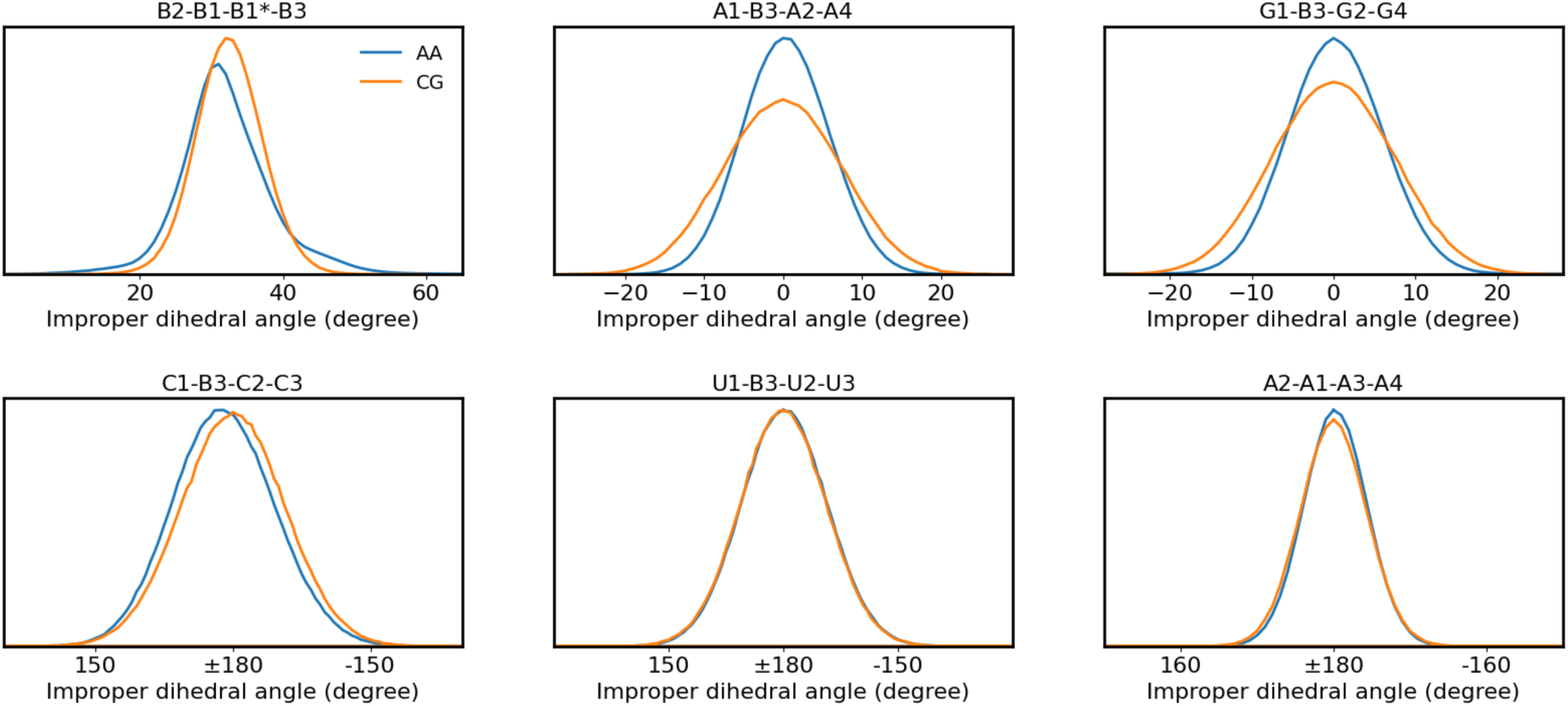
AA and CG distributions of dihedral angles and improper dihedral angles. The average distributions from the AMBER AA simulations are shown in blue, whereas the average CG distributions are shown in orange. The title of each subplot lists the bead names that participate in the angle following the same order as the dihedral angles. B1* denotes the B1 bead of the next residue.

**Figure S8.**
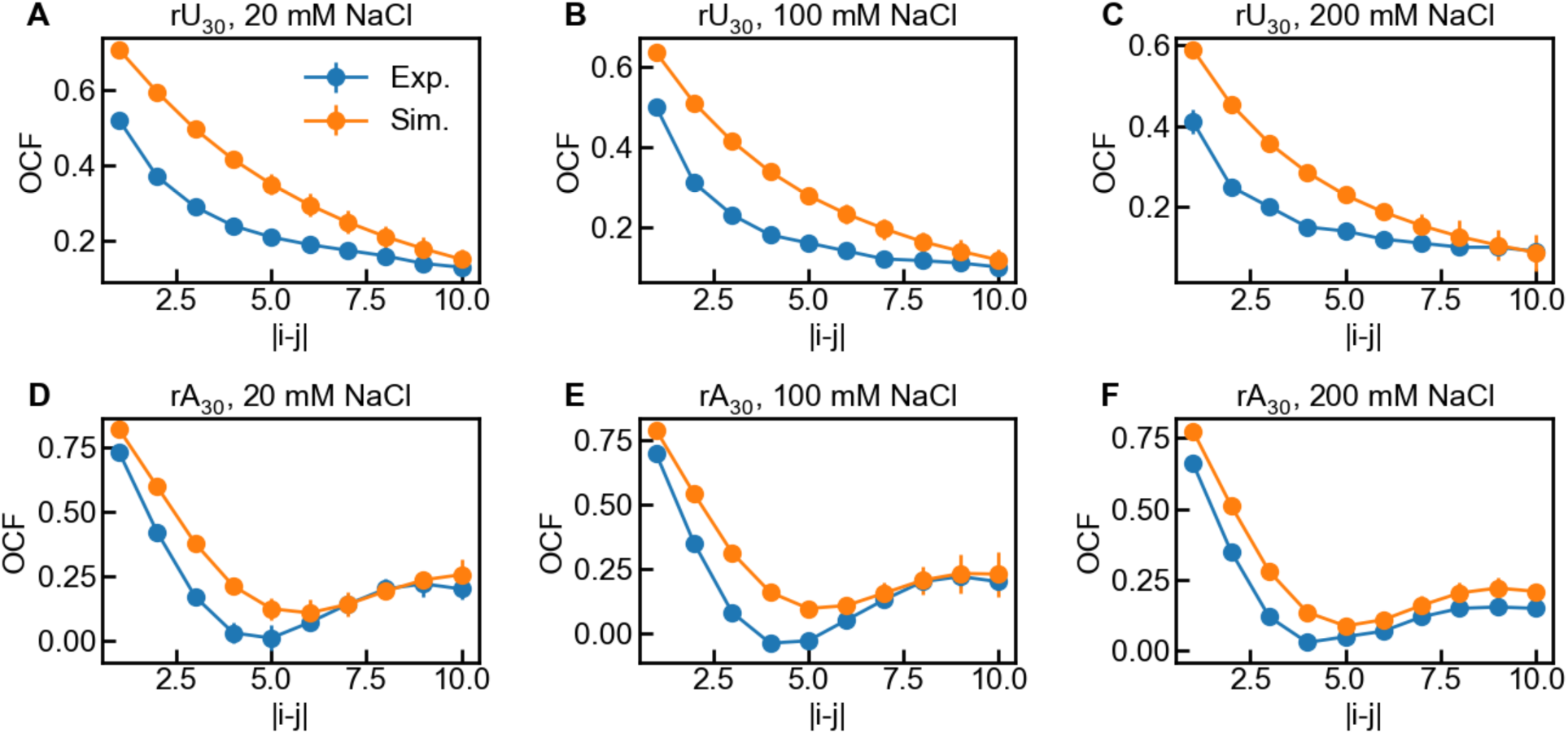
Orientation correlation function (OCF) of rU_30_ (**A-C**) and rA_30_ (**D-F**). The NaCl concentrations were set as 20 mM, 100 mM, and 200 mM to show the salt dependence of OCF. |*i*-*j*| is the difference in residue numbers between the *i*th and *j*th phosphate-phosphate bond vectors (Eqn. 6 in main text). Experimental results are shown in blue whereas simulation results are in orange. The error bars of simulation results were estimated from block analysis.

**Figure S9.**
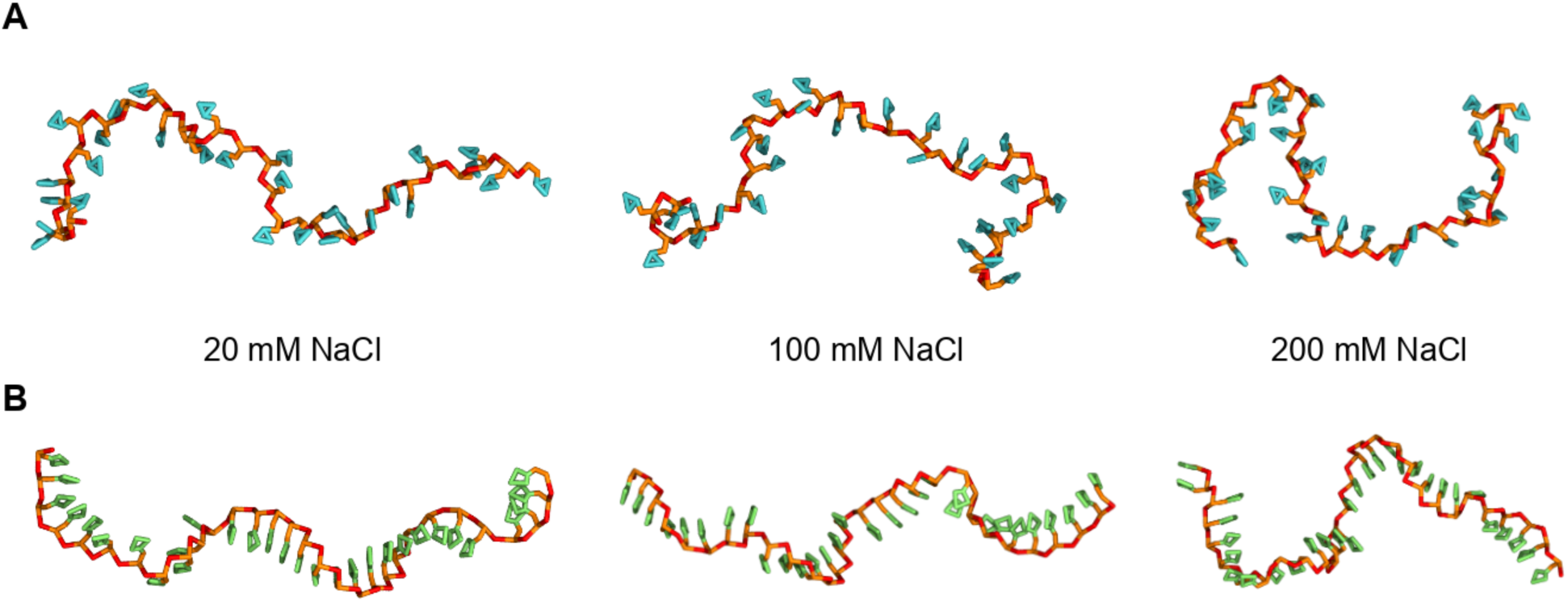
Representative structures of (**A**) rU_30_ and (**B**) rA_30_ at 20 mM, 100 mM, and 200 mM NaCl solution. Phosphate beads B1 are shown in red, while B2 and B3 for ribose are in orange. Bases U and A are in cyan and lime, respectively.

**Figure S10.**
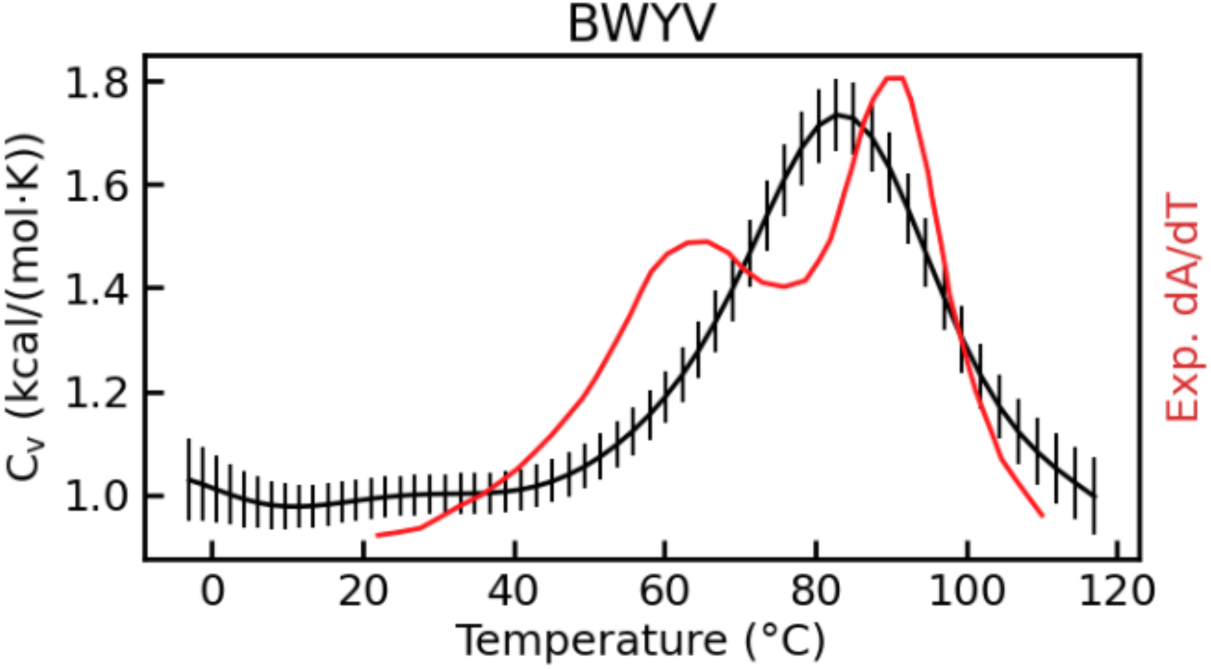
Heat capacity (C_v_) of Beet Western Yellow Virus (BWYV) pseudoknot. Experimental data is the red line, where the first derivative of UV-absorbance with respect to temperature (dA/dT) at 280 nm is used as the reference melting profile (2).

**Figure S11.**
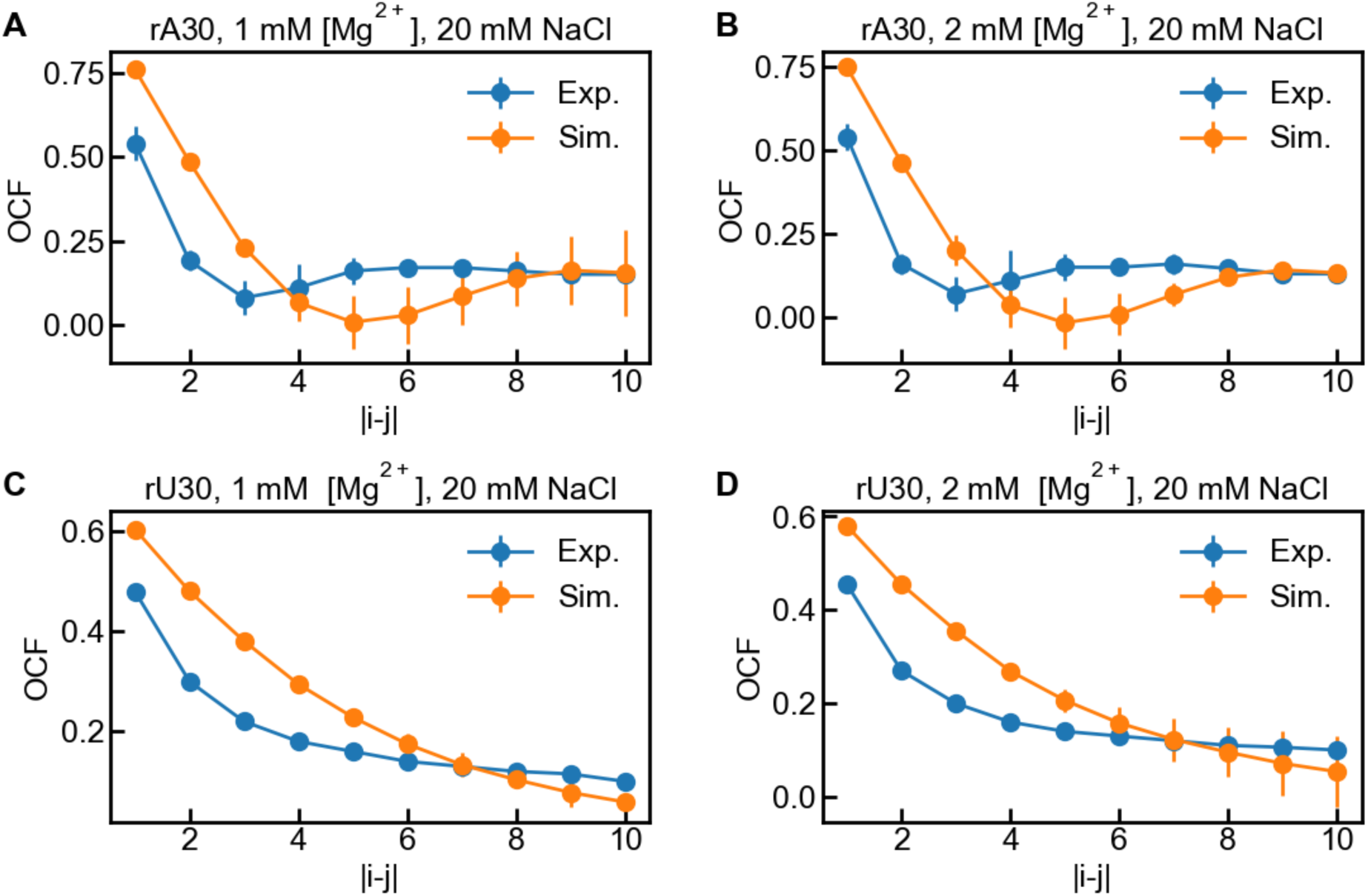
OCFs of rA_30_ (**A, B**) and rU_30_ (**C, D**) at different ion conditions. |*i*-*j*| is the difference in residue numbers between the *i*th and *j*th phosphate-phosphate bond vectors (Eqn. 6 in main text). The experimental results are shown in blue whereas CG simulation results are in orange. The error bars of simulation results were estimated from block analysis.

**Figure S12.**
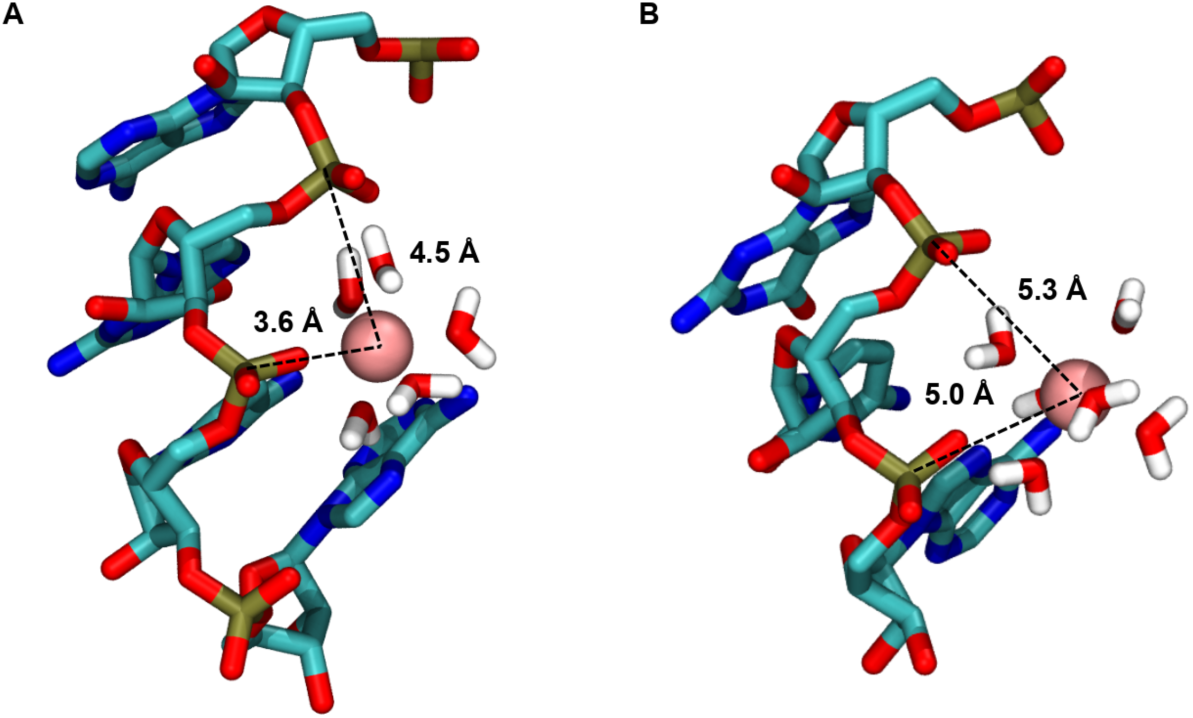
Different binding models of Mg^2+^ with phosphate groups. (**A**) one inner-sphere binding with a distance of 3.6 Å and one outer-sphere binding with a distance of 4.5 Å. (**B**) Two outer-sphere binding with distances of 5.0 and 5.3 Å. A standard colour scheme is used for elements, and Mg^2+^ is shown in a pink vdW sphere.

**Figure S13.**
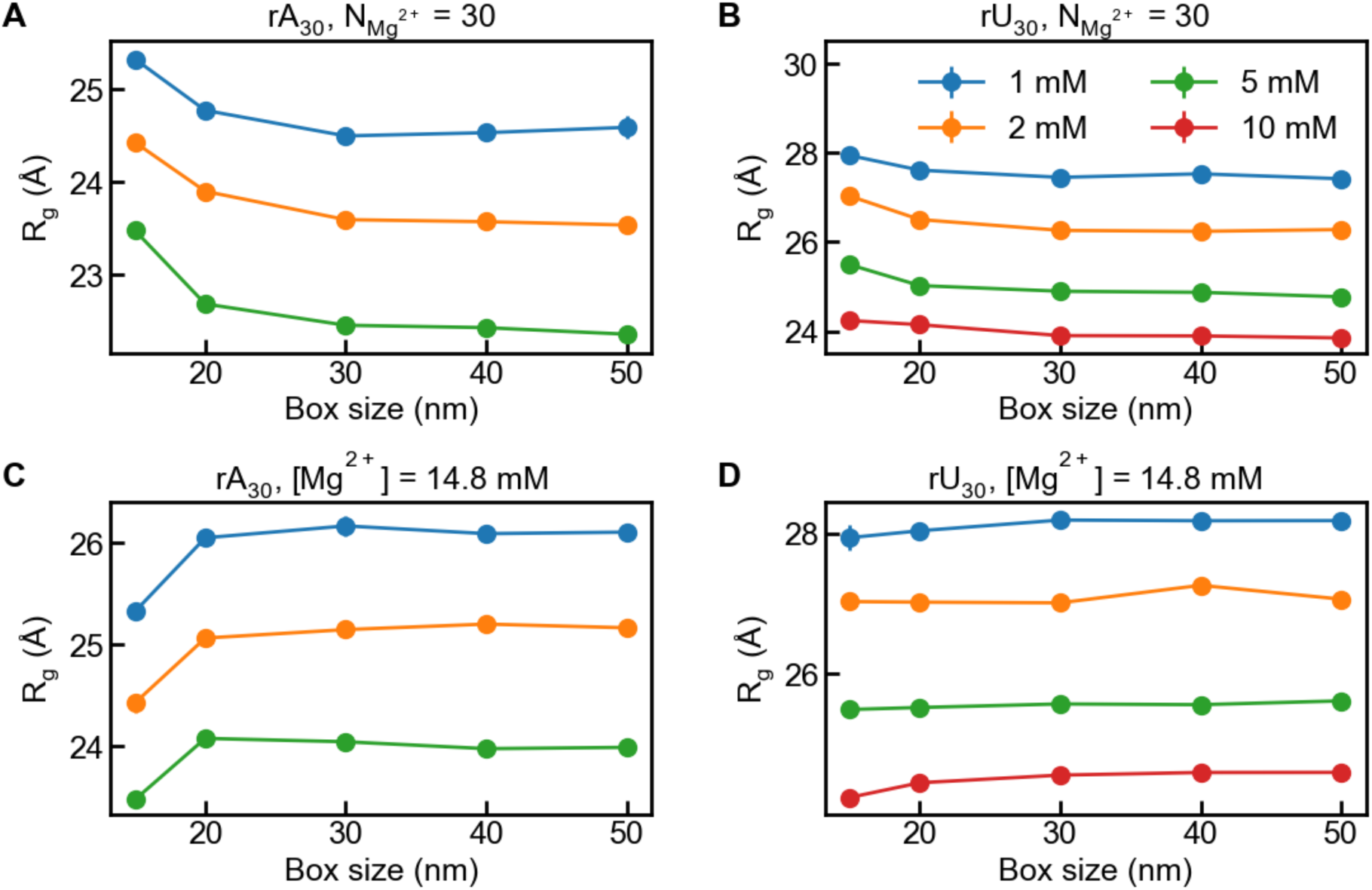
Effect of the simulation box size on *R*_g_ with fixed number of Mg^2+^ (**A, B**) and fixed concentration of Mg^2+^ (**C, D**). Panels A and C are for rA_30_, while panels B and D are for rU_30_.

**Figure S14.**
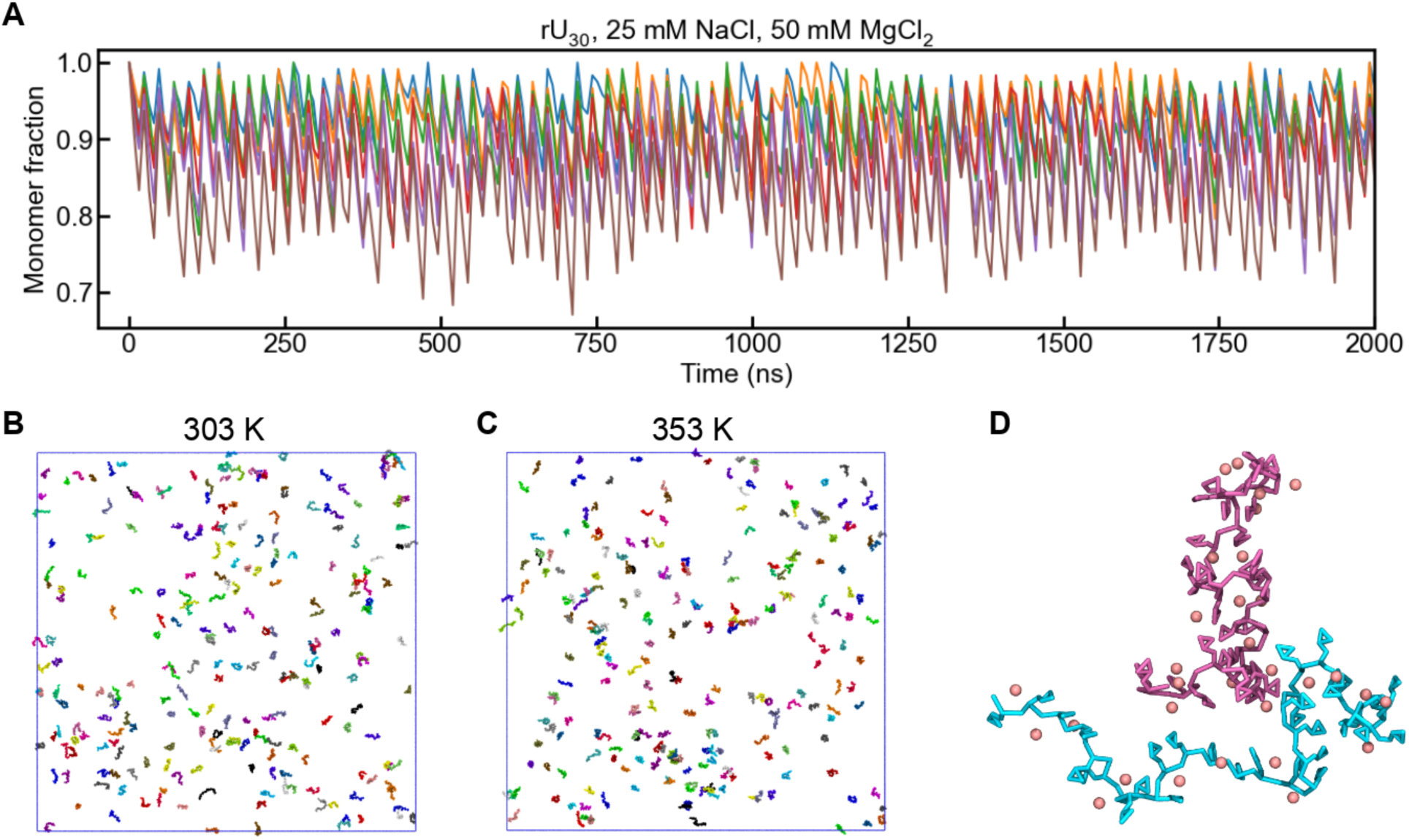
Phase separation simulations of rU_30_. (**A**) Monomer fractions at 303, 313, 323, 333, 343, and 353 K under 25 mM NaCl and 50 mM MgCl_2_. The initial RNA concentration is 50 µM. Note that the system remains dispersed at all temperatures. (**B, C**) Snapshots at 303 and 353 K. Mg^2+^ ions were omitted for clarity. (**D**) A representative snapshot of rU_30_ dimer, with Mg^2+^ ions shown as pink beads.

**Figure S15.**
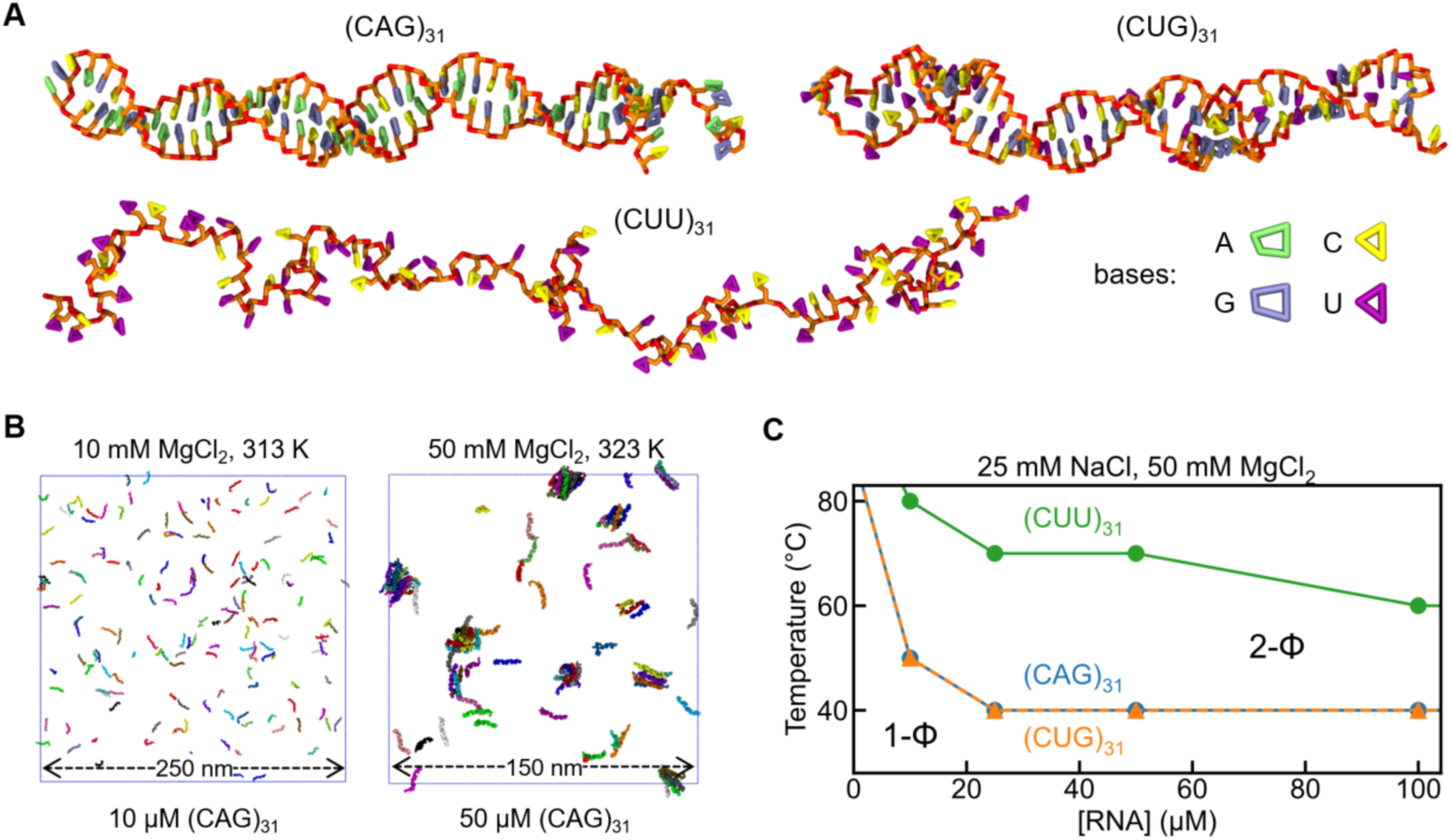
Phase separation of RNA triplet repeats. (**A**) Snapshots of representative structures of monomeric (CAG)_31_, (CUG)_31_, and (CUU)_31_ under 25 mM NaCl at 303 K. Phosphate bead B1 and ribose beads B2/B3 are in red and orange, while bases A, G, C, and U are in lime, ice blue, yellow, and purple, respectively. (**B**) Snapshots of representative points in 1-phase (1-Φ, left panel) and 2-phase (2-Φ, right panel) regions, which were obtained from simulations of 10 μM (CAG)_31_ at 313 K and 50 μM (CAG)_31_ at 323 K, respectively (with 25 mM NaCl and 10 mM MgCl_2_). RNAs are coloured in chains. Mg were omitted for clarity. (**C**) Phase diagrams of (CAG)_31_ (blue), (CUG)_31_ (orange), and (CUU)_31_ (green) under 25 mM NaCl and 50 mM MgCl_2_, where transition temperatures are plotted as a function of [RNA]. For each RNA, the lower left area of the boundary line is the 1-Φ region, while the upper right one is the 2-Φ region.

**Figure S16.**
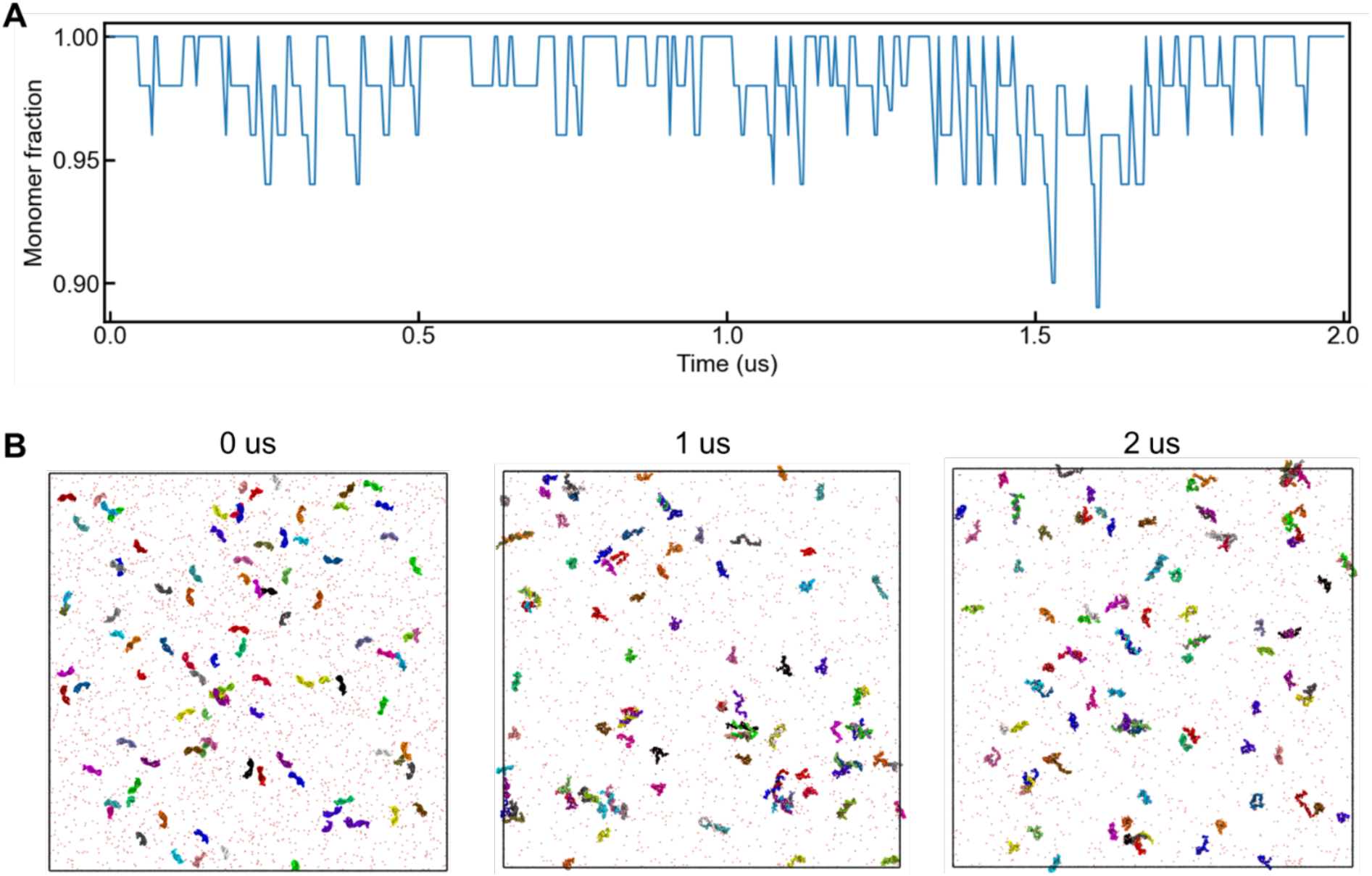
Phase separation simulations of 100 µM (CAG)_10_ under 25 mM NaCl and 50 mM MgCl_2_ at 353 K. (**A**) Monomer fraction as a function of simulation time. (**B**) Simulation snapshots at 0, 1, and 2 μs.

## REFERENCES

1. Brangwynne, C. P., Eckmann, C. R., Courson, D. S., Rybarska, A., Hoege, C., Gharakhani, J., Julicher, F., and Hyman, A. A., Germline P granules are liquid droplets that localize by controlled dissolution/condensation. Science 324 (5935), 1729 (2009).

2. Banani, Salman F., Lee, Hyun O., Hyman, Anthony A., and Rosen, Michael K., Biomolecular condensates: organizers of cellular biochemistry. Nat. Rev. Mol. Cell Biol. 18 (5), 285 (2017).

3. Brangwynne, Clifford P., Tompa, Peter, and Pappu, Rohit V., Polymer physics of intracellular phase transitions. Nat. Phys. 11 (11), 899 (2015).

4. Holehouse, Alex S. and Pappu, Rohit V., Functional Implications of Intracellular Phase Transitions. Biochemistry 57 (17), 2415 (2018).

5. Alberti, Simon, Gladfelter, Amy, and Mittag, Tanja, Considerations and Challenges in Studying Liquid-Liquid Phase Separation and Biomolecular Condensates. Cell 176 (3), 419 (2019).

6. Alberti, Simon and Dormann, Dorothee, Liquid–Liquid Phase Separation in Disease. Annu. Rev. Genet. 53 (1), 171 (2019).

7. Zbinden, Aurélie, Pérez-Berlanga, Manuela, De Rossi, Pierre, and Polymenidou, Magdalini, Phase Separation and Neurodegenerative Diseases: A Disturbance in the Force. Dev. Cell 55 (1), 45 (2020).

8. Wegmann, Susanne, Eftekharzadeh, Bahareh, Tepper, Katharina, Zoltowska, Katarzyna M, Bennett, Rachel E, Dujardin, Simon, Laskowski, Pawel R, MacKenzie, Danny, Kamath, Tarun, Commins, Caitlin, Vanderburg, Charles, Roe, Allyson D, Fan, Zhanyun, Molliex, Amandine M, Hernandez-Vega, Amayra, Muller, Daniel, Hyman, Anthony A, Mandelkow, Eckhard, Taylor, J Paul, and Hyman, Bradley T, Tau protein liquid–liquid phase separation can initiate tau aggregation. EMBO J. 37 (7), e98049 (2018).

9. Taylor, J. Paul, Brown, Robert H., and Cleveland, Don W., Decoding ALS: from genes to mechanism. Nature 539 (7628), 197 (2016).

10. Davis, Richoo B., Moosa, Mahdi Muhammad, and Banerjee, Priya R., Ectopic biomolecular phase transitions: fusion proteins in cancer pathologies. Trends Cell Biol. 32 (8), 681 (2022).

11. Wiedner, H. J. and Giudice, J., It’s not just a phase: function and characteristics of RNA-binding proteins in phase separation. Nat Struct Mol Biol 28 (6), 465 (2021).

12. Hyman, Anthony A., Weber, Christoph A., and Jülicher, Frank, Liquid-Liquid Phase Separation in Biology. Annu. Rev. Cell. Dev. Biol. 30 (Volume 30, 2014), 39 (2014).

13. Pappu, Rohit V., Cohen, Samuel R., Dar, Furqan, Farag, Mina, and Kar, Mrityunjoy, Phase Transitions of Associative Biomacromolecules. Chem. Rev. 123 (14), 8945 (2023).

14. Guo, Qi, Shi, Xiangmin, and Wang, Xiangting, RNA and liquid-liquid phase separation. Non-Coding RNA Res. 6 (2), 92 (2021).

15. Mittag, T. and Pappu, R. V., A conceptual framework for understanding phase separation and addressing open questions and challenges. Mol. Cell 82 (12), 2201 (2022).

16. Schmit, J. D., Bouchard, J. J., Martin, E. W., and Mittag, T., Protein Network Structure Enables Switching between Liquid and Gel States. J. Am. Chem. Soc. 142 (2), 874 (2020).

17. Dignon, G. L., Best, R. B., and Mittal, J., Biomolecular Phase Separation: From Molecular Driving Forces to Macroscopic Properties. Annu. Rev. Phys. Chem. 71, 53 (2020).

18. Villegas, José A., Heidenreich, Meta, and Levy, Emmanuel D., Molecular and environmental determinants of biomolecular condensate formation. Nat. Chem. Biol. 18 (12), 1319 (2022).

19. Li, S., Zhang, Y., and Chen, J., Backbone interactions and secondary structures in phase separation of disordered proteins. Biochem. Soc. Trans. 52 (1), 319 (2024).

20. Pappu, R. V., Cohen, S. R., Dar, F., Farag, M., and Kar, M., Phase Transitions of Associative Biomacromolecules. Chem. Rev. 123 (14), 8945 (2023).

21. McCarty, J., Delaney, K. T., Danielsen, S. P. O., Fredrickson, G. H., and Shea, J. E., Complete Phase Diagram for Liquid-Liquid Phase Separation of Intrinsically Disordered Proteins. J Phys Chem Lett 10 (8), 1644 (2019).

22. Li, Siao-Fong and Muthukumar, Murugappan, Theory of Microphase Separation in Concentrated Solutions of Sequence-Specific Charged Heteropolymers. Macromolecules 55 (13), 5535 (2022).

23. Wessén, Jonas, Das, Suman, Pal, Tanmoy, and Chan, Hue Sun, Analytical Formulation and Field-Theoretic Simulation of Sequence-Specific Phase Separation of Protein-Like Heteropolymers with Short- and Long-Spatial-Range Interactions. The Journal of Physical Chemistry B 126 (45), 9222 (2022).

24. Biswas, Subhadip and Potoyan, Davit A., Molecular Drivers of Aging in Biomolecular Condensates: Desolvation, Rigidification, and Sticker Lifetimes. PRX Life 2 (2), 023011 (2024).

25. Valdes-Garcia, G., Heo, L., Lapidus, L. J., and Feig, M., Modeling Concentration-dependent Phase Separation Processes Involving Peptides and RNA via Residue-Based Coarse-Graining. J. Chem. Theory Comput. (2023).

26. Wadsworth, G. M., Srinivasan, S., Lai, L. B., Datta, M., Gopalan, V., and Banerjee, P. R., RNA-driven phase transitions in biomolecular condensates. Mol. Cell 84 (19), 3692 (2024).

27. Ripin, N. and Parker, R., Formation, function, and pathology of RNP granules. Cell 186 (22), 4737 (2023).

28. Castello, Alfredo, Fischer, Bernd, Eichelbaum, Katrin, Horos, Rastislav, Beckmann, Benedikt M., Strein, Claudia, Davey, Norman E., Humphreys, David T., Preiss, Thomas, Steinmetz, Lars M., Krijgsveld, Jeroen, and Hentze, Matthias W., Insights into RNA Biology from an Atlas of Mammalian mRNA-Binding Proteins. Cell 149 (6), 1393 (2012).

29. Molliex, Amandine, Temirov, Jamshid, Lee, Jihun, Coughlin, Maura, Kanagaraj, Anderson P, Kim, Hong Joo, Mittag, Tanja, and Taylor, J. Paul, Phase Separation by Low Complexity Domains Promotes Stress Granule Assembly and Drives Pathological Fibrillization. Cell 163 (1), 123 (2015).

30. Maharana, Shovamayee, Wang, Jie, Papadopoulos, Dimitrios K., Richter, Doris, Pozniakovsky, Andrey, Poser, Ina, Bickle, Marc, Rizk, Sandra, Guillén-Boixet, Jordina, Franzmann, Titus M., Jahnel, Marcus, Marrone, Lara, Chang, Young-Tae, Sterneckert, Jared, Tomancak, Pavel, Hyman, Anthony A., and Alberti, Simon, RNA buffers the phase separation behavior of prion-like RNA binding proteins. Science 360 (6391), 918 (2018).

31. Elbaum-Garfinkle, Shana, Kim, Younghoon, Szczepaniak, Krzysztof, Chen, Carlos Chih-Hsiung, Eckmann, Christian R., Myong, Sua, and Brangwynne, Clifford P., The disordered P granule protein LAF-1 drives phase separation into droplets with tunable viscosity and dynamics. Proc. Natl. Acad. Sci. U. S. A. 112 (23), 7189 (2015).

32. Protter, David S. W., Rao, Bhalchandra S., Van Treeck, Briana, Lin, Yuan, Mizoue, Laura, Rosen, Michael K., and Parker, Roy, Intrinsically Disordered Regions Can Contribute Promiscuous Interactions to RNP Granule Assembly. Cell Reports 22 (6), 1401 (2018).

33. Yoshizawa, Takuya, Ali, Rustam, Jiou, Jenny, Fung, Ho Yee Joyce, Burke, Kathleen A., Kim, Seung Joong, Lin, Yuan, Peeples, William B., Saltzberg, Daniel, Soniat, Michael, Baumhardt, Jordan M., Oldenbourg, Rudolf, Sali, Andrej, Fawzi, Nicolas L., Rosen, Michael K, and Chook, Yuh Min, Nuclear Import Receptor Inhibits Phase Separation of FUS through Binding to Multiple Sites. Cell 173 (3), 693 (2018).

34. Lin, Yuan, Protter, David S. W., Rosen, Michael K., and Parker, Roy, Formation and Maturation of Phase-Separated Liquid Droplets by RNA-Binding Proteins. Mol. Cell 60 (2), 208 (2015).

35. Pessina, Fabio, Giavazzi, Fabio, Yin, Yandong, Gioia, Ubaldo, Vitelli, Valerio, Galbiati, Alessandro, Barozzi, Sara, Garre, Massimiliano, Oldani, Amanda, Flaus, Andrew, Cerbino, Roberto, Parazzoli, Dario, Rothenberg, Eli, and d’Adda di Fagagna, Fabrizio, Functional transcription promoters at DNA double-strand breaks mediate RNA-driven phase separation of damage-response factors. Nat. Cell Biol. 21 (10), 1286 (2019).

36. Langdon, Erin M., Qiu, Yupeng, Ghanbari Niaki, Amirhossein, McLaughlin, Grace A., Weidmann, Chase A., Gerbich, Therese M., Smith, Jean A., Crutchley, John M., Termini, Christina M., Weeks, Kevin M., Myong, Sua, and Gladfelter, Amy S., mRNA structure determines specificity of a polyQ-driven phase separation. Science 360 (6391), 922 (2018).

37. Zhang, Huaiying, Elbaum-Garfinkle, Shana, Langdon, Erin M., Taylor, Nicole, Occhipinti, Patricia, Bridges, Andrew A., Brangwynne, Clifford P., and Gladfelter, Amy S., RNA Controls PolyQ Protein Phase Transitions. Mol. Cell 60 (2), 220 (2015).

38. Van Treeck, Briana and Parker, Roy, Emerging Roles for Intermolecular RNA-RNA Interactions in RNP Assemblies. Cell 174 (4), 791 (2018).

39. Boeynaems, Steven, Holehouse, Alex S., Weinhardt, Venera, Kovacs, Denes, Van Lindt, Joris, Larabell, Carolyn, Van Den Bosch, Ludo, Das, Rhiju, Tompa, Peter S., Pappu, Rohit V., and Gitler, Aaron D., Spontaneous driving forces give rise to protein−RNA condensates with coexisting phases and complex material properties. Proc. Natl. Acad. Sci. U. S. A. 116 (16), 7889 (2019).

40. Jain, Ankur and Vale, Ronald D., RNA phase transitions in repeat expansion disorders. Nature 546 (7657), 243 (2017).

41. Van Treeck, Briana, Protter, David S. W., Matheny, Tyler, Khong, Anthony, Link, Christopher D., and Parker, Roy, RNA self-assembly contributes to stress granule formation and defining the stress granule transcriptome. Proc. Natl. Acad. Sci. U. S. A. 115 (11), 2734 (2018).

42. Poudyal, Raghav R., Sieg, Jacob P., Portz, Bede, Keating, Christine D., and Bevilacqua, Philip C., RNA sequence and structure control assembly and function of RNA condensates. RNA 27 (12), 1589 (2021).

43. Aumiller, William M., Jr., Pir Cakmak, Fatma, Davis, Bradley W., and Keating, Christine D., RNA-Based Coacervates as a Model for Membraneless Organelles: Formation, Properties, and Interfacial Liposome Assembly. Langmuir 32 (39), 10042 (2016).

44. Wadsworth, Gable M., Zahurancik, Walter J., Zeng, Xiangze, Pullara, Paul, Lai, Lien B., Sidharthan, Vaishnavi, Pappu, Rohit V., Gopalan, Venkat, and Banerjee, Priya R., RNAs undergo phase transitions with lower critical solution temperatures. Nature Chemistry 15 (12), 1693 (2023).

45. Tom, Jenna K. A., Onuchic, Paulo L., and Deniz, Ashok A., Short PolyA RNA Homopolymers Undergo Mg2+-Mediated Kinetically Arrested Condensation. The Journal of Physical Chemistry B 126 (46), 9715 (2022).

46. Gatchel, Jennifer R. and Zoghbi, Huda Y., Diseases of Unstable Repeat Expansion: Mechanisms and Common Principles. Nat. Rev. Genet. 6 (10), 743 (2005).

47. Renton, Alan E., Majounie, Elisa, Waite, Adrian, Simón-Sánchez, Javier, Rollinson, Sara, Gibbs, J. Raphael, Schymick, Jennifer C., Laaksovirta, Hannu, van Swieten, John C., Myllykangas, Liisa, Kalimo, Hannu, Paetau, Anders, Abramzon, Yevgeniya, Remes, Anne M., Kaganovich, Alice, Scholz, Sonja W., Duckworth, Jamie, Ding, Jinhui, Harmer, Daniel W., Hernandez, Dena G., Johnson, Janel O., Mok, Kin, Ryten, Mina, Trabzuni, Danyah, Guerreiro, Rita J., Orrell, Richard W., Neal, James, Murray, Alex, Pearson, Justin, Jansen, Iris E., Sondervan, David, Seelaar, Harro, Blake, Derek, Young, Kate, Halliwell, Nicola, Callister, Janis Bennion, Toulson, Greg, Richardson, Anna, Gerhard, Alex, Snowden, Julie, Mann, David, Neary, David, Nalls, Michael A., Peuralinna, Terhi, Jansson, Lilja, Isoviita, Veli-Matti, Kaivorinne, Anna-Lotta, Hölttä-Vuori, Maarit, Ikonen, Elina, Sulkava, Raimo, Benatar, Michael, Wuu, Joanne, Chiò, Adriano, Restagno, Gabriella, Borghero, Giuseppe, Sabatelli, Mario, Heckerman, David, Rogaeva, Ekaterina, Zinman, Lorne, Rothstein, Jeffrey D., Sendtner, Michael, Drepper, Carsten, Eichler, Evan E., Alkan, Can, Abdullaev, Ziedulla, Pack, Svetlana D., Dutra, Amalia, Pak, Evgenia, Hardy, John, Singleton, Andrew, Williams, Nigel M., Heutink, Peter, Pickering-Brown, Stuart, Morris, Huw R., Tienari, Pentti J., and Traynor, Bryan J., A Hexanucleotide Repeat Expansion in C9ORF72 Is the Cause of Chromosome 9p21-Linked ALS-FTD. Neuron 72 (2), 257 (2011).

48. La Spada, Albert R. and Taylor, J. Paul, Repeat expansion disease: progress and puzzles in disease pathogenesis. Nat. Rev. Genet. 11 (4), 247 (2010).

49. Krzyzosiak, Wlodzimierz J., Sobczak, Krzysztof, Wojciechowska, Marzena, Fiszer, Agnieszka, Mykowska, Agnieszka, and Kozlowski, Piotr, Triplet repeat RNA structure and its role as pathogenic agent and therapeutic target. Nucleic Acids Res. 40 (1), 11 (2012).

50. Shea, J. E., Best, R. B., and Mittal, J., Physics-based computational and theoretical approaches to intrinsically disordered proteins. Curr. Opin. Struct. Biol. 67, 219 (2021).

51. Choi, J. M., Holehouse, A. S., and Pappu, R. V., Physical Principles Underlying the Complex Biology of Intracellular Phase Transitions. Annu Rev Biophys 49, 107 (2020).

52. Bari, Khandekar Jishan and Prakashchand, Dube Dheeraj, Fundamental Challenges and Outlook in Simulating Liquid–Liquid Phase Separation of Intrinsically Disordered Proteins. J. Phys. Chem. Lett. 12 (6), 1644 (2021).

53. Shea, Joan-Emma, Best, Robert B., and Mittal, Jeetain, Physics-based computational and theoretical approaches to intrinsically disordered proteins. Curr. Opin. Struct. Biol. 67, 219 (2021).

54. Dignon, Gregory L., Zheng, Wenwei, and Mittal, Jeetain, Simulation methods for liquid–liquid phase separation of disordered proteins. Curr. Opin. Chem. Eng. 23, 92 (2019).

55. Zhang, Yumeng, Li, Shanlong, Gong, Xiping, and Chen, Jianhan, Toward Accurate Simulation of Coupling between Protein Secondary Structure and Phase Separation. J. Am. Chem. Soc. 146 (1), 342 (2024).

56. Choi, J. M., Dar, F., and Pappu, R. V., LASSI: A lattice model for simulating phase transitions of multivalent proteins. PLoS Comput. Biol. 15 (10), e1007028 (2019).

57. Šponer, Jiří, Bussi, Giovanni, Krepl, Miroslav, Banáš, Pavel, Bottaro, Sandro, Cunha, Richard A., Gil-Ley, Alejandro, Pinamonti, Giovanni, Poblete, Simón, Jurečka, Petr, Walter, Nils G., and Otyepka, Michal, RNA Structural Dynamics As Captured by Molecular Simulations: A Comprehensive Overview. Chem. Rev. 118 (8), 4177 (2018).

58. Leontis, N. B. and Westhof, E., Geometric nomenclature and classification of RNA base pairs. RNA 7 (4), 499 (2001).

59. Uusitalo, J. J., Ingolfsson, H. I., Marrink, S. J., and Faustino, I., Martini Coarse-Grained Force Field: Extension to RNA. Biophys. J. 113 (2), 246 (2017).

60. Pasquali, S. and Derreumaux, P., HiRE-RNA: a high resolution coarse-grained energy model for RNA. J Phys Chem B 114 (37), 11957 (2010).

61. Boniecki, M. J., Lach, G., Dawson, W. K., Tomala, K., Lukasz, P., Soltysinski, T., Rother, K. M., and Bujnicki, J. M., SimRNA: a coarse-grained method for RNA folding simulations and 3D structure prediction. Nucleic Acids Res. 44 (7), e63 (2016).

62. Xia, Z., Gardner, D. P., Gutell, R. R., and Ren, P., Coarse-grained model for simulation of RNA three-dimensional structures. J Phys Chem B 114 (42), 13497 (2010).

63. Denesyuk, N. A. and Thirumalai, D., Coarse-grained model for predicting RNA folding thermodynamics. J Phys Chem B 117 (17), 4901 (2013).

64. Bernauer, J., Huang, X., Sim, A. Y., and Levitt, M., Fully differentiable coarse-grained and all-atom knowledge-based potentials for RNA structure evaluation. RNA 17 (6), 1066 (2011).

65. Valdes-Garcia, Gilberto, Heo, Lim, Lapidus, Lisa J., and Feig, Michael, Modeling Concentration-dependent Phase Separation Processes Involving Peptides and RNA via Residue-Based Coarse-Graining. J. Chem. Theory Comput. 19 (2), 669 (2023).

66. Joseph, Jerelle A., Reinhardt, Aleks, Aguirre, Anne, Chew, Pin Yu, Russell, Kieran O., Espinosa, Jorge R., Garaizar, Adiran, and Collepardo-Guevara, Rosana, Physics-driven coarse-grained model for biomolecular phase separation with near-quantitative accuracy. Nature Computational Science 1 (11), 732 (2021).

67. Regy, Roshan Mammen, Dignon, Gregory L., Zheng, Wenwei, Kim, Young C., and Mittal, Jeetain, Sequence dependent phase separation of protein-polynucleotide mixtures elucidated using molecular simulations. Nucleic Acids Res. 48 (22), 12593 (2020).

68. Joseph, Jerelle A., Espinosa, Jorge R., Sanchez-Burgos, Ignacio, Garaizar, Adiran, Frenkel, Daan, and Collepardo-Guevara, Rosana, Thermodynamics and kinetics of phase separation of protein-RNA mixtures by a minimal model. Biophys. J. 120 (7), 1219 (2021).

69. Nguyen, Hung T., Hori, Naoto, and Thirumalai, D., Condensates in RNA repeat sequences are heterogeneously organized and exhibit reptation dynamics. Nature Chemistry 14 (7), 775 (2022).

70. Liu, X. R. and Chen, J. H., HyRes: a coarse-grained model for multi-scale enhanced sampling of disordered protein conformations. Phys. Chem. Chem. Phys. 19 (48), 32421 (2017).

71. Zhang, Y., Liu, X., and Chen, J., Toward Accurate Coarse-Grained Simulations of Disordered Proteins and Their Dynamic Interactions. J. Chem. Inf. Model. 62 (18), 4523 (2022).

72. Bulacu, Monica, Goga, Nicolae, Zhao, Wei, Rossi, Giulia, Monticelli, Luca, Periole, Xavier, Tieleman, D. Peter, and Marrink, Siewert J., Improved Angle Potentials for Coarse-Grained Molecular Dynamics Simulations. J. Chem. Theory Comput. 9 (8), 3282 (2013).

73. Uusitalo, Jaakko J., Ingólfsson, Helgi I., Akhshi, Parisa, Tieleman, D. Peter, and Marrink, Siewert J., Martini Coarse-Grained Force Field: Extension to DNA. J. Chem. Theory Comput. 11 (8), 3932 (2015).

74. Li, Jun and Chen, Shi-Jie, RNAJP: enhanced RNA 3D structure predictions with non-canonical interactions and global topology sampling. Nucleic Acids Res. 51 (7), 3341 (2023).

75. Plumridge, Alex, Andresen, Kurt, and Pollack, Lois, Visualizing Disordered Single-Stranded RNA: Connecting Sequence, Structure, and Electrostatics. J. Am. Chem. Soc. 142 (1), 109 (2020).

76. Draper, David E., A guide to ions and RNA structure. RNA 10 (3), 335 (2004).

77. Bowman, Jessica C., Lenz, Timothy K., Hud, Nicholas V., and Williams, Loren Dean, Cations in charge: magnesium ions in RNA folding and catalysis. Curr. Opin. Struct. Biol. 22 (3), 262 (2012).

78. Draper, David E., Grilley, Dan, and Soto, Ana Maria, Ions and RNA Folding. Annu. Rev. Biophys. Biomol. Struct. 34 (1), 221 (2005).

79. Hayes, R. L., Noel, J. K., Mandic, A., Whitford, P. C., Sanbonmatsu, K. Y., Mohanty, U., and Onuchic, J. N., Generalized Manning Condensation Model Captures the RNA Ion Atmosphere. Phys. Rev. Lett. 114 (25), 258105 (2015).

80. Yu, Tao and Chen, Shi-Jie, Hexahydrated Mg2+ Binding and Outer-Shell Dehydration on RNA Surface. Biophys. J. 114 (6), 1274 (2018).

81. Nguyen, Hung T., Hori, Naoto, and Thirumalai, D., Theory and simulations for RNA folding in mixtures of monovalent and divalent cations. Proc. Natl. Acad. Sci. U. S. A. 116 (42), 21022 (2019).

82. Xi, Kun, Wang, Feng-Hua, Xiong, Gui, Zhang, Zhong-Liang, and Tan, Zhi-Jie, Competitive Binding of Mg2+ and Na+ Ions to Nucleic Acids: From Helices to Tertiary Structures. Biophys. J. 114 (8), 1776 (2018).

83. Misra, Vinod K. and Draper, David E., The interpretation of Mg2+ binding isotherms for nucleic acids using Poisson-Boltzmann theory1 1Edited by B. Honig. J. Mol. Biol. 294 (5), 1135 (1999).

84. Jacobson, David R. and Saleh, Omar A., Counting the ions surrounding nucleic acids. Nucleic Acids Res. 45 (4), 1596 (2017).

85. Bai, Yu, Greenfeld, Max, Travers, Kevin J., Chu, Vincent B., Lipfert, Jan, Doniach, Sebastian, and Herschlag, Daniel, Quantitative and Comprehensive Decomposition of the Ion Atmosphere around Nucleic Acids. J. Am. Chem. Soc. 129 (48), 14981 (2007).

86. Grilley, Dan, Soto, Ana Maria, and Draper, David E., Mg2+–RNA interaction free energies and their relationship to the folding of RNA tertiary structures. Proc. Natl. Acad. Sci. U. S. A. 103 (38), 14003 (2006).

87. Malmberg, C. G. and Maryott, A. A., Dielectric constant of water from 0 to 100 C. J. RES. NATL. BUR. STAN. 56 (1), 1 (1956).

88. Ylitalo, Andrew S., Balzer, Christopher, Zhang, Pengfei, and Wang, Zhen-Gang, Electrostatic Correlations and Temperature-Dependent Dielectric Constant Can Model LCST in Polyelectrolyte Complex Coacervation. Macromolecules 54 (24), 11326 (2021).

89. Adhikari, Sabin, Prabhu, Vivek M., and Muthukumar, Murugappan, Lower Critical Solution Temperature Behavior in Polyelectrolyte Complex Coacervates. Macromolecules 52 (18), 6998 (2019).

90. Singh, Aditya N. and Yethiraj, Arun, Driving Force for the Complexation of Charged Polypeptides. J. Phys. Chem. B 124 (7), 1285 (2020).

91. Eisenberg, Henryk and Felsenfeld, Gary, Studies of the temperature-dependent conformation and phase separation of polyriboadenylic acid solutions at neutral pH. J. Mol. Biol. 30 (1), 17 (1967).

92. Pullara, Paul, Alshareedah, Ibraheem, and Banerjee, Priya R., Temperature-dependent reentrant phase transition of RNA–polycation mixtures. Soft Matter 18 (7), 1342 (2022).

93. Zeng, Xiangze, Liu, Chengwen, Fossat, Martin J., Ren, Pengyu, Chilkoti, Ashutosh, and Pappu, Rohit V., Design of intrinsically disordered proteins that undergo phase transitions with lower critical solution temperatures. APL Materials 9 (2) (2021).

94. Chen, Huimin, Meisburger, Steve P., Pabit, Suzette A., Sutton, Julie L., Webb, Watt W., and Pollack, Lois, Ionic strength-dependent persistence lengths of single-stranded RNA and DNA. Proc. Natl. Acad. Sci. U. S. A. 109 (3), 799 (2012).

95. Richards, Edward G., Flessel, C. Peter, and Fresco, Jacques R., Polynucleotides. VI. Molecular properties and conformation of polyribouridylic acid. Biopolymers 1 (5), 431 (1963).

96. Inners, L. Daniel and Felsenfeld, Gary, Conformation of polyribouridylic acid in solution. J. Mol. Biol. 50 (2), 373 (1970).

97. Kapoor, Utkarsh, Kim, Young C., and Mittal, Jeetain, Coarse-Grained Models to Study Protein–DNA Interactions and Liquid–Liquid Phase Separation. J. Chem. Theory Comput. 20 (4), 1717 (2024).

98. de Mezer, Mateusz, Wojciechowska, Marzena, Napierala, Marek, Sobczak, Krzysztof, and Krzyzosiak, Wlodzimierz J., Mutant CAG repeats of Huntingtin transcript fold into hairpins, form nuclear foci and are targets for RNA interference. Nucleic Acids Res. 39 (9), 3852 (2011).

99. Theimer, Carla A., Finger, L. David, Trantirek, Lukas, and Feigon, Juli, Mutations linked to dyskeratosis congenita cause changes in the structural equilibrium in telomerase RNA. Proc. Natl. Acad. Sci. U. S. A. 100 (2), 449 (2003).

100. Walker, Christopher C., Meek, Garrett A., Fobe, Theodore L., and Shirts, Michael R., Using a Coarse-Grained Modeling Framework to Identify Oligomeric Motifs with Tunable Secondary Structure. J. Chem. Theory Comput. 17 (10), 6018 (2021).

101. Kankia, Besik I., Binding of Mg2+ to single-stranded polynucleotides: hydration and optical studies. Biophys. Chem. 104 (3), 643 (2003).

102. Tan, Zhi-Jie and Chen, Shi-Jie, Electrostatic correlations and fluctuations for ion binding to a finite length polyelectrolyte. J. Chem. Phys. 122 (4), 044903 (2005).

103. Kirmizialtin, Serdal, Silalahi, Alexander R. J., Elber, Ron, and Fenley, Marcia O., The Ionic Atmosphere around A-RNA: Poisson-Boltzmann and Molecular Dynamics Simulations. Biophys. J. 102 (4), 829 (2012).

104. Tan, Zhi-Jie and Chen, Shi-Jie, Predicting Ion Binding Properties for RNA Tertiary Structures. Biophys. J. 99 (5), 1565 (2010).

105. Pörschke, Dietmar, The mode of Mg ++ binding to oligonucleotides. Inner sphere complexes as markers for recognition? Nucleic Acids Res. 6 (3), 883 (1979).

106. Nguyen, Hung T. and Thirumalai, D., Charge Density of Cation Determines Inner versus Outer Shell Coordination to Phosphate in RNA. J. Phys. Chem. B 124 (20), 4114 (2020).

107. Onuchic, Paulo L., Milin, Anthony N., Alshareedah, Ibraheem, Deniz, Ashok A., and Banerjee, Priya R., Divalent cations can control a switch-like behavior in heterotypic and homotypic RNA coacervates. Sci. Rep. 9 (1), 12161 (2019).

108. Ramachandran, V. and Potoyan, D. A., Energy landscapes of homopolymeric RNAs revealed by deep unsupervised learning. Biophys. J. 123 (9), 1152 (2024).

109. Kiliszek, Agnieszka, Kierzek, Ryszard, Krzyzosiak, Wlodzimierz J., and Rypniewski, Wojciech, Atomic resolution structure of CAG RNA repeats: structural insights and implications for the trinucleotide repeat expansion diseases. Nucleic Acids Res. 38 (22), 8370 (2010).

110. Sobczak, Krzysztof, Michlewski, Gracjan, de Mezer, Mateusz, Kierzek, Elzbieta, Krol, Jacek, Olejniczak, Marta, Kierzek, Ryszard, and Krzyzosiak, Wlodzimierz J., Structural Diversity of Triplet Repeat RNAs. J. Biol. Chem. 285 (17), 12755 (2010).

111. Lemieux, Sébastien and Major, François, RNA canonical and non-canonical base pairing types: a recognition method and complete repertoire. Nucleic Acids Res. 30 (19), 4250 (2002).

112. Mokdad, Ali, Krasovska, Maryna V., Sponer, Jiri, and Leontis, Neocles B., Structural and evolutionary classification of G/U wobble basepairs in the ribosome. Nucleic Acids Res. 34 (5), 1326 (2006).

113. Abu Almakarem, Amal S., Petrov, Anton I., Stombaugh, Jesse, Zirbel, Craig L., and Leontis, Neocles B., Comprehensive survey and geometric classification of base triples in RNA structures. Nucleic Acids Res. 40 (4), 1407 (2012).

114. Abraham, Mark James, Murtola, Teemu, Schulz, Roland, Páll, Szilárd, Smith, Jeremy C., Hess, Berk, and Lindahl, Erik, GROMACS: High performance molecular simulations through multi-level parallelism from laptops to supercomputers. SoftwareX 1–2, 19 (2015).

115. Bauer, Paul, Hess, Berk, and Lindahl, Erik, GROMACS 2022 Manual. Zenodo (2022).

116. Paul Bauer, Berk Hess, and Lindahl, Erik, GROMACS 2022 Source code (Version 2022). Zenodo (2022).

117. Case, David A., Aktulga, Hasan Metin, Belfon, Kellon, Cerutti, David S., Cisneros, G. Andrés, Cruzeiro, Vinícius Wilian D., Forouzesh, Negin, Giese, Timothy J., Götz, Andreas W., Gohlke, Holger, Izadi, Saeed, Kasavajhala, Koushik, Kaymak, Mehmet C., King, Edward, Kurtzman, Tom, Lee, Tai-Sung, Li, Pengfei, Liu, Jian, Luchko, Tyler, Luo, Ray, Manathunga, Madushanka, Machado, Matias R., Nguyen, Hai Minh, O’Hearn, Kurt A., Onufriev, Alexey V., Pan, Feng, Pantano, Sergio, Qi, Ruxi, Rahnamoun, Ali, Risheh, Ali, Schott-Verdugo, Stephan, Shajan, Akhil, Swails, Jason, Wang, Junmei, Wei, Haixin, Wu, Xiongwu, Wu, Yongxian, Zhang, Shi, Zhao, Shiji, Zhu, Qiang, Cheatham, Thomas E., III, Roe, Daniel R., Roitberg, Adrian, Simmerling, Carlos, York, Darrin M., Nagan, Maria C., and Merz, Kenneth M., Jr., AmberTools. J. Chem. Inf. Model. 63 (20), 6183 (2023).

118. Cornell, Wendy D., Cieplak, Piotr, Bayly, Christopher I., Gould, Ian R., Merz, Kenneth M., Ferguson, David M., Spellmeyer, David C., Fox, Thomas, Caldwell, James W., and Kollman, Peter A., A Second Generation Force Field for the Simulation of Proteins, Nucleic Acids, and Organic Molecules. J. Am. Chem. Soc. 117 (19), 5179 (1995).

119. Jorgensen, William L., Chandrasekhar, Jayaraman, Madura, Jeffry D., Impey, Roger W., and Klein, Michael L., Comparison of simple potential functions for simulating liquid water. J. Chem. Phys. 79 (2), 926 (1983).

120. Zgarbová, Marie, Otyepka, Michal, Šponer, Jiří, Mládek, Arnošt, Banáš, Pavel, Cheatham, Thomas E., III, and Jurečka, Petr, Refinement of the Cornell et al. Nucleic Acids Force Field Based on Reference Quantum Chemical Calculations of Glycosidic Torsion Profiles. J. Chem. Theory Comput. 7 (9), 2886 (2011).

121. Pérez, Alberto, Marchán, Iván, Svozil, Daniel, Sponer, Jiri, Cheatham, Thomas E., Laughton, Charles A., and Orozco, Modesto, Refinement of the AMBER Force Field for Nucleic Acids: Improving the Description of α/γ Conformers. Biophys. J. 92 (11), 3817 (2007).

122. Grotz, Kara K., Cruz-León, Sergio, and Schwierz, Nadine, Optimized Magnesium Force Field Parameters for Biomolecular Simulations with Accurate Solvation, Ion-Binding, and Water-Exchange Properties. J. Chem. Theory Comput. 17 (4), 2530 (2021).

123. Mamatkulov, Shavkat and Schwierz, Nadine, Force fields for monovalent and divalent metal cations in TIP3P water based on thermodynamic and kinetic properties. J. Chem. Phys. 148 (7) (2018).

124. Martínez, L., Andrade, R., Birgin, E. G., and Martínez, J. M., PACKMOL: A package for building initial configurations for molecular dynamics simulations. J. Comput. Chem. 30 (13), 2157 (2009).

125. Jo, Sunhwan, Kim, Taehoon, Iyer, Vidyashankara G., and Im, Wonpil, CHARMM-GUI: A web-based graphical user interface for CHARMM. J. Comput. Chem. 29 (11), 1859 (2008).

126. Lee, Jumin, Cheng, Xi, Swails, Jason M., Yeom, Min Sun, Eastman, Peter K., Lemkul, Justin A., Wei, Shuai, Buckner, Joshua, Jeong, Jong Cheol, Qi, Yifei, Jo, Sunhwan, Pande, Vijay S., Case, David A., Brooks, Charles L., III, MacKerell, Alexander D., Jr., Klauda, Jeffery B., and Im, Wonpil, CHARMM-GUI Input Generator for NAMD, GROMACS, AMBER, OpenMM, and CHARMM/OpenMM Simulations Using the CHARMM36 Additive Force Field. J. Chem. Theory Comput. 12 (1), 405 (2016).

127. Lee, Jumin, Hitzenberger, Manuel, Rieger, Manuel, Kern, Nathan R., Zacharias, Martin, and Im, Wonpil, CHARMM-GUI supports the Amber force fields. J. Chem. Phys. 153 (3) (2020).

128. Darden, Tom, York, Darrin, and Pedersen, Lee, Particle mesh Ewald: An N⋅log(N) method for Ewald sums in large systems. J. Chem. Phys. 98 (12), 10089 (1993).

129. Essmann, Ulrich, Perera, Lalith, Berkowitz, Max L., Darden, Tom, Lee, Hsing, and Pedersen, Lee G., A smooth particle mesh Ewald method. J. Chem. Phys. 103 (19), 8577 (1995).

130. Bussi, Giovanni, Donadio, Davide, and Parrinello, Michele, Canonical sampling through velocity rescaling. J. Chem. Phys. 126 (1) (2007).

131. Nosé, Shuichi and Klein, M. L., Constant pressure molecular dynamics for molecular systems. Mol. Phys. 50 (5), 1055 (1983).

132. Yoo, Jejoong and Aksimentiev, Aleksei, Competitive Binding of Cations to Duplex DNA Revealed through Molecular Dynamics Simulations. J. Phys. Chem. B 116 (43), 12946 (2012).

133. Kirmizialtin, Serdal and Elber, Ron, Computational Exploration of Mobile Ion Distributions around RNA Duplex. J. Phys. Chem. B 114 (24), 8207 (2010).

134. Eastman, Peter, Galvelis, Raimondas, Peláez, Raúl P., Abreu, Charlles R. A., Farr, Stephen E., Gallicchio, Emilio, Gorenko, Anton, Henry, Michael M., Hu, Frank, Huang, Jing, Krämer, Andreas, Michel, Julien, Mitchell, Joshua A., Pande, Vijay S., Rodrigues, João Pglm, Rodriguez-Guerra, Jaime, Simmonett, Andrew C., Singh, Sukrit, Swails, Jason, Turner, Philip, Wang, Yuanqing, Zhang, Ivy, Chodera, John D., De Fabritiis, Gianni, and Markland, Thomas E., OpenMM 8: Molecular Dynamics Simulation with Machine Learning Potentials. The Journal of Physical Chemistry B 128 (1), 109 (2024).

135. Shirts, Michael R. and Chodera, John D., Statistically optimal analysis of samples from multiple equilibrium states. J. Chem. Phys. 129 (12) (2008).

## Supplementary References

1. Plumridge, A., Andresen, K. and Pollack, L. (2020) Visualizing Disordered Single-Stranded RNA: Connecting Sequence, Structure, and Electrostatics. J. Am. Chem. Soc., 142, 109–119.

2. Theimer, C.A., Finger, L.D., Trantirek, L. and Feigon, J. (2003) Mutations linked to dyskeratosis congenita cause changes in the structural equilibrium in telomerase RNA. Proc. Natl. Acad. Sci. U. S. A., 100, 449–454.

