## Supplemental tables and figures for "Driving Forces of RNA Condensation Revealed through Coarse-Grained Modeling with Explicit Mg^2+^"

Shanlong Li and Jianhan Chen*

Department of Chemistry

University of Massachusetts, Amherst, MA 01003, USA

### Supplementary Tables

**Table S1.** Atomistic to CG mapping scheme (also see Figure 1)

| | Bead name | Bead type | Mapped atoms[a] |
|---|---|---|---|
| **Phosphate** | B1 | BP | P, OP1, OP2, O3’, O5’ |
| **Ribose** | B2 | BC1 | C3’, C4’, C5’, O4’* |
| | B3 | BC2 | C1’, C2’, O2’, O4’* |
| **Adenine (A)** | A1 | N1C | C4, N9 |
| | A2 | CNC | C5, N7, C8 |
| | A3 | NCN | N1, C6, N6 |
| | A4 | N2C | C2, N3 |
| **Guanine (G)** | G1 | N1C | C4, N9 |
| | G2 | CNC | C5, N7, C8 |
| | G3 | OCN | N1, C6, O6 |
| | G4 | NCN | C2, N2, N3 |
| **Cytosine (C)** | C1 | NCC | N1, C5, C6 |
| | C2 | NCN | N3, C4, N4 |
| | C3 | CO | C2, O2 |
| **Uracil (U)** | U1 | NCC | N1, C5, C6 |
| | U2 | OCN | N3, C4, O4 |
| | U3 | CO | C2, O2 |

*: shared atom
[a]: all connected hydrogen atoms are included

**Table S2.** Relative stacking strength to A//A stacking. In the final model, $\varepsilon_{A//A}$ = 2.05 kcal/mol (see main text).

| **stacks** | A//A | A//G | A//C | A//U | G//A | G//G | G//C | G//U |
|---|---|---|---|---|---|---|---|---|
| $\boldsymbol{\varepsilon}_{\mathbf{relative}}$ | 1.0 | 1.0 | 0.8 | 0.8 | 1.0 | 1.0 | 0.8 | 0.8 |
| **stacks** | C//A | C//G | C//C | C//U | U//A | U//G | U//C | U//U |
| $\boldsymbol{\varepsilon}_{\mathbf{relative}}$ | 0.4 | 0.4 | 0.2 | 0.2 | 0.4 | 0.4 | 0.2 | 0.2 |

**Table S3.** Equilibrium distance for different hydrogen bonding CG beads and the optimized hydrogen bonding pair strength (see main text).

| **base pair** | **bead pair** | $\boldsymbol{r_0}$ **(nm)** | $\boldsymbol{\varepsilon_{ij}^{stack}}$ **(kcal/mol)** |
|---|---|---|---|
| C-G | C2-G3 | 0.33 | 2.15 |
| | C3-G4 | 0.38 | 2.15 |
| A-U | A3-U2 | 0.33 | 1.44 |
| | A4-U3 | 0.40 | 1.44 |

**Table S4.** RNA concentrations (*c*), RNA chain numbers (*N*), and box sizes for the simulations of phase separation.

| **RNA** | ***c*** | ***N*** | **box size (nm)** |
|---|---|---|---|
| $rA_{30}/rU_{30}$ | 50 μM | 240 | 200 |
| CAG/CUG/CUU repeats | 10 μM | 100 | 250 |
| | 25 μM | 100 | 190 |
| | 50 μM | 100 | 150 |
| | 100 μM | 100 | 120 |

**Table S5.** The sequences of all RNAs simulated in this work (except RNA tetramers used for deriving bonded parameters, which are provided in the main text).

| RNA | Sequence |
|---|---|
| $rA_{30}$ | AAAAA AAAAA AAAAA AAAAA AAAAA AAAAA |
| $rU_{30}$ | UUUUU UUUUU UUUUU UUUUU UUUUU UUUUU |
| $rU_{40}$ | UUUUU UUUUU UUUUU UUUUU UUUUU UUUUU UUUUU UUUUU |
| $(CAG)_{10}$ | CAGCAGCAGCAGCAG CAGCAGCAGCAGCAG |
| $(CAG)_{20}$ | CAGCAGCAGCAGCAG CAGCAGCAGCAGCAG CAGCAGCAGCAGCAG CAGCAGCAGCAGCAG |
| $(CAG)_{31}$ | CAGCAGCAGCAGCAG CAGCAGCAGCAGCAG CAGCAGCAGCAGCAG CAGCAGCAGCAGCAG CAGCAGCAGCAGCAG CAGCAGCAGCAGCAG CAG |
| $(CUG)_{31}$ | CUGCUGCUGCUGCUG CUGCUGCUGCUGCUG CUGCUGCUGCUGCUG CUGCUGCUGCUGCUG CUGCUGCUGCUGCUG CUGCUGCUGCUGCUG CUG |
| $(CUU)_{31}$ | CUUCUUCUUCUUCUU CUUCUUCUUCUUCUU CUUCUUCUUCUUCUU CUUCUUCUUCUUCUU CUUCUUCUUCUUCUU CUUCUUCUUCUUCUU CUU |
| hTR-hairpin | GGGCU GUUUU UCUCG CUGAC UUUCA GCCCC |
| BWYV-PK | GGGCU GUUUU UCUCG CUGAC UUUCA GCCCCA AACAA AAAAU GUCAG CA |
| 2KOC | GGCAC UUCGG UGCC |
| 1SDR | 5'-UAAGGAGGUGAU-3'<br>3'-AUUCCUCCACUA-5' |
| 1Q9A | UGCUC CUAGU ACGAG AGGAC CGGAG UG |
| 4FNJ | CUGCU GGCUA AGGCA UGAAA GUGCU AUGCC UGCUG |
| 1MNX | GGGUG ACGAU ACUGU AGGCG AGAGC CUGCG GAAAA AUAGC CC |
| 2NC1 | GGGGC UGAAG GAUGC CCAGA GAGAU CUGGG GCCUC GGGAG AUCGA GGUUA AAAAA CGUCU AGGCC CC |

**Table S6.** Bond parameters, $k_b$ in kcal/(mol·Å$^2$) and $b_0$ in Å

| $i$ | $j$ | $k_b$ | $b_0$ | $i$ | $j$ | $k_b$ | $b_0$ |
|---|---|---|---|---|---|---|---|
| B1 | B2 | 15.0 | 3.65 | G1 | G4 | 130.0 | 3.05 |
| B2 | B3 | 100.0 | 2.32 | G2 | G3 | 130.0 | 3.05 |
| B3 | A1/G1 | 150.0 | 1.93 | G3 | G4 | 150.0 | 2.90 |
| B3 | C1/U1 | 150.0 | 2.53 | C1 | C2 | 200.0 | 2.48 |
| A1/G1 | A2/G2 | 500.0 | 1.63 | C1 | C3 | 180.0 | 2.70 |
| A1 | A4 | 200.0 | 2.4 | C2 | C3 | 120.0 | 2.95 |
| A2 | A3 | 120.0 | 3.06 | U1 | U2 | 250.0 | 2.49 |
| A3 | A4 | 200.0 | 2.72 | U2 | U3 | 150.0 | 2.99 |

**Table S7.** Angle parameters, $k_\theta$ in kcal/(mol·rad$^2$) and $\theta_0$ in degree

| $i$ | $j$ | $k$ | $k_\theta$ | $\theta_0$ | $i$ | $j$ | $k$ | $k_\theta$ | $\theta_0$ |
|---|---|---|---|---|---|---|---|---|---|
| B2 | B1 | B2 | 2.0 | 105.0 | B3 | A1 | G4 | 20.0 | 116.0 |
| B1 | B2 | B1 | 4.0 | 105.0 | B3 | C1 | C3 | 50.0 | 59.0 |
| B1 | B2 | B3 | 4.0 | 112.0 | A1 | A2 | A3 | 80.0 | 90.0 |
| B2 | B3 | B1 | 10.0 | 55.0 | A2 | A3 | A4 | 50.0 | 73.0 |
| B2 | B3 | A1/G1 | 4.0 | 136.0 | A3 | A4 | A1 | 50.0 | 86.0 |
| B2 | B3 | C1/U1 | 5.0 | 115.0 | G1 | G2 | G3 | 80.0 | 90.0 |
| B3 | A1/G1 | A2/G2 | 25.0 | 127.0 | G2 | G3 | G4 | 50.0 | 84.0 |
| B3 | A1 | A4 | 25.0 | 117.0 | G3 | G4 | G1 | 50.0 | 72.0 |

**Table S8.** Dihedral angle parameters, $k_X$ in kcal/mol and $\delta$ in degree.

| $i$ | $j$ | $k$ | $l$ | $k_X$ | $n$ | $\delta$ |
|---|---|---|---|---|---|---|
| B1 | B2 | B3 | A1/G1 | 8.0 * | 1 | -165.0 |
| B1 | B2 | B3 | C1/U1 | 5.0 * | 1 | 180.0 |
| B2 | B3 | A1/G1 | A2/G2 | 1.0 | 1 | 165.0 |
| B2 | B3 | C1 | C3 | 5.0 | 1 | 0.0 |
| A1 | A2 | A4 | A3 | 80.0 | 1 | 0.0 |
| G1 | G2 | G4 | G3 | 50.0 | 1 | 0.0 |

* Force constants for these two dihedrals are increased to 35 kcal/mol in the final model to improve local structures reflected in OCF profiles.

**Table S9.** Improper dihedral parameters, $k_\psi$ in kcal/(mol·rad$^2$) and $\psi_0$ in degree

| $i$ | $j$ | $k$ | $l$ | $k_\psi$ | $\psi_0$ |
|---|---|---|---|---|---|
| B2 | B1 | B1* | B3 | 15.0 | 29.0 |
| A1 | B3 | A2 | A4 | 20.0 | 0.0 |
| G1 | B3 | G2 | G4 | 15.0 | 0.0 |
| C1 | B3 | C2 | C3 | 5.0 | -180.0 |
| U1 | B3 | U2 | U3 | 5.0 | -180.0 |

* The bead in the next residue

**Table S10.** Final vdW parameters of all CG bead types.

| | Bead name | Bead type | $\varepsilon_i$ (kcal/mol) | $r_{min}/2$ (Å) |
|---|---|---|---|---|
| **Phosphate** | B1 | BP | -0.11583 | 2.4340 |
| **Ribose** | B2 | BC1 | -0.09922 | 2.2826 |
| | B3 | BC2 | -0.13244 | 2.2826 |
| **Adenine (A)** | A1 | N1C | -0.11583 | 1.7954 |
| | A2 | CNC | -0.13244 | 2.0309 |
| | A3 | NCN | -0.14905 | 2.0089 |
| | A4 | N2C | -0.11583 | 1.8215 |
| **Guanine (G)** | G1 | N1C | -0.11583 | 1.7954 |
| | G2 | CNC | -0.13244 | 2.0309 |
| | G3 | OCN | -0.14905 | 1.9241 |
| | G4 | NCN | -0.14905 | 2.0089 |
| **Cytosine (C)** | C1 | NCC | -0.11583 | 2.0452 |
| | C2 | NCN | -0.14905 | 2.0089 |
| | C3 | CO | -0.09922 | 1.7170 |
| **Uracil (U)** | U1 | NCC | -0.11583 | 2.0452 |
| | U2 | OCN | -0.14905 | 1.9241 |
| | U3 | CO | -0.09922 | 1.7170 |
| **$Mg^{2+}$** | MG | MG | -0.10000 | 3.0000 |

**Table S11.** NBFIX for Lennard_Jones parameters

| Bead type *i* | Bead type *j* | $\varepsilon_{ij}$ (kcal/mol) | $r_{min}/2$ (Å) |
|---|---|---|---|
| BP | MG | -0.50000 | 5.0 |
| NCN | NCN | -0.14905 | 3.6 |
| NCN | OCN | -0.14905 | 3.3 |
| NCN | CO | -0.12414 | 3.8 |
| N2C | CO | -0.10753 | 4.0 |

**Table S12.** $\lambda_{P\text{-}Mg}$ and $\Delta n_{Mg}$ of $rA_{30}$ and $rU_{30}$ at different concentration of $Mg^{2+}$

| **[$Mg^{2+}$] (mM)** | **$rA_{30}$** | | **$rU_{30}$** | |
|---|---|---|---|---|
| | $\lambda_{P\text{-}Mg}$ | $\Delta n_{Mg}$ | $\lambda_{P\text{-}Mg}$ | $\Delta n_{Mg}$ |
| 1 | 0.71 | 0.2749 | 0.67 | 0.2126 |
| 2 | 0.75 | 0.3292 | 0.72 | 0.2827 |
| 5 | 0.78 | 0.3933 | 0.77 | 0.3636 |
| 10 | - | - | 0.80 | 0.4076 |

**Table S13.** Hill fit parameters for $rA_{30}$, $rU_{30}$ and $(CAG)_{31}$

| **RNA** | **$F_{Mg}$** | **$M_{1/2}$** | **n** |
|---|---|---|---|
| $rA_{30}$[a] | 0.54 | 0.94 | 0.59 |
| $rU_{30}$[a] | 0.48 | 1.31 | 0.85 |
| $(CAG)_{31}$ | 0.53 | 0.68 | 0.28 |

a: Experimental data from Ref (1).

### Supplementary Figures

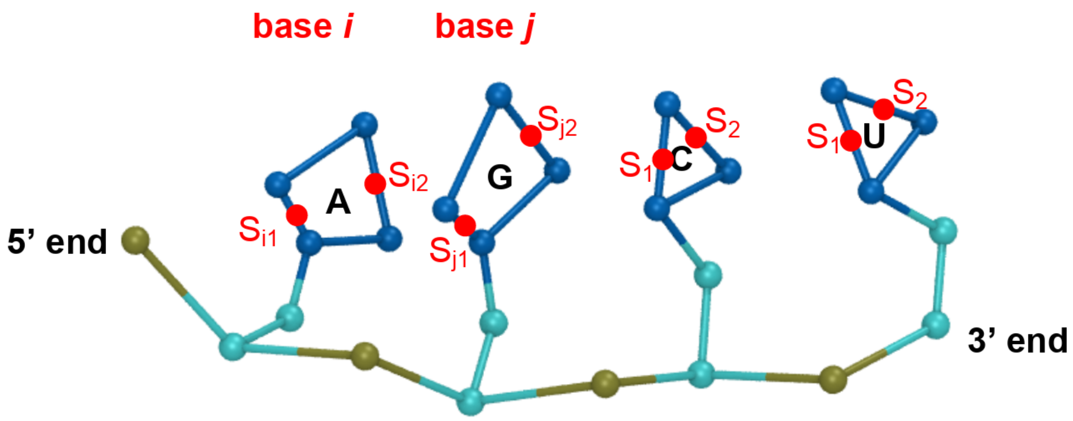


**Figure S1.** Scheme of the assignment of base stacking. Red points $S_{i1}$ and $S_{i2}$ are the two virtual sites of base *i*. From the 5' end to the 3' end, the stacking interaction is applied between $S_{i2}$ and $S_{j1}$.

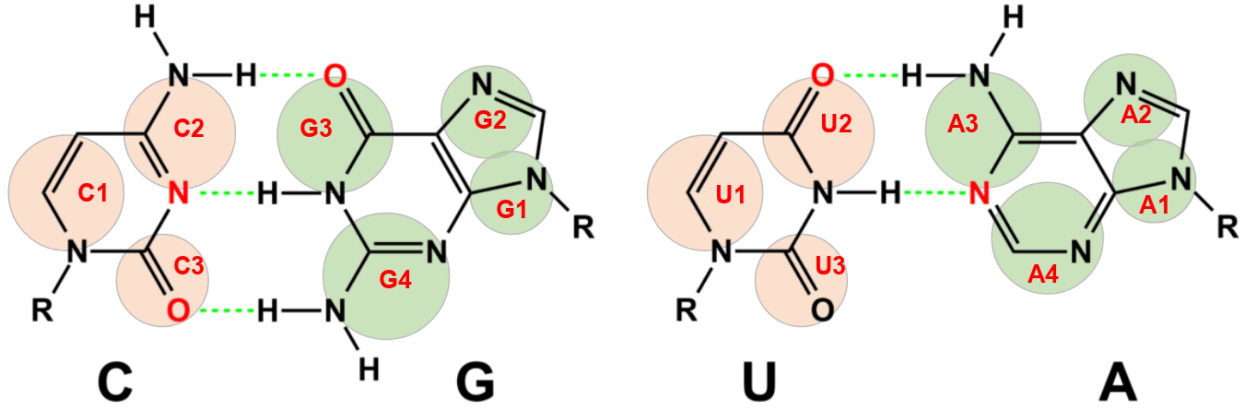


**Figure S2.** Scheme of A-U and G-C base pairing. Related bead names are in red fonts. The green dashed lines show the hydrogen bonds for each pair.

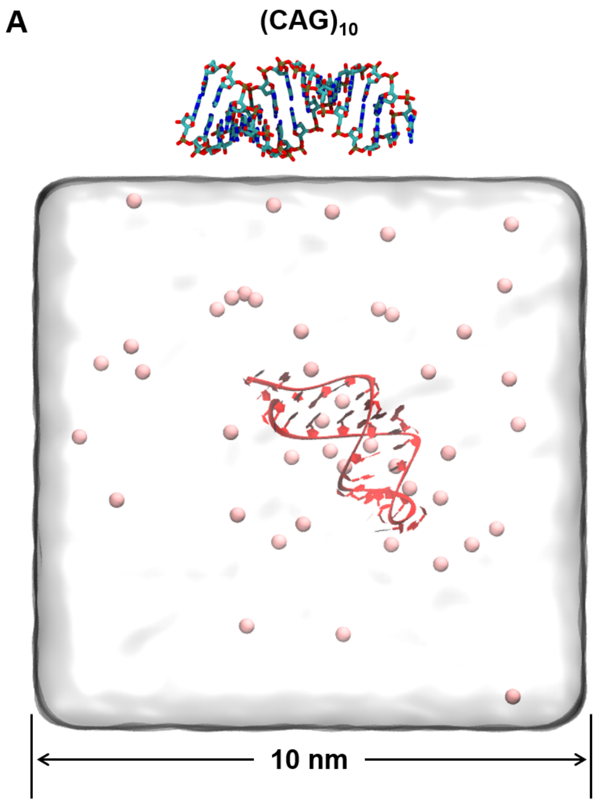


**Figure S3.** The ideal hairpin structures and equilibrated states with $Mg^{2+}$ ions (pink beads) of $(CAG)_{10}$ (**A**) and $(CAG)_{31}$ (**B**).

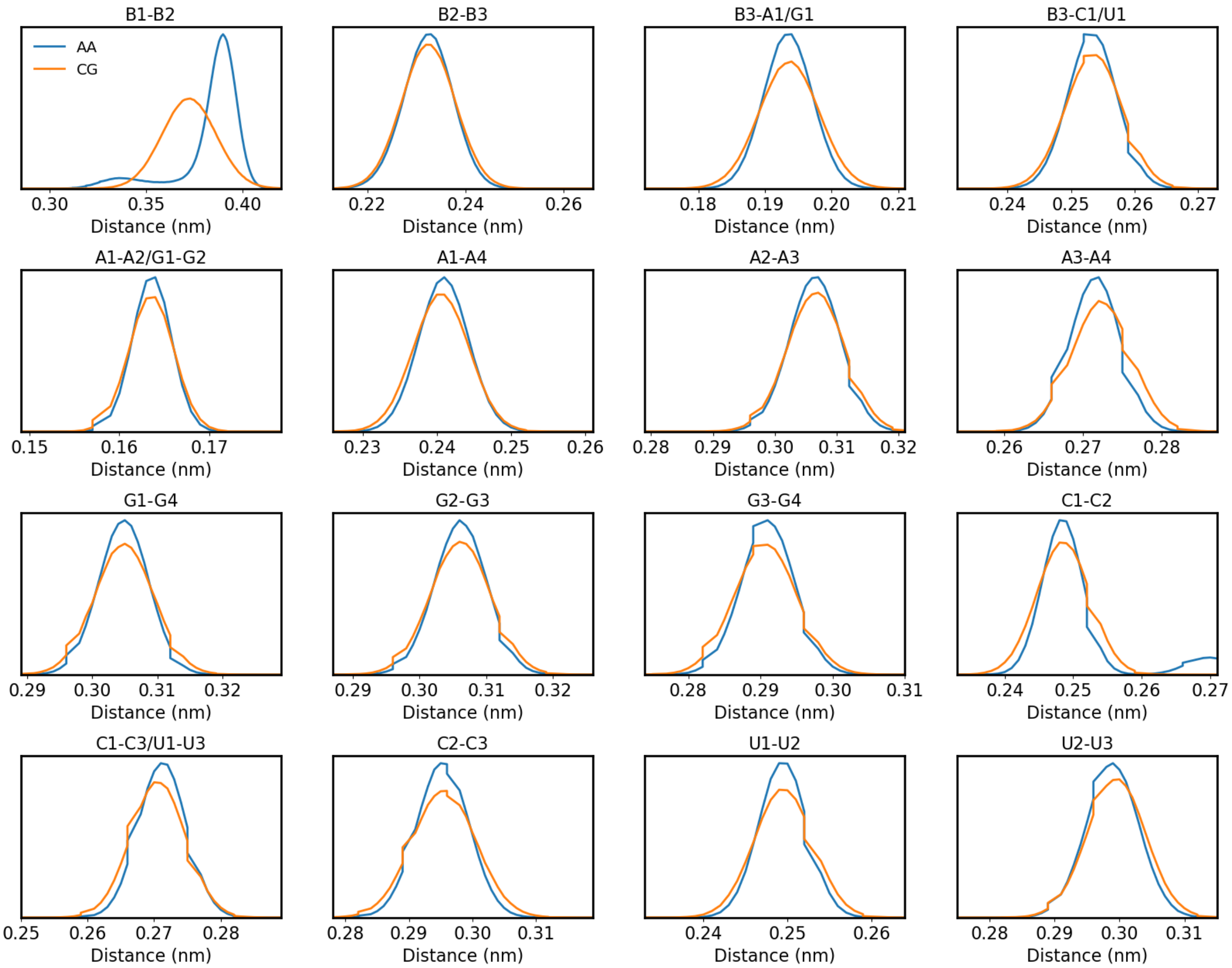


**Figure S4.** AA and CG distributions of bonds. The average distributions from the AMBER AA simulations are shown in blue, whereas the average CG distributions are shown in orange. The title of each subplot lists the bead names that participate in the bond.

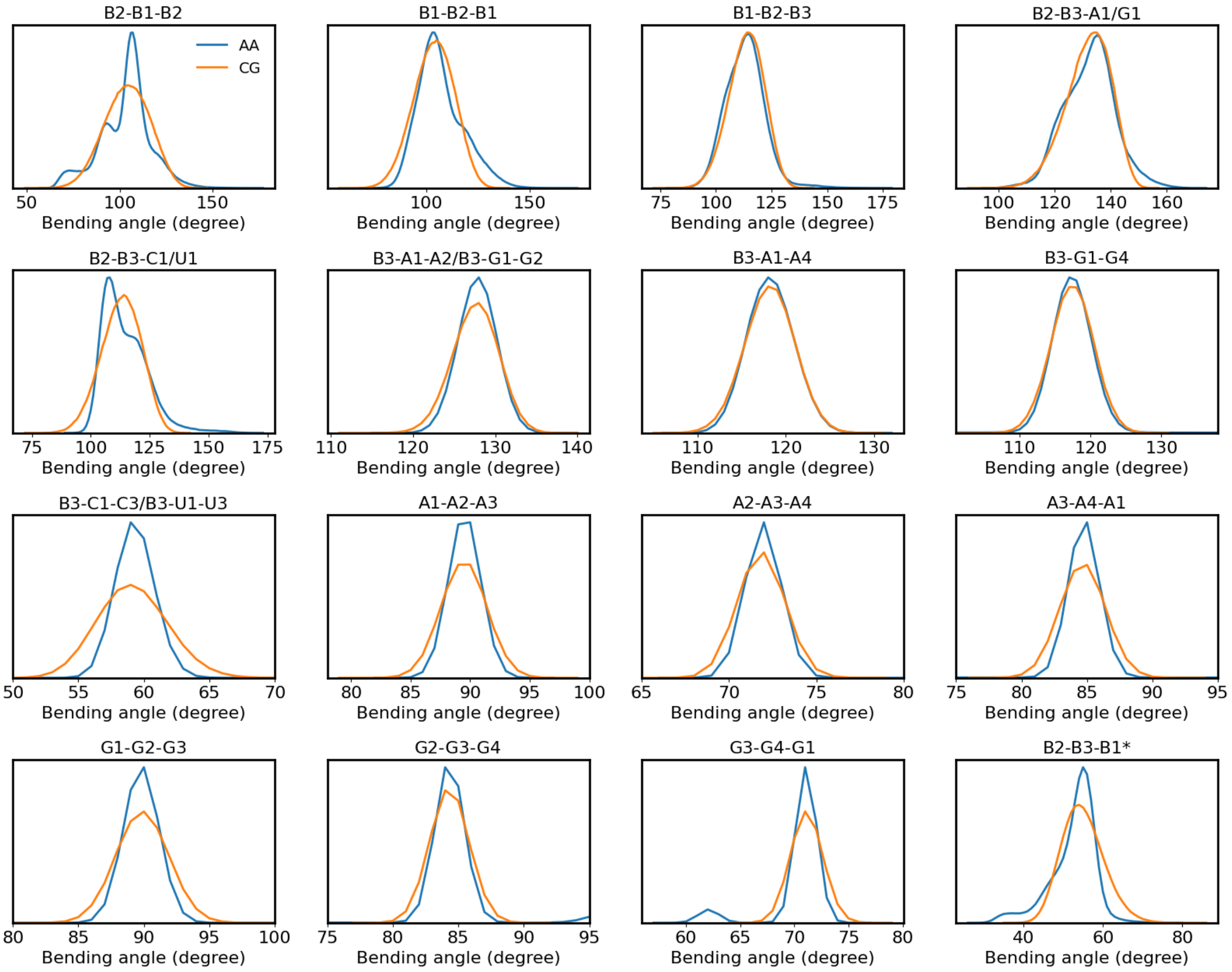


**Figure S5.** AA and CG distributions of angles. The average distribution from the AMBER AA simulations is shown in blue, whereas the average CG distributions are shown in orange. The title of each subplot lists the bead names that participate in the angle following the same order as the angle.

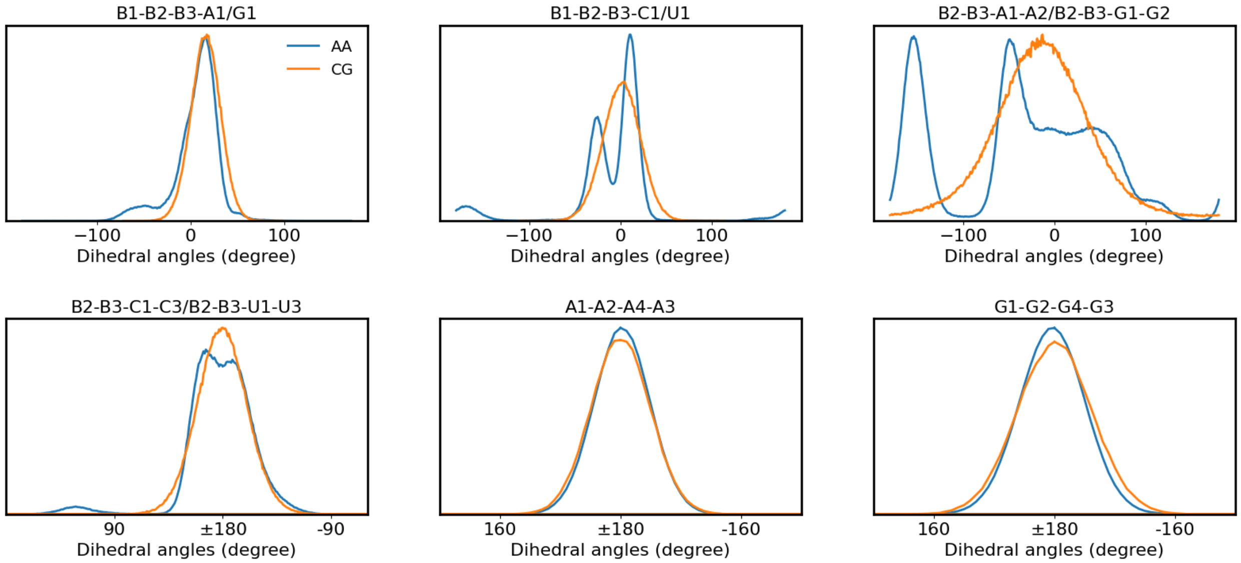


**Figure S6.** AA and CG distributions of dihedral angles. The average distributions from the Amber AA simulation are shown in blue, whereas the average CG distributions are shown in orange. The title of each subplot lists the bead names that participate in the angle following the same order as the dihedral angles.

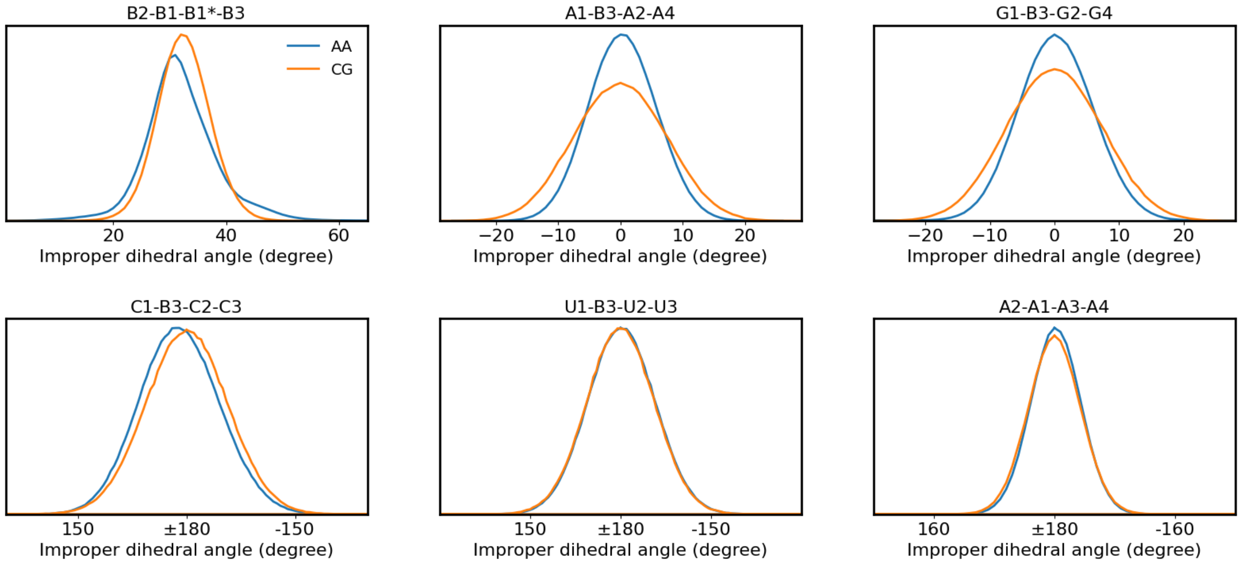


**Figure S7.** AA and CG distributions of dihedral angles and improper dihedral angles. The average distributions from the AMBER AA simulations are shown in blue, whereas the average CG distributions are shown in orange. The title of each subplot lists the bead names that participate in the angle following the same order as the dihedral angles. B1* denotes the B1 bead of the next residue.

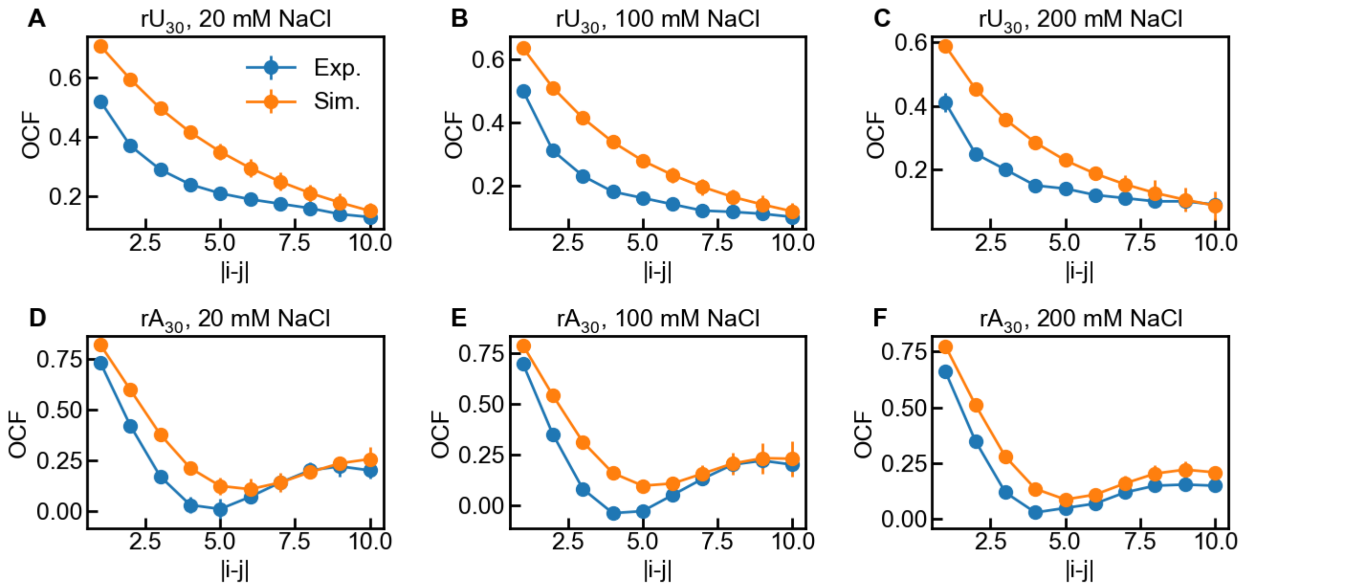


**Figure S8.** Orientation correlation function (OCF) of $rU_{30}$ (**A-C**) and $rA_{30}$ (**D-F**). The NaCl concentrations were set as 20 mM, 100 mM, and 200 mM to show the salt dependence of OCF. |*i*-*j*| is the difference in residue numbers between the *i*th and *j*th phosphate-phosphate bond vectors (Eqn. 6 in main text). Experimental results are shown in blue whereas simulation results are in orange. The error bars of simulation results were estimated from block analysis.

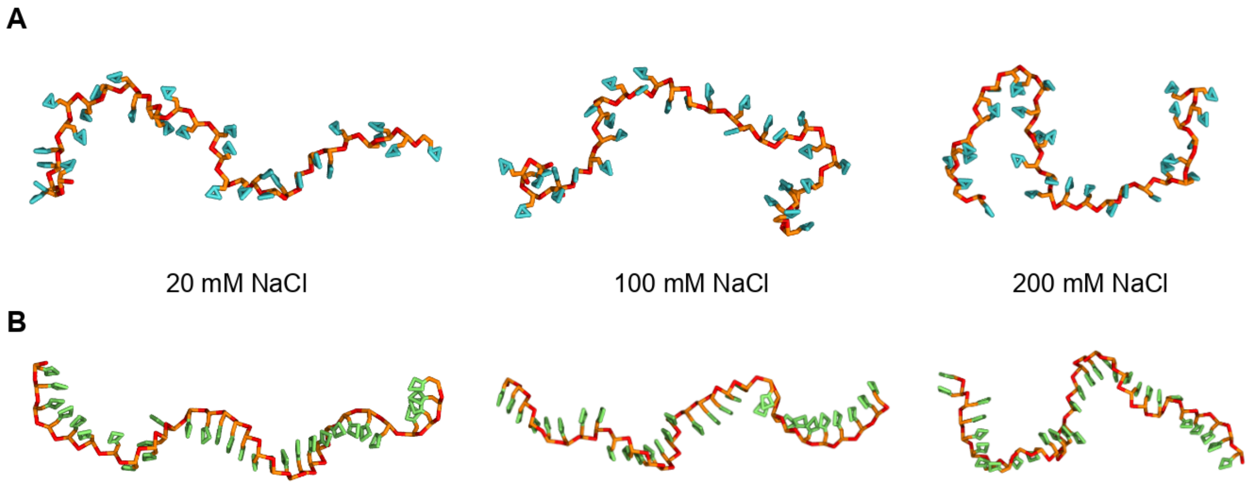


**Figure S9.** Representative structures of (**A**) $rU_{30}$ and (**B**) $rA_{30}$ at 20 mM, 100 mM, and 200 mM NaCl solution. Phosphate beads B1 are shown in red, while B2 and B3 for ribose are in orange. Bases U and A are in cyan and lime, respectively.

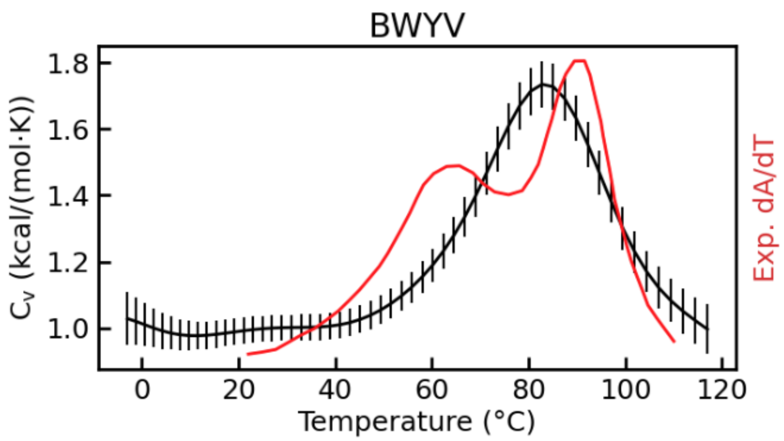


**Figure S10.** Heat capacity ($C_v$) of Beet Western Yellow Virus (BWYV) pseudoknot. Experimental data is the red line, where the first derivative of UV-absorbance with respect to temperature (dA/dT) at 280 nm is used as the reference melting profile (2).

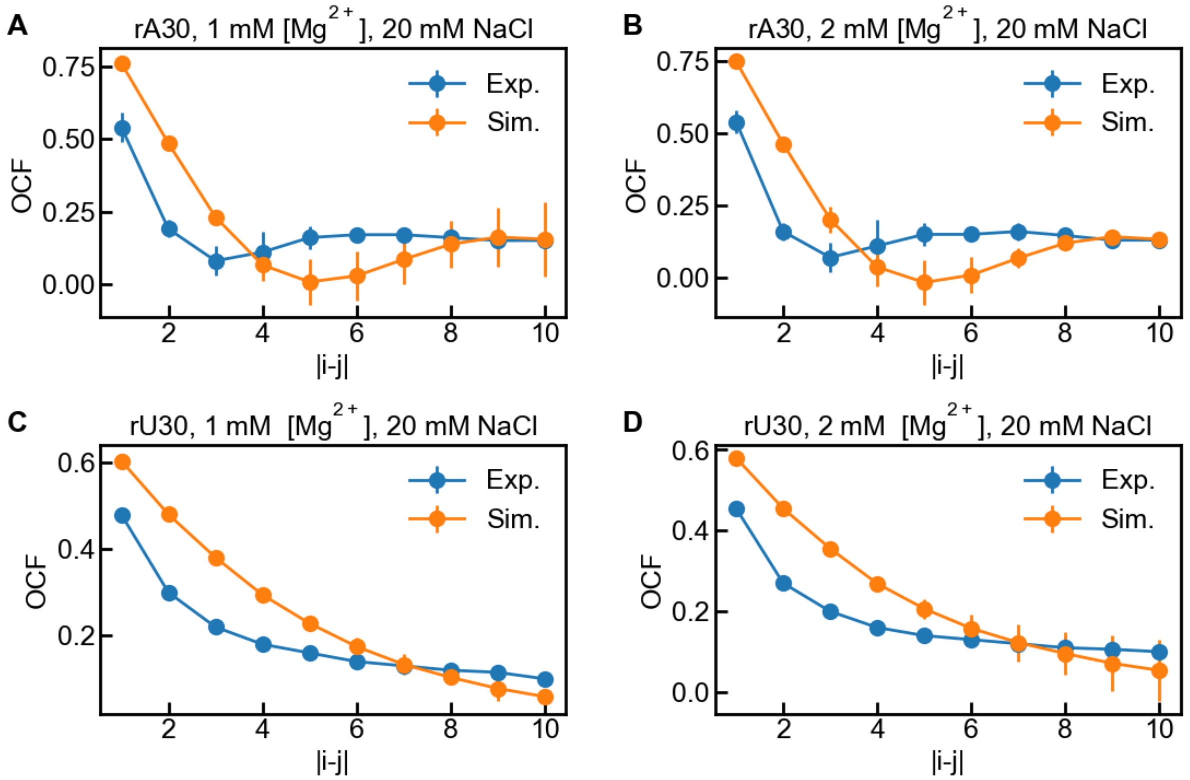


**Figure S11.** OCFs of $rA_{30}$ (**A, B**) and $rU_{30}$ (**C, D**) at different ion conditions. |*i*-*j*| is the difference in residue numbers between the *i*th and *j*th phosphate-phosphate bond vectors (Eqn. 6 in main text). The experimental results are shown in blue whereas CG simulation results are in orange. The error bars of simulation results were estimated from block analysis.

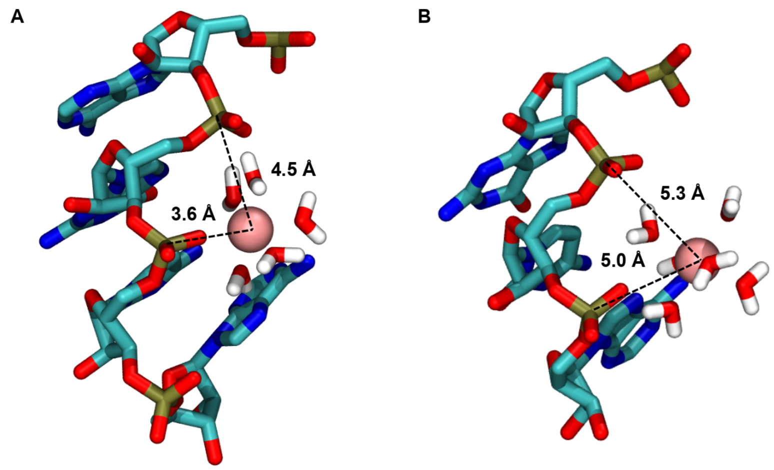


**Figure S12.** Different binding models of $Mg^{2+}$ with phosphate groups. (**A**) one inner-sphere binding with a distance of 3.6 Å and one outer-sphere binding with a distance of 4.5 Å. (**B**) Two outer-sphere binding with distances of 5.0 and 5.3 Å. A standard colour scheme is used for elements, and $Mg^{2+}$ is shown in a pink vdW sphere.

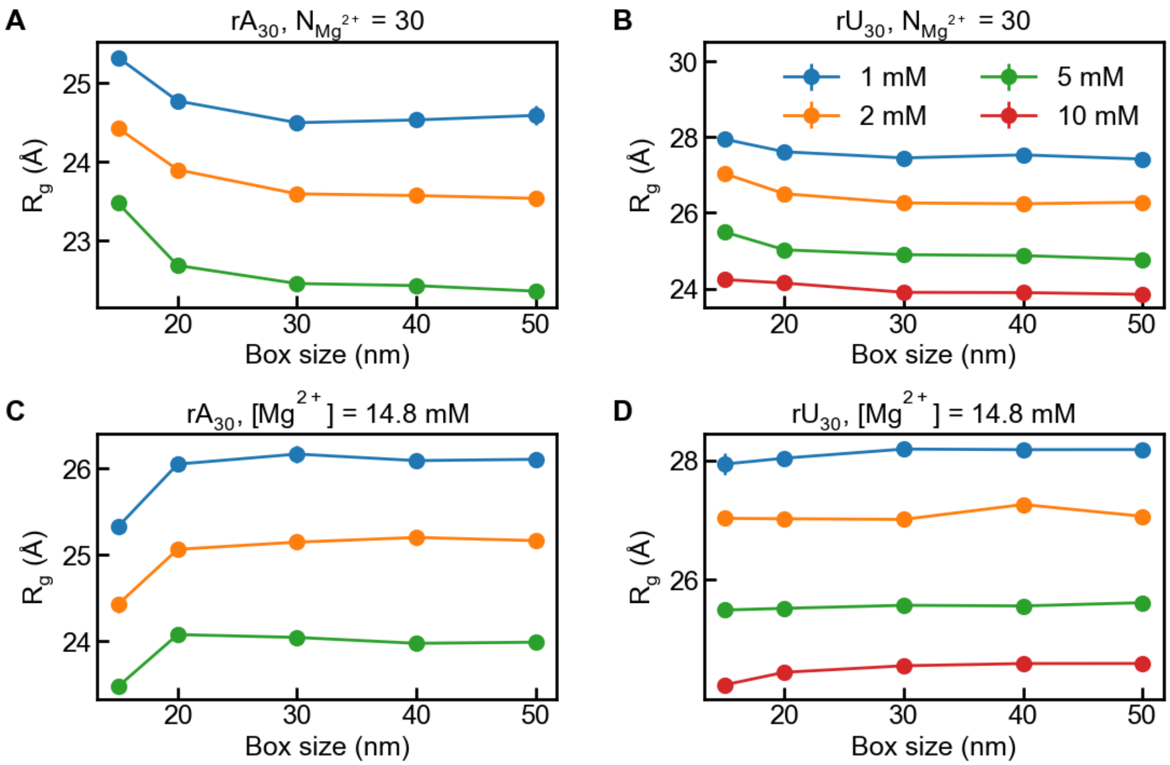


**Figure S13.** Effect of the simulation box size on $R_g$ with fixed number of $Mg^{2+}$ (**A, B**) and fixed concentration of $Mg^{2+}$ (**C, D**). Panels A and C are for $rA_{30}$, while panels B and D are for $rU_{30}$.

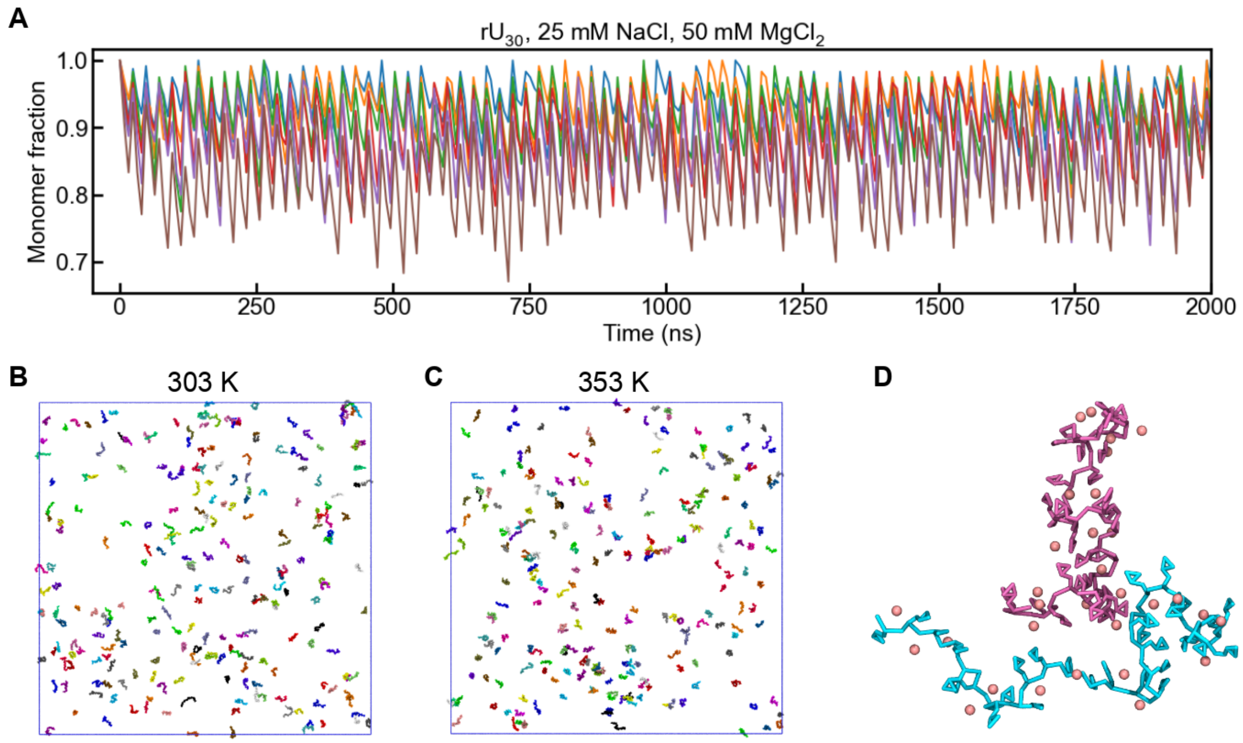


**Figure S14.** Phase separation simulations of $rU_{30}$. (**A**) Monomer fractions at 303, 313, 323, 333, 343, and 353 K under 25 mM NaCl and 50 mM $MgCl_2$. The initial RNA concentration is 50 μM. Note that the system remains dispersed at all temperatures. (**B, C**) Snapshots at 303 and 353 K. $Mg^{2+}$ ions were omitted for clarity. (**D**) A representative snapshot of $rU_{30}$ dimer, with $Mg^{2+}$ ions shown as pink beads.

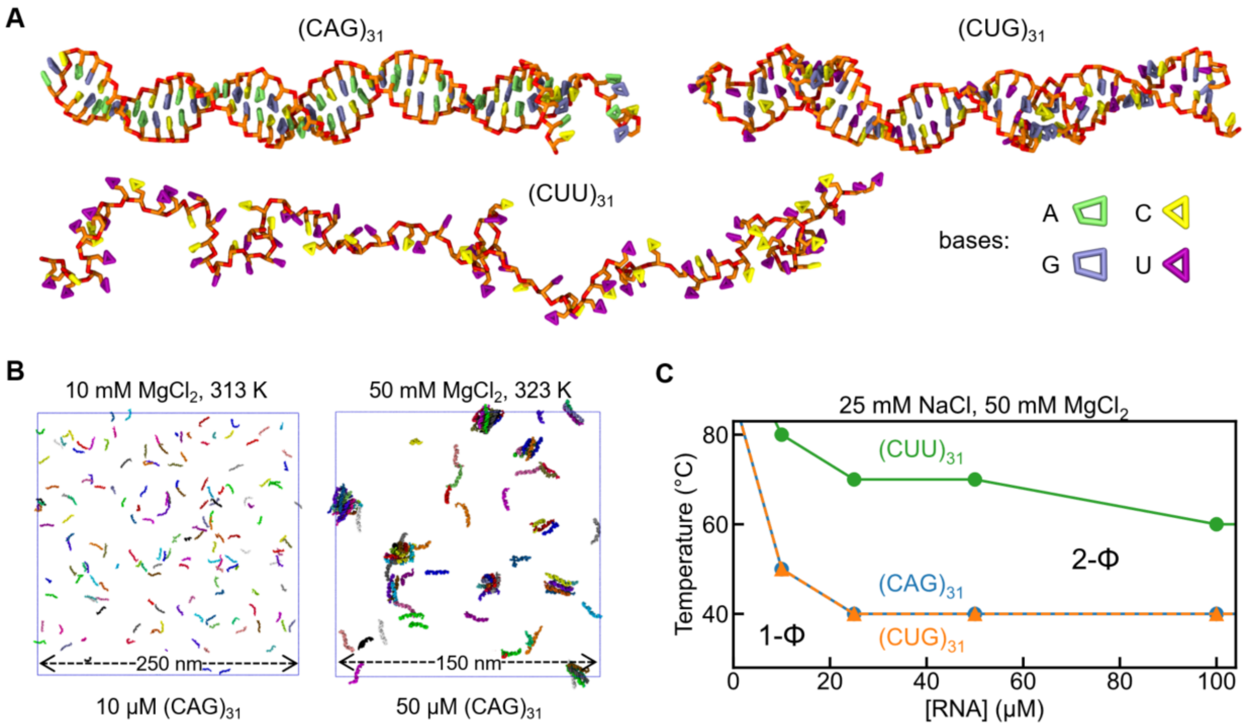


**Figure S15.** Phase separation of RNA triplet repeats. (**A**) Snapshots of representative structures of monomeric $(CAG)_{31}$, $(CUG)_{31}$, and $(CUU)_{31}$ under 25 mM NaCl at 303 K. Phosphate bead B1 and ribose beads B2/B3 are in red and orange, while bases A, G, C, and U are in lime, ice blue, yellow, and purple, respectively. (**B**) Snapshots of representative points in 1-phase (1-Φ, left panel) and 2-phase (2-Φ, right panel) regions, which were obtained from simulations of 10 μM $(CAG)_{31}$ at 313 K and 50 μM $(CAG)_{31}$ at 323 K, respectively (with 25 mM NaCl and 10 mM $MgCl_2$). RNAs are coloured in chains. Mg were omitted for clarity. (**C**) Phase diagrams of $(CAG)_{31}$ (blue), $(CUG)_{31}$ (orange), and $(CUU)_{31}$ (green) under 25 mM NaCl and 50 mM $MgCl_2$, where transition temperatures are plotted as a function of [RNA]. For each RNA, the lower left area of the boundary line is the 1-Φ region, while the upper right one is the 2-Φ region.

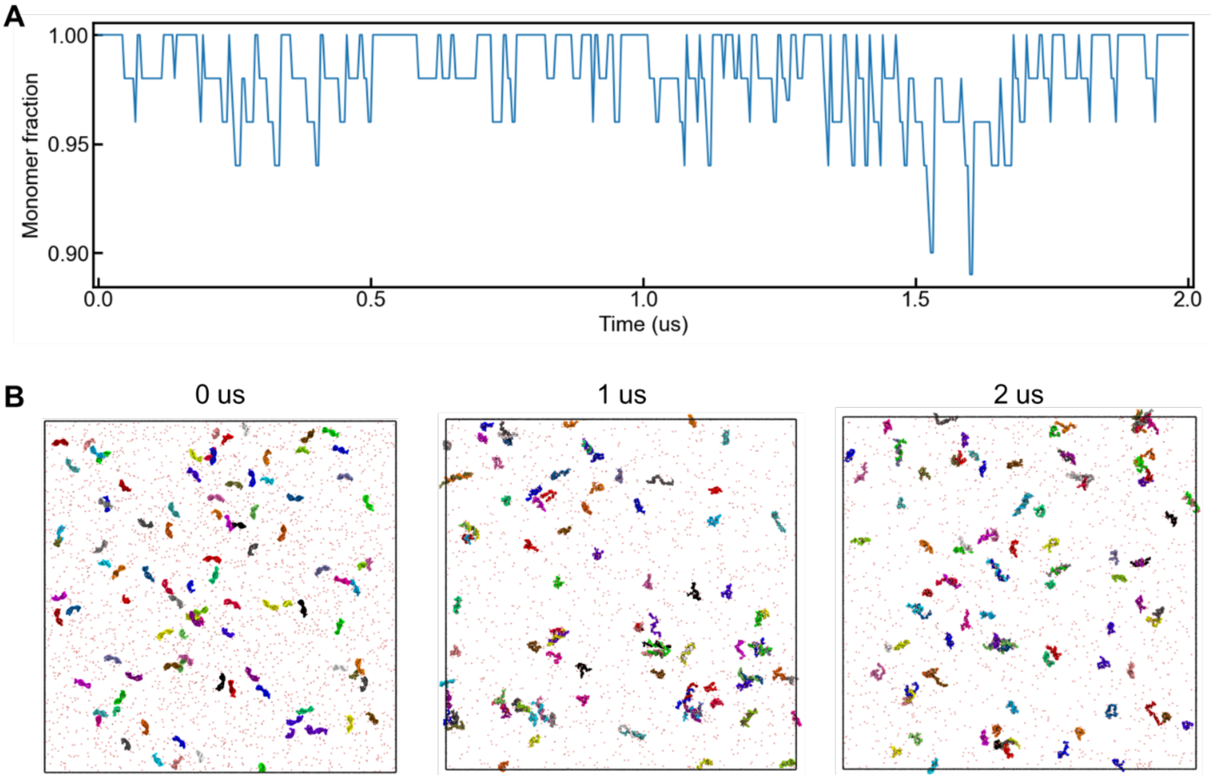


**Figure S16.** Phase separation simulations of 100 μM $(CAG)_{10}$ under 25 mM NaCl and 50 mM $MgCl_2$ at 353 K. (**A**) Monomer fraction as a function of simulation time. (**B**) Simulation snapshots at 0, 1, and 2 μs.

### Supplementary References

1. Plumridge, A., Andresen, K. and Pollack, L. (2020) Visualizing Disordered Single-Stranded RNA: Connecting Sequence, Structure, and Electrostatics. *J. Am. Chem. Soc.*, **142**, 109-119.
2. Theimer, C.A., Finger, L.D., Trantirek, L. and Feigon, J. (2003) Mutations linked to dyskeratosis congenita cause changes in the structural equilibrium in telomerase RNA. *Proc. Natl. Acad. Sci. U. S. A.*, **100**, 449-454.
